# Motif-based model of transcription predicts effects of sequence variants in AR enhancers and reveals distinct functions for AR-associated transcription factors

**DOI:** 10.64898/2026.09.02.748967

**Authors:** Hoda Taeb, Amin Safaeesirat, Emirhan Tekoglu, Kevin Xiao, Chia-Chi Flora Huang, Nathan A. Lack, Eldon Emberly

## Abstract

Androgen receptor (AR)-mediated transcription plays a central role in prostate cancer development and progression, yet the contributions of individual transcription factors (TFs) to AR-dependent enhancer activity remain incompletely understood. Here we use a biophysically motivated, interpretable motif-based model to dissect these contributions from STARR-seq data in LNCaP cells. By fitting the model separately to androgen inducibility and to baseline enhancer activity, we resolve TFs into three functional classes: hormone-dependent drivers, constitutive activators, and dual-role factors that contribute to both. These patterns suggest that inducibility is associated not only with the presence of AR and co-activator motifs, but also with the relative absence of constitutive activators that may saturate enhancer output. We validate the model against an independent saturation-mutagenesis dataset spanning 40 AR enhancers, predicting mutational effects at single-base resolution (AUC = 0.76), and show that direct fitting to these data independently recovers known AR regulators. Finally, we apply the model to prostate cancer GWAS risk alleles in AR binding site regions, prioritizing four candidate variants predicted to reduce the DHT/EtOH enhancer activity ratio at these loci.

## 1 Introduction

Transcription factors (TFs) regulate gene expression by controlling the recruitment and activity of RNA polymerase at target promoters [1, 2], governing critical cellular processes such as proliferation, differentiation, and apoptosis. Therefore, alterations in transcriptional regulation can shift gene expression programs in ways that promote tumor initiation and progression [3, 4]. In prostate cancer, the androgen receptor (AR), a nuclear hormone receptor activated by androgen signaling, acts as a key TF driving disease development and progression [5, 6]. AR-mediated transcription is shaped not only by AR itself but also by co-regulatory TFs including FOXA1, HOXB13, and GATA2, which influence AR binding and chromatin accessibility [7–14]. Dysregulation of these networks can drive tumor growth, metastasis, and resistance to therapy [14, 15].

Identifying functional TFs and characterizing their regulatory roles has increasingly relied on computational strategies informed by targeted experimental data. Methods such as ChIP-seq and ATAC-seq provide information on TF binding and chromatin accessibility, while Massively Parallel Reporter Assays (MPRAs) such as STARR-seq enable direct, high-throughput measurement of enhancer activity across thousands of sequence variants [16–18]. Early computational models leveraged biophysical principles, estimating TF–DNA binding affinity through thermodynamic frameworks that treat gene regulation as a function of molecular binding equilibria [19–23]. More recently, deep learning approaches have demonstrated strong predictive power in learning TF binding landscapes directly from sequence [24, 25], and models such as DeepSTARR have shown that enhancer activity measured by STARR-seq can be predicted directly from DNA sequence [26]. Building on these advances, large-scale genome language models (gLMs) treat DNA as biological text to capture complex, long-range regulatory interactions [27], though their inherent complexity limits biological interpretability [28]. While post-hoc methods such as DeepLIFT [29, 30] can extract regulatory insight from these models, complementary efforts have sought to develop biologically interpretable frameworks that directly link sequence features to quantitative regulatory activity [31, 32], the strategy we build upon here.

In the context of AR-mediated transcription, several TFs have already been identified as functional regulators [7–13]; however, many TFs remain uncharacterized, and the precise contributions of known TFs to these transcriptional programs are not fully understood. The availability of large-scale STARR-seq datasets in this system now creates an opportunity to apply interpretable sequence-based models to systematically infer TF contributions to AR-driven enhancer activity. Defining these roles is critical for advancing our understanding of prostate cancer biology and for developing more effective therapeutic strategies, particularly in advanced stages such as castration-resistant prostate cancer (CRPC) [33].

Here, we present a biophysically motivated, motif-based model of transcription, adapting the linear TF-occupancy framework of de Boer et al. [31], to systematically investigate TF contributions to AR-mediated transcription in LNCaP cells. The model is trained on STARR-seq data measuring enhancer activity across thousands of clinical AR binding sites [34]. It is separately fit to basal and hormone-induced enhancer activities as well as their relative changes, allowing us to quantify the effective interactions of specific TFs with RNA polymerase II and their contributions to AR-driven enhancer activity. Based on these contributions, TFs were grouped according to their inferred functional roles. We further validated the model against an independent saturation mutagenesis dataset, forecasting mutational effects at single-base resolution across 40 AR enhancers and attributing predicted activity changes to specific TF groups, providing mechanistic interpretability alongside predictive performance.

## 2 Methods

### 2.1 Data

#### 2.1.1 Quantifying the differential enhancer activity of androgen-responsive elements using STARR-seq

The enhancer activity of 365,530 DNA fragments spanning 7,422 genomic regions was measured in LNCaP cells under androgen-deprived (ethanol; EtOH) and androgen-containing (dihydrotestosterone; DHT) conditions using the STARR-seq assay [18, 34] (Fig. 1A-C). The genomic regions were selected to represent three categories: (i) 4,139 AR binding site (ARBS) regions with confirmed AR occupancy, (ii) 2,783 regions containing an androgen response element (ARE) motif but lacking AR binding, serving as negative controls, and (iii) 500 previously identified active LNCaP enhancers not associated with AR, acting as positive controls. Fragments ranged in length from 144 to 975 bp (mean = 528 bp, SD = 89 bp), with enhancer activity measured in three replicates per condition as RNA counts, and input DNA counts recorded to normalize for input DNA abundance. Fragment DNA sequences served as model inputs, with condition-specific enhancer activities and their log-fold-change (LogFC) between DHT and EtOH conditions as response variables.

**Figure 1.**
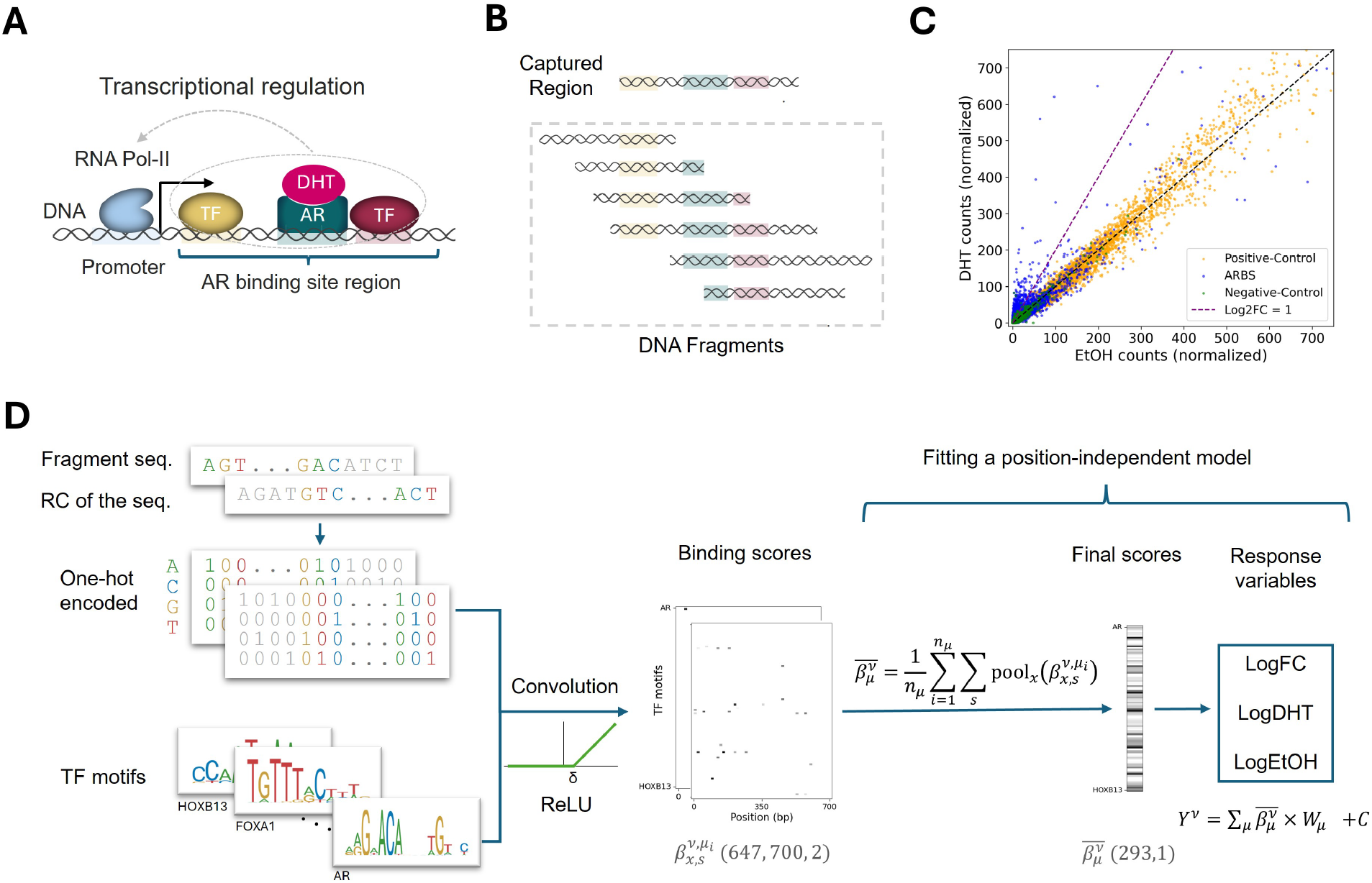
**A**, Transcriptional regulation at AR binding sites. The androgen receptor (AR, dark cyan) and co-regulatory TFs (yellow, red) bind to their respective binding sites (shown in matching colors on the DNA) and collectively regulate Pol II recruitment. The blue bracket indicates the androgen receptor binding site (ARBS) region. **B**, Fragmentation of enhancer regions. Colors on DNA fragments indicate TF binding sites. In the STARR-seq experiment, each enhancer region is fragmented into overlapping DNA segments (gray), which may contain all, part, or none of the TF binding sites. **C**, Normalized RNA counts under DHT versus EtOH conditions for DNA fragments overlapping ARBS (blue), positive-control (orange), and negative-control (green) regions. Counts were summed across three replicates, normalized by input DNA counts, and scaled for sequencing depth. The purple dashed line indicates Log2FC = 1; axis limits are set to 750. **D**, Model workflow. From left to right: (i) DNA fragments and their reverse complements (RC) are one-hot encoded after random padding. (ii) The energy matrix for each of the 647 TF motifs is convolved with the encoded sequences (Eq. 5), and an activation function (ReLU or Sigmoid) is applied (Eq. 6, 7) to generate the full matrix of binding scores 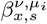. (iii) These scores are aggregated across motifs *µ*_*i*_, positions *x*, and strands *s* (Eq. 8) to yield the final binding score vector 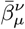. (iv) Finally, these scores are fitted to observed inducibility (LogFC) or enhancer activity (LogDHT, LogEtOH) using ridge regression to learn the TF weights *W*_*µ*_ (Eq. 9).

#### 2.1.2 Measuring effect of single-nucleotide substitutions on AR enhancers

Forty androgen-induced AR enhancers identified in Huang et al. [34] were targeted for saturation mutagenesis (Tekoglu et al., manuscript in preparation). The core 310 bp of each enhancer was cloned separately and inserted into the hSTARR ORI plasmid using error-prone PCR, which introduces random mutations throughout the insert. Simultaneously, random barcodes were introduced to each end of the insert, acting as unique molecular identifiers (UMIs) for matching of unique inserts to variants. Equal amounts of plasmid from each enhancer was pooled to generate the library. Enhancer activity for each variant and its corresponding wild-type (WT) sequence was quantified via STARR-seq under DHT treatment, with RNA and DNA read counts measured across three experimental replicates. Only variants carrying a single mutation were retained for downstream analysis. The generated raw data and count table is available on NCBI’s Gene Expression Omnibus (GEO) under the accession GSE335266.

### 2.2 Data preparation

#### 2.2.1 Enhancer activity data processing Response variables

For each DNA fragment *ν*, we define the normalized read count for condition *α* as:

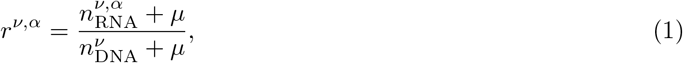

where 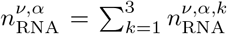 denotes RNA read counts summed across three replicates (*k*), and 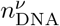 represents the corresponding input DNA counts. A pseudocount of *µ* = 5 is added to stabilize the ratio and avoid undefined values at low counts. This aggregation corresponds to the maximum-likelihood estimate of transcriptional activity under a Poisson sampling model.

We define three related response variables derived from the normalized read counts: condition-specific activities LogDHT = ln(*r*^*ν*,DHT^) and LogEtOH = ln(*r*^*ν*,EtOH^), as well as the log-fold-change LogFC = ln(*r*^*ν*,DHT^*/r*^*ν*,EtOH^), where ln denotes the natural logarithm. This logarithmic transformation stabilizes variance and compresses the dynamic range of expression levels. LogFC measures the relative change in enhancer activity between conditions, while LogDHT and LogEtOH measure absolute enhancer activity within each condition independently.

##### Input dataset construction and filtering

To obtain reliable activity estimates, we retain only fragments with 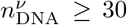. We further exclude fragments with LogFC *<* 0, keeping only androgen-activated or unresponsive fragments.

We select DNA fragments ranging from 400 to 700 bp and pad them to a uniform length of 700 bp. The padding consists of random sequences generated using nucleotide probabilities (*A* = 0.3, *C* = 0.2, *G* = 0.2, *T* = 0.3) to match the average GC content of the human genome [35] and our specific dataset. To ensure balanced representation across genomic regions, overlapping fragments are resampled to a uniform count of 14 per region, determined from the mean fragment coverage across regions. Finally, we exclude fragments overlapping the 40 AR enhancers used in the mutagenesis dataset from model training to prevent data leakage between training and evaluation sets. After applying these steps, the final dataset contains 99,610 DNA fragments for model training and evaluation.

#### 2.2.2 Mutagenesis library data processing

For each wild-type fragment and variant, we sum RNA counts across all unique plasmids and three replicates, and compute the input-normalized enhancer activity 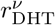 following Eq. 1, with the same pseudocount *µ* = 5.

We quantify the variant effect of each mutation on enhancer activity under DHT treatment as the log-fold-change in normalized activity relative to the corresponding wild-type sequence:

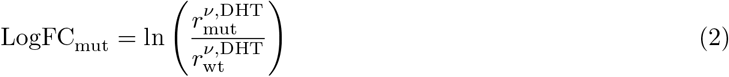

where 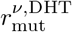 and 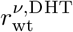 denote the normalized enhancer activity for the mutated and wild-type sequences, respectively.

To ensure data quality, we exclude variants observed in fewer than three unique plasmids and retain only those with 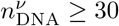, yielding a final dataset of 20,057 variants, corresponding to approximately 54% of all possible single-nucleotide substitutions across the 310-bp core sequences of the 40 AR enhancers. Despite this incomplete substitution-level coverage, at least one variant was observed at 77% of positions, indicating that the filtered dataset provides broad positional coverage sufficient for training and evaluating positional mutational effects across the 40 AR enhancers.

### 2.3 Transcription factor motif selection

We first identify protein-coding genes expressed in the LNCaP cell line using expression data from the Cancer Dependency Map (DepMap) portal (Public 25Q2 release) [36, 37], retaining only those with log_2_(TPM+1) *>* 3 to focus on robustly expressed genes. These genes are then intersected with human TFs cataloged in the JASPAR [38] and HOCOMOCO [39] databases, resulting in a final set of 293 TFs.

For these TFs, we construct a motif library using models from the JASPAR 2024 CORE *Homo sapiens* collection [38] and the HOCOMOCO v12 CORE set [39], yielding a combined library of 647 TF motif models. By including multiple motif variants for individual TFs, we improve the identification of potential TF binding sites across the analyzed genomic regions.

### 2.4 Motif-based biophysical model of transcription

We model enhancer activity across three related response variables: the differential enhancer activity (LogFC) and the condition-specific activities (LogDHT and LogEtOH). For each DNA fragment *ν*, we denote the observed output as *Y* ^*ν*^ and model it using a biophysically motivated and interpretable framework. This framework incorporates the binding scores of known TF motifs along the fragment’s sequence, following a strategy similar to [31], and predicts enhancer activity based on a linear combination of these scores. The computational pipeline, from one-hot encoding of padded STARR-seq fragments to the estimation of TF-specific weights linking binding to expression, is summarized in the workflow schematic (Fig. 1D).

Motifs for the 293 selected TFs are represented as Position Count Matrices (PCMs), where 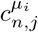 denotes the number of times nucleotide *n* is observed at position *j* of motif *µ*_*i*_. Here, *µ* denotes a TF and the index *i* distinguishes between different motif models associated with the same TF. Each PCM is converted to an energy matrix 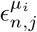 as:

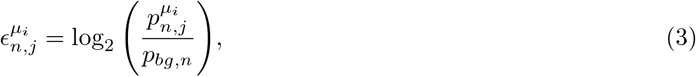

where

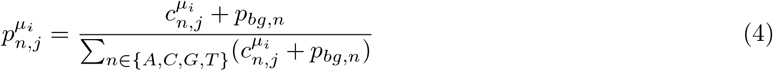

is the probability of finding nucleotide *n* at position *j* of motif *µ*_*i*_. The background probability *p*_*bg,n*_ is set to 0.3, 0.2, 0.2, and 0.3 for A, C, G, and T, respectively, matching the GC content of the human genome [35] and the nucleotide distribution of our processed dataset. These same background probabilities serve as pseudocounts to avoid zero probabilities at positions with limited observations [40]. The energy matrix 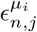 has dimensions 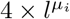, where 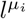 is the length of motif *µ*_*i*_.

For each TF motif *µ*_*i*_, we calculate its binding energy, 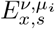, at each position *x* on each strand *s* of the DNA fragment *ν*. The motif is centered at position *x* using appropriate padding, and the binding energy is computed as:

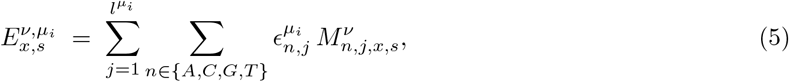

where 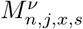 is the one-hot encoded representation of base *n* at motif position *j*. Eq. 5 follows the standard convolution of the motif with the DNA sequence [41].

Assuming independent binding of each TF to each position on a given strand, the probability of binding follows a logistic function of the binding energy and also depends on the TF’s chemical potential. To capture this non-linearity, we evaluate two candidate activation functions to compute the binding score 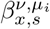 from the raw binding energies 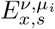: a thresholded Rectified Linear Unit (ReLU) and a shifted Sigmoid function, both parameterized by *δ*, analogous to the chemical potential, which sets the threshold for specific binding. The ReLU-based binding score is defined as:

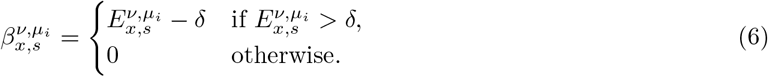

and the Sigmoid-based activation function is defined as:

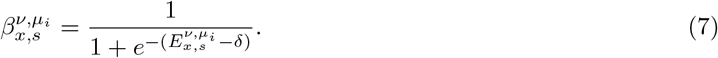

The final choice between these two activation functions and the optimal value for *δ* are determined through hyperparameter optimization, as detailed in Methods 2.5.

We relate the transcriptional output *Y* ^*ν*^ to TF binding scores using a linear model motivated by equilibrium thermodynamics of Pol II binding (see Section S1 for the biophysical motivation). To establish the input features for this model, we calculate an aggregated binding score, 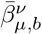, for each TF *µ* over a spatial unit *b*. This aggregated score is computed in two steps. First, binding scores 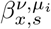 are pooled over the sequence positions *x* within unit *b* and summed over both strands *s*. Second, these values are aggregated across the *n*_*µ*_ motifs associated with that TF to account for both motif variants and potential model redundancies between the JASPAR and HOCOMOCO databases. We evaluate two motif-level aggregation strategies: averaging the scores across all available models or selecting the maximum score among them. All motifs for each TF are pre-aligned in the same orientation to ensure consistent strand assignment across motif variants. This procedure results in a single binding score per TF for each spatial unit:

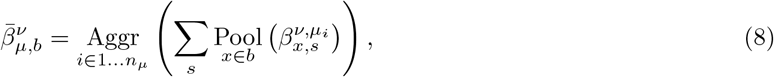

where the operator Pool(·) denotes either average or max pooling over the positions *x* within the unit *b*, and Aggr(·) represents the second aggregation step across motif models (mean or max).

We fit different models to these features to evaluate their performance and extract biological insights regarding TF contributions:

1. **Position-independent model**: The spatial unit *b* encompasses the entire 700-bp fragment (*b* = *L*), yielding one aggregated binding score per TF (293 features per fragment) for a position-independent linear regression [42]:

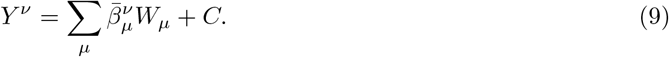

Here, *W*_*µ*_ represents the global contribution of TF *µ* to the response variable, and *C* is the bias term.
2. **Position-dependent model**: The spatial unit *b* corresponds to non-overlapping bins of length *l*_*b*_. For 700-bp fragments, we set *l*_*b*_ = 100 bp, resulting in seven discrete bins (*b* ∈ {1, …, 7}) and 2,051 features per fragment (293 × 7). The spatial regression model is expressed as:

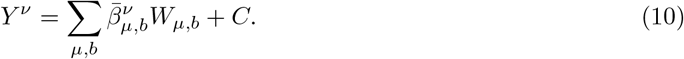

Here, the weights *W*_*µ,b*_ quantify the contribution of TF *µ* within bin *b* along the fragment sequence.

### 2.5 Model training

We train three separate linear models, each named after its response variable: the LogFC model to predict inducibility, the LogDHT model to predict enhancer activity under DHT conditions, and the LogEtOH model to predict enhancer activity under EtOH conditions. All three models are trained on the same processed dataset described in Methods 2.2 using ridge regression (*L*_2_ regularization) as implemented in the `scikit-learn` Python package [43].

For the position-independent model, given binding score vectors 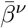 and observed target values *Y* ^*ν*^, where *Y* ^*ν*^ is one of LogFC, LogDHT, or LogEtOH, ridge regression estimates the weights *W*_*µ*_ and intercept *C* by minimizing the following objective function:

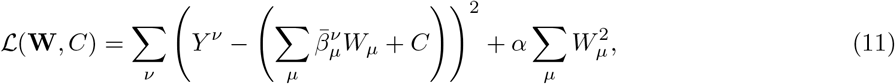

where the first term is the residual sum of squares and the second term penalizes large weights, with regularization strength *α >* 0. In this framework, the magnitude and sign of the fitted weights *W*_*µ*_ indicate the relative contribution of each TF to inducibility or enhancer activity, with positive weights identifying candidate drivers of the transcriptional response. The position-dependent model follows an identical formulation but includes the bin index *b* (as defined in Eq. 10) and is applied primarily to the LogFC target to investigate position-dependent TF contributions to androgen-induced activity. The hyperparameters for the ridge regression models include the spatial pooling method, motif aggregation approach, activation function, energy threshold *δ*, and regularization strength *α*, and are optimized via an exhaustive grid search. For model selection and evaluation, the dataset is partitioned by chromosome to ensure training and testing occur on independent genomic regions. Chromosomes 10 and 12 are reserved as a held-out test set, while the remaining 22 chromosomes (including X and Y) are used for hyperparameter tuning. Within the tuning set, we perform 5-fold cross-validation (CV), training on four folds and validating on the fifth in each iteration. To mitigate imbalances in fragment counts across folds, the random seed for fold assignment was chosen to yield a relatively balanced distribution of fragments. We evaluate each hyperparameter configuration based on the average mean-squared error (MSE) over the five validation folds and select the configuration that minimizes the average validation MSE. The grid search space is defined as follows:

- **Spatial Pooling (2):** Average or maximum pooling.
- **Motif Aggregation (2):** Average or maximum across TF motif models.
- **Activation Function (2):** ReLU or Sigmoid.
- **Energy Threshold** *δ* **(10):** Values from −2 to 16 in increments of 2.
- **Regularization** *α* **(16):** log_10_-spaced values from 10^−1^ to 10^6^ in steps of 0.5 dex, including *α* = 0 (no regularization).

This grid search yields 1,280 unique hyperparameter combinations, evaluated independently for the position-independent and position-dependent model architectures. Once hyperparameters are selected, we use 1,000 bootstrapped samples to report the mean and 95% confidence interval (CI) of the fitted weights.

### 2.6 Model validation on saturation mutagenesis data

We apply our fitted models to calculate the predicted effect of each point mutation on enhancer activity. For each variant and its corresponding wild-type fragment, the models predict 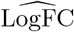 or 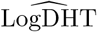 values (where the hat denotes model-predicted quantities). The predicted variant effect is calculated as the difference between the mutated and wild-type predictions. For the LogFC model, this is expressed as:

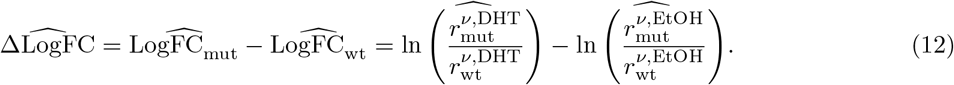

For the LogDHT model, the predicted variant effect is defined as:

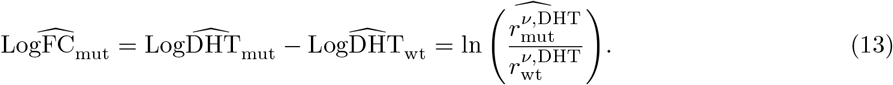

To quantify the overall mutational effect at each genomic position, the predicted values are averaged across all three variants at that base pair. For the LogFC and LogDHT models, these position-averaged predictions are denoted 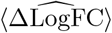 and 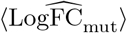, respectively. Similarly, the experimental mutational effect at each position, ⟨LogFC_mut_ ⟩, is calculated by averaging measurements over all available substitutions at that base pair.

To evaluate the model’s ability to distinguish functional from non-functional positions at single-base resolution, we classify genomic positions, using experimental and predicted effects averaged across the available substitutions at each base, into binary classes based on their experimental mutational effect ⟨LogFC_mut_⟩: 1 (functional) and 0 (non-functional). A position is assigned to class 1 if ⟨LogFC_mut_⟩ falls below a threshold *τ* for LoF effects or exceeds a threshold *τ* for GoF effects, and to class 0 otherwise. This procedure defines a ground truth for each threshold *τ*, allowing us to calculate the area under the curve (AUC) of the Receiver Operating Characteristic (ROC) for our model predictions. To assess how model performance varies with the stringency of the functional classification, we calculate the AUC across a range of *τ* values spanning the distribution of observed negative (for LoF) and positive (for GoF) ⟨LogFC_mut_⟩ values.

### 2.7 Direct model fitting to saturation mutagenesis data

We model the mutational effects on AR enhancer activity as a linear function of the change in aggregated TF binding scores upon mutation, 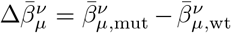, using the saturation mutagenesis dataset of 40 AR enhancers:

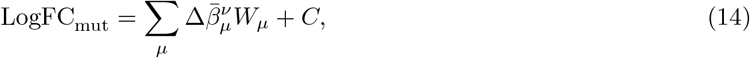

where 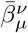 is aggregated over the entire 310-bp fragment following Eq. 8, and *C* is a constant intercept. We fit the model using ridge regression as implemented in `scikit-learn` [43]. The fitted weights *W*_*µ*_ represent the sensitivity of AR enhancer activity to perturbations in TF binding induced by single-nucleotide substitutions.

To ensure the model is evaluated on unseen genomic contexts, we partition the dataset by enhancer region rather than individual variants. Variants from 36 enhancers are used for model selection and 5-fold cross-validation, while variants from the remaining four enhancers are reserved as an independent test set. Hyperparameter optimization follows the grid search procedure described in Methods 2.5, with the regularization strength *α* varied from 10^−2^ to 10^4^ in increments of 0.5 dex. Final fitted weights and their corresponding 95% confidence intervals are estimated from 1,000 bootstrapped samples.

### 2.8 Direct prediction of GWAS variant effects

To assess whether prostate cancer risk variants may act through changes in enhancer activity, we applied the fitted LogFC model to single-nucleotide polymorphisms (SNPs) from the GWAS Catalog [44] (MONDO 0008315; prostate cancer and child traits) that overlap ARBS regions.

Starting from 4,374 GWAS Catalog entries, we lifted coordinates from GRCh38 to GRCh37 using the UCSC liftOver tool [45] and intersected them with the 4,139 ARBS regions used in model training. Each ARBS region spans a fixed 700-bp window centered on the AR ChIP-seq peak summit, consistent with the overlapping tiling design of the original STARR-seq assay [34]. This yielded 94 overlapping entries across 17 distinct ARBS regions, corresponding to 27 unique risk allele–region combinations at 19 genomic positions.

For each risk allele, we computed the predicted change in enhancer activity, 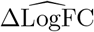, by substituting the variant into the corresponding 700-bp ARBS sequence and applying the fitted LogFC model, following the same procedure used for mutagenesis validation (Methods 2.6). Some GWAS Catalog entries report a risk allele as ambiguous, denoted by a question mark (“?”), meaning the specific risk-associated nucleotide at that position is not resolved in the catalog. For these entries, we computed predictions for all three possible non-reference alleles at that position and averaged them to obtain a single 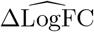 value.

To compare the magnitude of each predicted effect, we constructed a reference distribution from all possible non-reference single-nucleotide substitutions across the same 17 ARBS regions (*n* ≈ 35,700). GWAS variants with predicted effects falling outside the mean ± one standard deviation (SD) of this distribution were prioritized for further interpretation. To attribute the predicted effects to specific TFs, we decomposed each variant’s 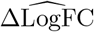 into individual TF contributions by leveraging the linearity of the model, as described for the mutagenesis analysis (Methods 2.6).

## 3 Results

### 3.1 Transcriptional models identify TF motifs contributing to androgen response and enhancer activity

We characterized the regulatory logic of androgen-responsive elements using STARR-seq data from 7,422 enhancer regions in LNCaP cells [34]. The dataset comprised 365,530 DNA fragments that tiled these enhancers under both androgen-containing (DHT) and androgen-deprived (EtOH) conditions (Fig. 1A, B). Because these fragments were inserted into a fixed plasmid context, the specific binding sites and potentially their spatial arrangement (i.e. grammar) were primary determinants of each fragment’s enhancer activity. As fragments tiled across an AR-bound enhancer, the AR binding site shifted positions, and neighboring motifs were captured or excluded depending on the fragment boundaries (Fig. 1B). This positional variation provided a rich substrate for identifying which factors and configurations drove differential output.

To decode this regulatory information, we developed a biophysically motivated model based on a mean-field, linearized thermodynamic framework (Fig. 1A; Section S1), in which each TF is assigned an effective weight representing its positive or negative influence on enhancer output (see Methods 2). We fit separate models to three response variables: inducibility (LogFC) and overall enhancer activity (LogDHT and LogEtOH). This let us distinguish TFs that specifically mediate androgen induction from those that establish baseline enhancer strength. Hyperparameter optimization via five-fold cross-validation yielded a positive optimal energy threshold for both inducibility and activity models (*δ* = 6 and *δ* = 8, respectively), effectively filtering out low-affinity interactions. Full cross-validation results and optimal hyperparameters are provided in Fig. S1 and Table S1, respectively.

The learned weights for the LogFC model are shown in Fig. 2A. TFs with positive weights are associated with higher inducibility (LogFC), with AR ranking highest, followed by ZBTB16, HOXB13, and FOXA1. For the LogDHT model, the top 40 TFs with the strongest positive and negative weights are shown in Fig. 2B, with the complete ranking provided in Fig. S2A. LogEtOH model weights are presented in Fig. S2B. Among the activity models, TP53, ZBTB45, and IRF7 emerge as the top contributors to overall enhancer activity. The LogDHT and LogEtOH models exhibit high consistency, sharing 172 TFs with common signs (Fig. S2C). A minority of TFs show condition-specific weights, such as ATF6 and ELF2, though their weight magnitudes are small relative to top contributors (Fig. S2D).

**Figure 2.**
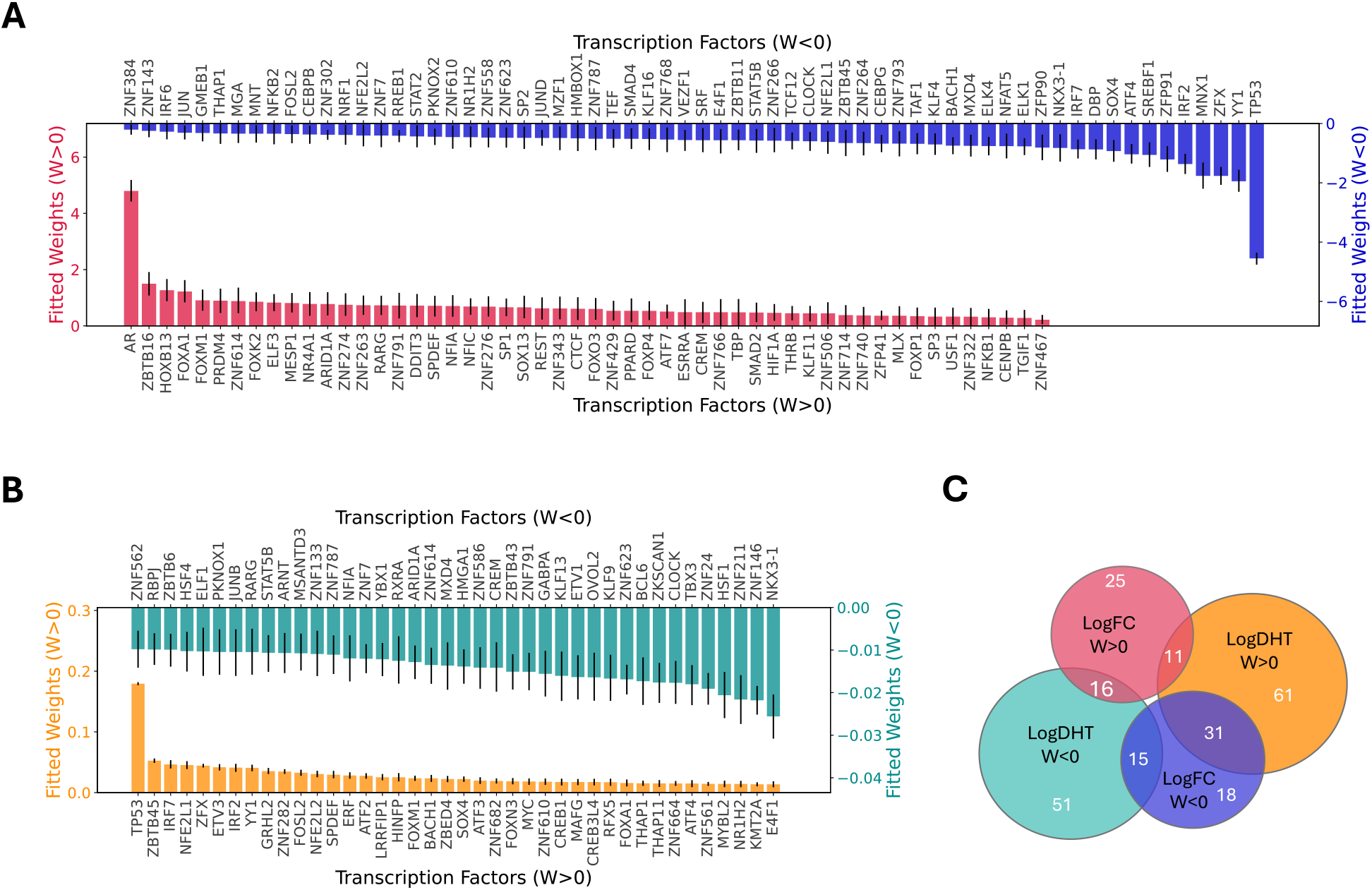
Distribution and stability of fitted TF weights across regulatory models. **A**, Ranked weights for the LogFC model, representing the contribution of TFs to inducibility. Bar heights indicate the mean weight 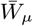 across 1,000 bootstrap samples; only TFs with a 95% confidence interval (CI) that does not cross zero are displayed. Error bars denote the 95% CI. **B**, Ranked weights for the LogDHT model, representing TF contributions to absolute enhancer activity. For clarity, only the 40 most positive and 40 most negative significant weights are displayed, with bootstrap statistics calculated as in **A. C**, Venn diagrams illustrating the overlap of TFs with significant positive (*W >* 0) or negative (*W <* 0) weights across the inducibility (LogFC) and activity models (LogDHT or LogEtOH), where significance is defined by a 95% CI that does not cross zero. Numbers denote the TF count within each unique or overlapping set.

Given the fitted weights of the 293 TFs in each model, we grouped them based on their contributions to inducibility and overall enhancer activity. The Venn diagram in Fig. 2C illustrates the subsets of TFs with positive and negative weights across models (full TF assignments provided in Table S2). A TF may enhance inducibility while simultaneously being associated with lower overall enhancer activity, as seen in the overlapping TFs between the LogFC *W >* 0 and LogDHT *W <* 0 sets. This is specifically the case for AR, which exerts the largest influence on inducibility despite being predictive of low enhancer activity. This pattern aligns with experimental observations that inducible DNA fragments often maintain low baseline activity rather than driving high constitutive expression. This pattern is illustrated in Fig. 1C: ARBS fragments cluster at the lower end of the activity spectrum yet are prominently positioned above the log_2_ FC = 1 threshold (purple dashed line), identifying them as primary inducible elements. Consistent with this low baseline, AR also carries a negative weight in the LogEtOH model (Fig. S2B); because AR is not activated under androgen-deprived conditions, this may reflect other factors binding the AR motif to keep transcription low when androgen is absent.

TFs at the intersection of LogFC *W >* 0 and LogDHT *W >* 0 sets are predictive of both androgen-induced and baseline enhancer activity, suggesting dual functional roles. These TFs may promote Pol II recruitment and cooperate with AR to stabilize its DNA binding. FOXA1 and SPDEF belong to this group, exhibiting substantial positive weights in both inducibility and activity models, consistent with their established roles as AR co-activators in prostate cancer [8, 46]. Notably, since STARR-seq operates in a plasmid context lacking canonical chromatin structure, the contribution of FOXA1 and other pioneer factors observed here likely reflects activity largely independent of their chromatin remodeling function [47], suggesting they play additional roles in AR-mediated transcription beyond establishing chromatin accessibility.

Another informative group is the intersection of LogFC *W <* 0 and LogDHT *W >* 0 sets, representing TFs with negative inducibility weights despite positive activity weights. TP53, carrying the largest negative inducibility weight yet the strongest positive activity weight, is the most prominent example. Other factors in this category include ATF4, SOX4, and IRF2 (Fig. 2C). The positive LogDHT and negative LogFC weights for these TFs suggest that fragments with high scores for these motifs tend to be highly active under both conditions yet poorly induced by androgen, consistent with a potential ceiling effect in which sequences with high constitutive enhancer activity have limited dynamic range for further induction by androgen. For TP53, this pattern could reflect transcription driven by the DNA damage response to the plasmid-based assay rather than by androgen signaling [48].

### 3.2 The LogFC model predicts mutational effects in AR enhancers

We next validated our fitted LogFC and LogDHT models by quantifying how well they predicted mutational effects at single-base resolution across AR enhancers. We compared model predictions directly with experimental measurements of enhancer activity differences between wild-type and mutated sequences (Fig. 3A). The experimental profile (top panel) shows the average change in activity, ⟨LogFC_mut_⟩, across a representative AR enhancer, with several prominent peaks corresponding to loss-of-function (LoF) mutations, while gain-of-function (GoF) effects are weaker and less frequent. This imbalance suggests these enhancer sequences are already highly optimized for androgen-induced activity, making random single-nucleotide substitutions far more likely to disrupt than to improve function.

**Figure 3.**
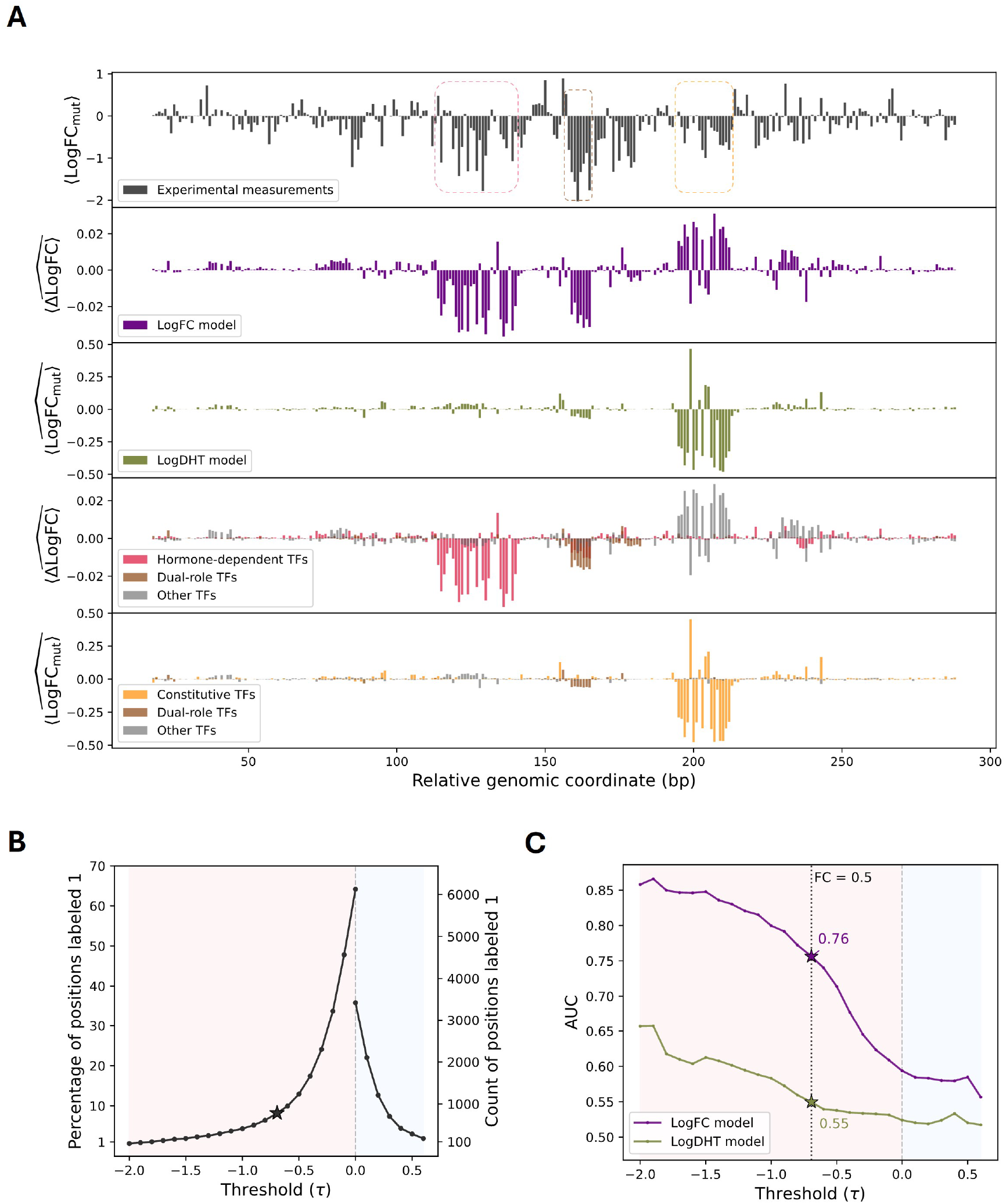
Performance of motif-based models in predicting mutational impacts. **A**, The top panel shows the experimental mutational profile of a representative AR enhancer, illustrating the average change in enhancer activity (⟨LogFC_mut_⟩) at each position, selected to illustrate contributions from all three TF groups including constitutive factors. The second and third panels display predictions generated by the LogFC and LogDHT models, respectively. The bottom panels show the linear decomposition of these predictions into contributions from three functional TF groups: hormone-dependent (red), dual-role (brown), and constitutive (orange). **B**, Classification of positions as functional (class 1) or non-functional (class 0) based on thresholds *τ* applied to experimental measurements across 40 AR enhancers. Data points show the absolute number of functional positions (left axis) and their percentage of the total position pool (right axis). Red and blue shaded regions indicate LoF and GoF thresholds, respectively; the starred threshold (*τ* = ln(0.5) ≈ −0.69) corresponds to a two-fold reduction in enhancer activity, at which 777 positions are classified as functional. **C**, AUC of the LogFC (purple) and LogDHT (olive green) models in classifying functional positions across thresholds *τ*. Shaded regions distinguish performance for LoF (red) and GoF (blue) positions.

For androgen-induced AR enhancers, transcriptional activity under the EtOH condition is typically minimal; consequently, single-nucleotide substitutions are expected to have negligible impact on EtOH activity, and the second term of 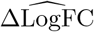 approaches zero. Thus, LogFC model predictions for these enhancers approximate the predicted change in DHT activity. Although both the LogFC and LogDHT models therefore predict changes in DHT activity, they were trained on different response variables and learned distinct TF weights, so their predictions capture complementary sets of variant effects. The second and third panels of Fig. 3A display the predictions generated by the LogFC and LogDHT models, respectively. The LogFC model captures the most prominent LoF peaks, while the LogDHT model predicts additional, lower-magnitude variations not captured by the LogFC model. To interpret which TF groups contribute to these predictions, we leveraged the linearity of our model to decompose the output into contributions from specific TF functional groups identified in our weight analysis (Fig. 2C). We categorized the contributing TFs into three groups based on their weights: hormone-dependent TFs (red; unique to LogFC *W >* 0), constitutive TFs (orange; unique to LogDHT *W >* 0), and dual-role TFs (brown; LogFC *W >* 0 ∩ LogDHT *W >* 0). As illustrated in the fourth subplot of Fig. 3A, this decomposition reveals which TF groups contribute to the observed LoF effects: peaks captured by the LogFC model primarily result from disruption of hormone-dependent TFs (red), such as AR or its coactivators, whereas peaks better explained by the LogDHT model arise from loss of constitutive factor binding that affects baseline enhancer activity (orange). Additionally, peaks in the central region are largely predicted by dual-role TFs (brown), which contribute to both inducibility and baseline activity, though some hormone-dependent TF contributions (red) are also present. Similar mutational profiles for all 40 AR enhancers are provided as Supplementary Data.

To quantitatively evaluate predictive performance across all 40 enhancer regions, we classified genomic positions as functional or non-functional based on their averaged experimental mutational effect and a sliding threshold *τ* (Methods 2.6). As shown in Fig. 3B, the number of positions labeled as functional narrows as *τ* increases, isolating positions that drive the most substantial changes in enhancer activity. The discriminative power of the two models across these thresholds is summarized in Fig. 3C. The LogFC model consistently achieves superior performance in separating functional from non-functional positions, with its AUC increasing as the classification criteria become more stringent. For instance, at the threshold where mutations reduce enhancer activity by half (*τ* = ln(0.5) ≈ −0.69), the LogFC model reaches an AUC of 0.76.

The LogFC model substantially outperforms the LogDHT model across these regions. This difference likely reflects two factors: first, because the 40 enhancers were selected for strong androgen inducibility, their activity is predominantly androgen-driven; functional mutations therefore disrupt hormone-dependent TF binding (such as AR) rather than constitutive factor binding; second, the LogDHT model was trained to predict absolute enhancer activity across a broad set of regions, and is therefore optimized for high-output enhancers; its fitted weights may not transfer well to these lower-activity, AR-specific regions. Together, these results validate our framework and demonstrate that our motif-based approach can identify genomic positions that are sensitive to sequence perturbation.

### 3.3 Direct model fitting to mutagenesis data confirms key AR-associated regulators

To gain deeper insight into the factors regulating AR-mediated transcription, we trained a complementary model using the mutagenesis dataset of 40 AR enhancers (Tekoglu et al., manuscript in preparation), which provides enhancer activity measurements for both wild-type sequences and their single-nucleotide substitutions (Methods 2.1.2). The model predicts how individual variants alter enhancer activity under DHT treatment by relating changes in TF binding scores 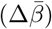 to the observed change in activity LogFC_mut_. Hyperparameter optimization yielded an energy threshold *δ* = 4 and regularization strength *α* = 10^−1.5^; grid search results and optimal hyperparameters are provided in Fig. S3 and Table S1, respectively.

The fitted weights of this model are shown in Fig. 4A. The top TFs with positive weights are AR, FOXA1, and HOXB13, consistent with their established roles in AR-mediated transcription [8]. FOXM1 also receives a positive weight, potentially due to motif similarity with FOXA1. Although these 40 regions represent only about 10% of inducible AR enhancers in LNCaP cells, they were selected to span a wide range of enhancer activity and are therefore representative of the broader inducible population. The single-nucleotide resolution of this dataset offers valuable insight into which TF motifs are associated with increased or decreased enhancer activity under DHT conditions, as inferred from the sign and magnitude of the fitted weights.

**Figure 4.**
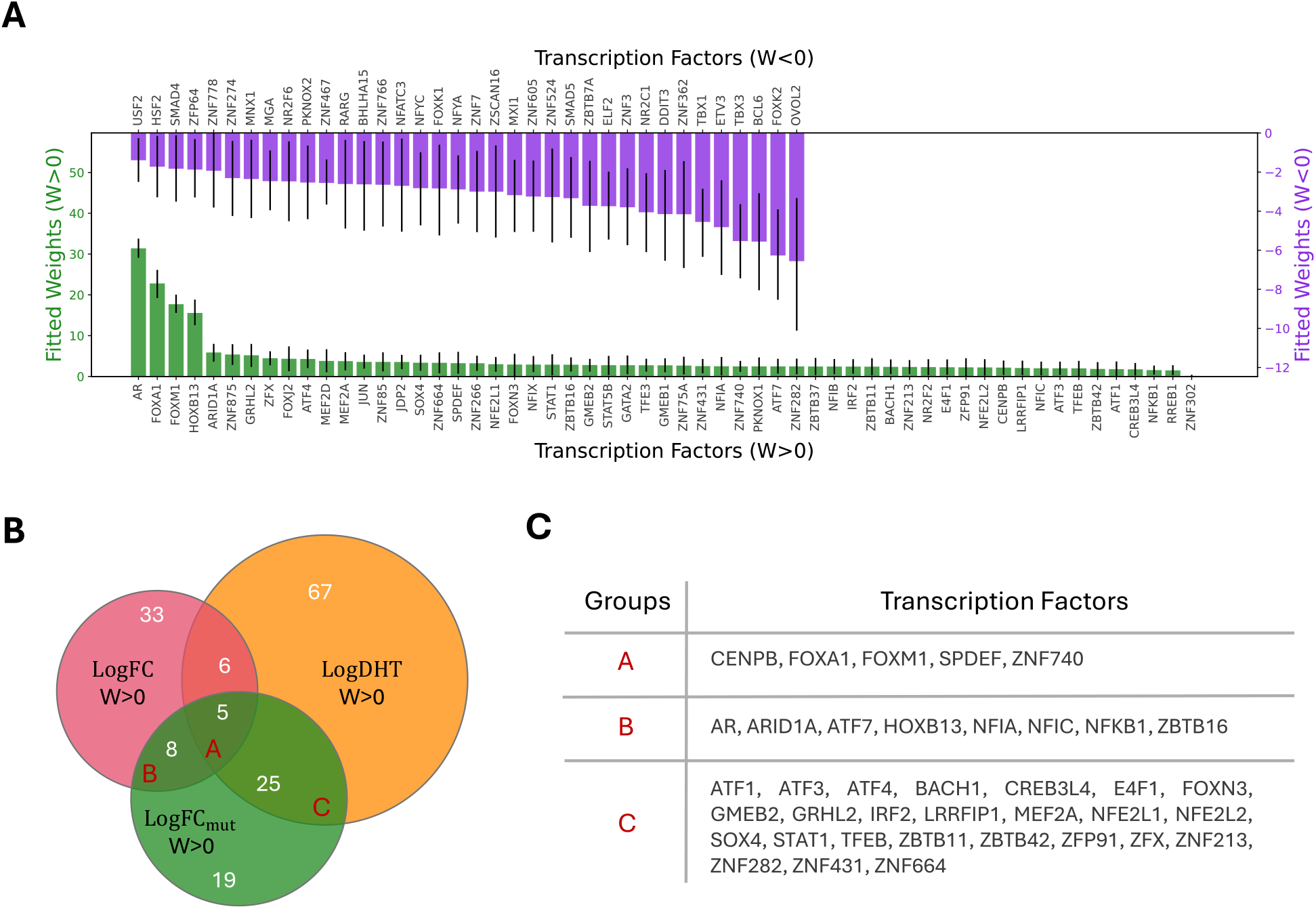
TF weights from direct mutagenesis fitting. **A**, Fitted TF weights from the mutagenesis model. Green bars indicate positive weights (*W >* 0), and purple bars indicate negative weights (*W <* 0), ranked by magnitude. Error bars represent the 95% CI from 1,000 bootstrap samples; only TFs whose CI does not cross zero are displayed. **B**, Venn diagram illustrating the overlap of TFs with positive weights across the LogFC, LogDHT, and mutagenesis models. **C**, Summary table of TFs with positive weights in the mutagenesis model that overlap with the LogFC model, LogDHT model, or both.

The Venn diagram in Fig. 4B shows overlapping TFs among the LogFC, LogDHT, and mutagenesis models, with the table in Fig. 4C listing TFs within each overlapping set. Because only a limited set of TF binding sites is represented in these 40 enhancers, we do not expect to observe all functionally important TFs identified by the broader models. Nevertheless, TFs that appear consistently with significant positive or negative weights across all three models represent the most robust regulatory factors, as their regulatory roles are consistently identified across multiple independent datasets and modeling approaches.

### 3.4 Spatial organization of TF sites in AR inducible enhancers plays a minor role in inducibility

In the STARR-seq assay, tested DNA fragments are positioned downstream of a promoter, where their regulatory activity can modulate transcriptional output (Fig. 5A). Because TFs can bind at different positions along these fragments, and consequently at different distances from the reporter transcription start site (TSS), we investigated whether TF binding position influenced androgen inducibility using the position-dependent model (Methods 2.4). To balance spatial resolution with statistical power, we discretized the 700-bp sequences into seven 100-bp bins, allowing the model to learn position-dependent weights that quantify the functional contribution of each TF across the sequence.

**Figure 5.**
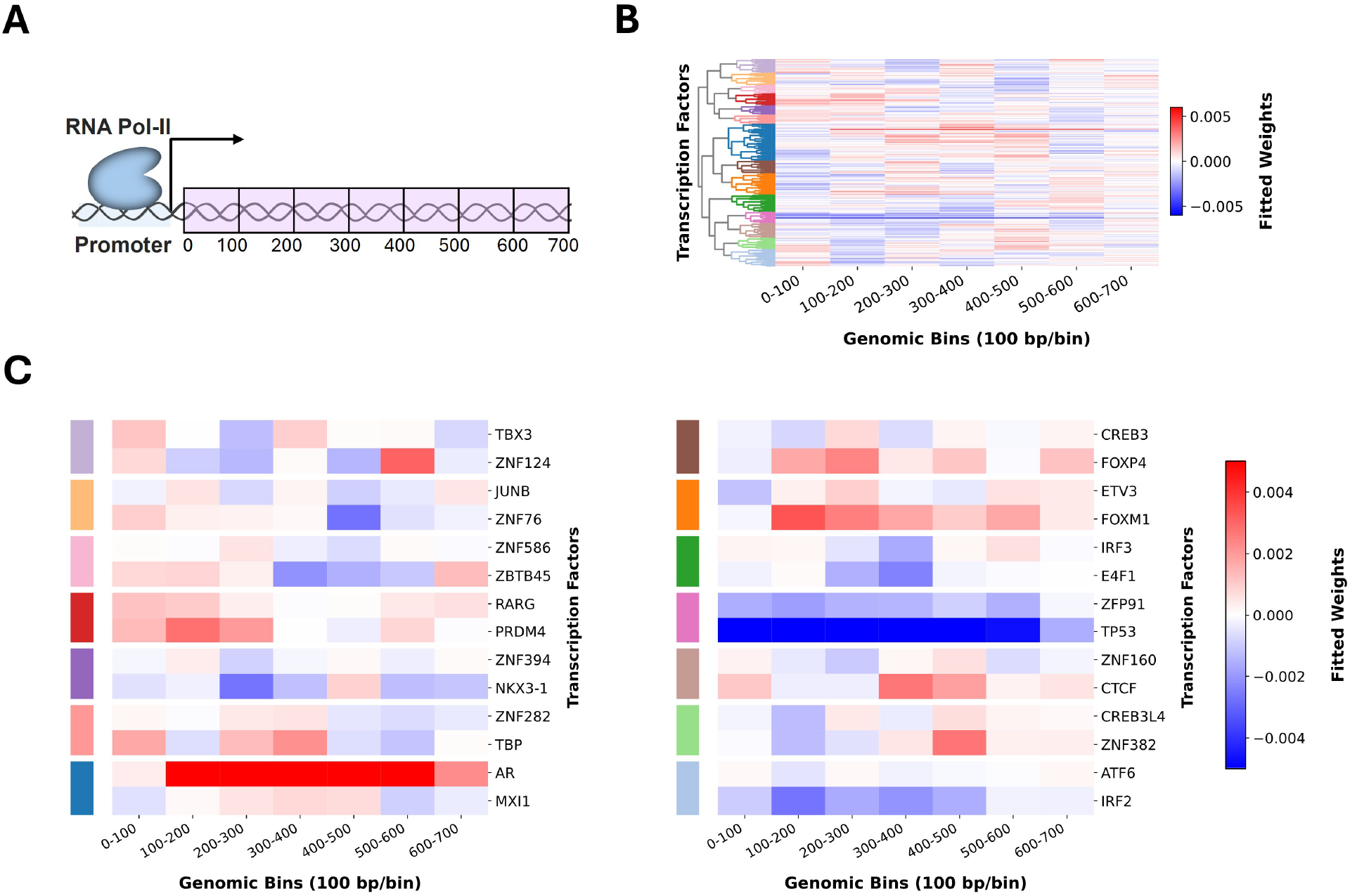
Spatial binding preferences of 293 transcription factors. **A**, Schematic illustrating the position of the promoter relative to the tested DNA fragment and its spatial bins. **B**, Heatmap of fitted weights from the position-dependent model, ordered by hierarchical clustering using correlation distance and average linkage. Fourteen distinct clusters were identified at a correlation threshold of 0.3, denoted by the color strip on the left. **C**, Representative TF weight profiles for the 14 clusters. Two TFs per cluster are shown: the top TF represents the member with minimum Euclidean distance to the cluster centroid (most typical example), while the bottom TF was selected based on the highest absolute weight across the seven spatial bins (strongest regulatory signal). Horizontal white bars separate the 14 clusters; the color strip (left) indicates cluster membership, consistent with panel B.

Despite the potential for positional effects, incorporating spatial information did not improve LogFC prediction or AUC compared to the position-independent model; furthermore, the average weights across bins remained highly correlated with those of the position-independent model (Fig. S4). However, the fitted weights revealed distinct spatial binding patterns among different TFs (Fig. 5B). To systematically characterize this diversity, we performed hierarchical clustering on the fitted weight profiles of 293 TFs using a correlation-based distance metric, identifying 14 distinct clusters by cutting the dendrogram at a correlation distance of 0.3 (Fig. 5B). These clusters define a spectrum of spatial preferences along the fragment, corresponding to different distances from the reporter TSS.

Two representative TFs from each cluster were selected to illustrate these spatial patterns (Fig. 5C). AR, in the dark blue cluster, exhibited a uniform positive pattern with consistent weights across all bins, indicating it contributes to inducibility regardless of position along the fragment, though other TFs in this cluster showed predominantly central activity. In contrast, most TFs across other clusters displayed position-specific patterns. The red cluster showed stronger positive weights in the proximal portion of the fragment, within the first 300 bp, while the dark pink cluster exhibited either uniform negative weights (as seen in TP53, consistent with its role in the position-independent model) or distal activity with proximal depletion. A similarly position-invariant profile for TP53 was observed in the LogDHT and LogEtOH position-dependent models (Fig. S5). Collectively, these results suggest that while spatial positioning is not a dominant driver of global model performance, many individual TFs exhibit characteristic position-dependent associations with androgen inducibility, whereas AR and TP53 show comparatively position-invariant profiles.

### 3.5 The model prioritizes four prostate cancer GWAS risk alleles predicted to reduce ARBS LogFC

Prostate cancer risk loci identified by genome-wide association studies (GWAS) frequently reside in non-coding regulatory regions, raising the question of whether they act, at least in part, through changes in enhancer activity. To investigate this, we applied our LogFC model, validated against saturation mutagenesis data (Section 3.2), directly to prostate cancer risk SNPs overlapping ARBS regions, without any additional training (Methods 2.8).

Of the 27 GWAS risk allele–region combinations tested, four fell below the 1-SD threshold, all with predicted decreases in LogFC (Fig. 6A). The largest predicted decrease was at rs72848121-G, followed by rs11986220-A, rs1160267-G, and rs11665748-A, with replication support ranging from 2 to 10 independent GWAS studies (Table S3). Of these, only rs11665748-A, located near the KLK3 promoter [49], falls within a region showing strong androgen induction in our STARR-seq data (region LogFC = 3.4); at this locus, a predicted decrease would specifically reflect reduced androgen-driven inducibility. The remaining three loci lie in regions with low or negative measured induction, where the model’s predicted decrease reflects a further shift in the predicted DHT/EtOH ratio rather than the loss of an established androgen response.

**Figure 6.**
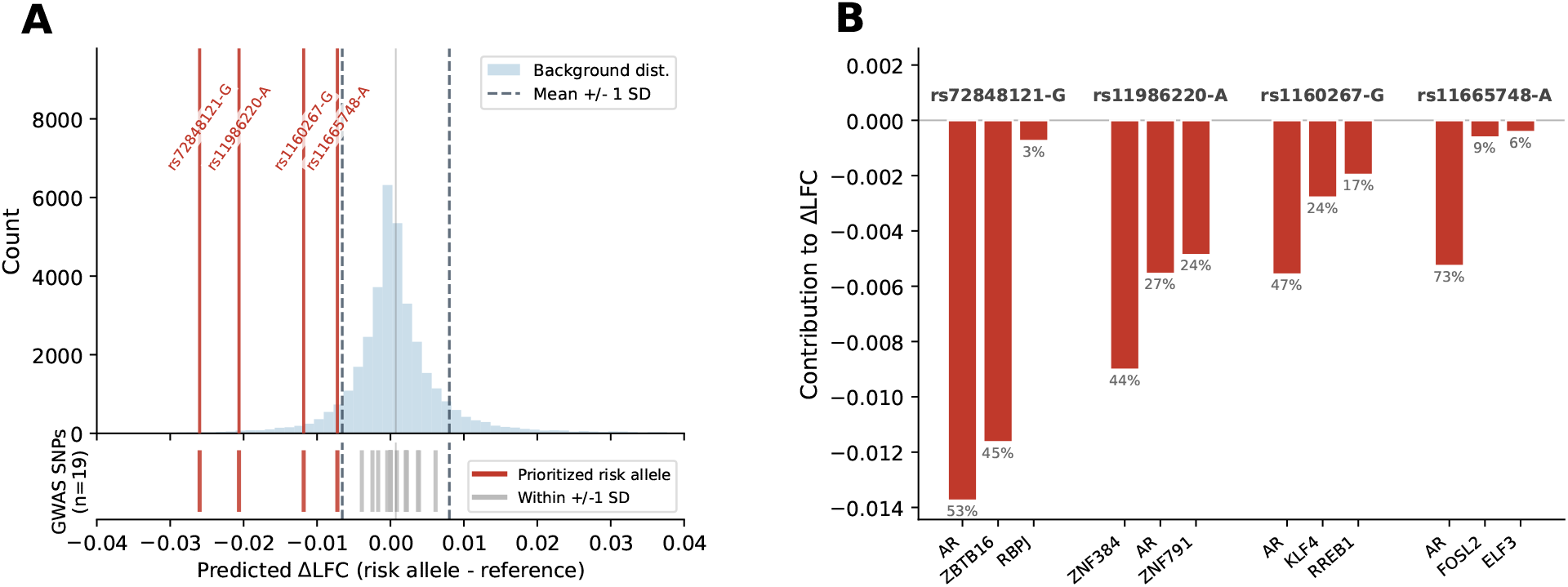
Predicted effects of prostate cancer GWAS risk alleles on inducibility of ARBS regions. **A**, Histogram shows the background distribution of predicted 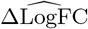 values for all possible non-reference single-nucleotide substitutions across 17 ARBS regions (*n* = 35,700 substitutions). Dashed vertical lines indicate the background mean ± 1 SD. Colored tick marks in the lower panel show the predicted ΔLogFC for each of the 27 unique GWAS risk allele–region combinations (GWAS Catalog, MONDO 0008315; prostate cancer and child traits). The four risk alleles with predicted ΔLogFC exceeding 1 SD below the background mean are highlighted in red and labeled with their rsID. **B**, Linear decomposition of the predicted ΔLogFC for each prioritized risk allele into individual transcription-factor contributions (ridge-model weights × change in motif binding score). Bars show the three TFs with the most negative contributions at the risk-allele position; percentages indicate each TF’s share of the total predicted effect.

To identify which TFs drive these predicted effects, we decomposed each variant’s 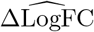 into individual TF contributions (Fig. 6B), as in Fig. 3A. For the rs11665748-A variant, AR is the dominant contributor, accounting for 73% of the predicted reduction in inducibility, consistent with this variant disrupting a functional androgen response element. Taken together, these results illustrate how the LogFC model can prioritize candidate regulatory GWAS variants and provide hypotheses about the motif-associated mechanisms underlying their predicted effects, without requiring variant-specific experimental data.

## 4 Discussion

AR-mediated transcription plays a central role in prostate cancer initiation and progression. Understanding its co-regulatory network is essential for uncovering the mechanisms that govern its control. In this study, we developed a motif-based model of transcription, inspired by biophysical modeling, to quantify the contributions of TFs to androgen-induced enhancer activity using STARR-seq data from LNCaP cells. Our framework provides a direct and interpretable link between DNA sequence and enhancer activity, highlighting regulators including well-characterized factors such as AR, FOXA1, and HOXB13, as well as less-studied co-regulators that have not been previously emphasized in the context of AR biology.

By fitting separate models to inducibility (LogFC) and overall enhancer activity (LogDHT and LogEtOH), we obtained a complementary view of TF function along two dimensions: driving androgen responsiveness and establishing baseline transcriptional output. We identified three major TF classes: hormone-dependent drivers (e.g., AR, HOXB13), constitutive activators (e.g., TP53), and dual-role factors (e.g., FOXA1, SPDEF). Notably, constitutive activators with negative inducibility weights but positive activity weights, such as TP53, illustrate that maximal androgen inducibility is shaped not only by the presence of AR and co-activator motifs, but also by the relative absence of these strong constitutive drivers that may saturate enhancer output and limit the dynamic range of the androgen response. However, the inferred role of TP53 warrants careful interpretation, as its condition-independent activation pattern may partly reflect a DNA damage response to the plasmid DNA inherent to the STARR-seq assay rather than a direct regulatory role in AR-mediated transcription.

To validate the inducibility model, we tested its predictions against an independent saturation mutagenesis dataset, demonstrating its ability to predict mutational effects at single-base resolution across 40 AR enhancers (AUC = 0.76 at the two-fold activity-reduction threshold). Training directly on the mutagenesis data provided an independent confirmation of key regulators: AR, FOXA1, and HOXB13 emerged with the strongest positive weights, consistent with their established roles in AR-mediated transcription. Comparing these findings with the STARR-seq models enabled the identification of a high-confidence subset of functional TFs.

Beyond validation on experimentally mutagenized sequences, we further demonstrated that the LogFC model can be applied directly to naturally occurring genetic variation. By intersecting prostate cancer GWAS risk alleles with ARBS regions, we prioritized four variants with predicted decreases in LogFC. Of these, rs11665748-A is particularly interpretable because it lies within a region showing strong experimentally measured androgen induction, and AR accounts for the majority of its predicted effect, consistent with reduced androgen responsiveness. The remaining three variants occur in regions with weak or negative measured induction, and their predicted effects therefore represent shifts in the DHT/EtOH activity ratio rather than disruption of an established androgen response. More broadly, this application illustrates how an interpretable, motif-based model can link non-coding disease risk variants to a candidate regulatory mechanism, without requiring variant-specific experimental data as input.

Incorporating position-dependent features suggests that predicted TF binding at specific locations can have distinct associations with enhancer activity. While spatial information had a limited effect on overall predictive performance, hierarchical clustering of position-dependent weights identified 14 distinct spatial patterns, ranging from uniform contributions across the fragment (e.g., AR and TP53) to highly localized activity at proximal or distal regions (e.g., SPDEF in proximal bins). These position-dependent weight profiles may reflect variation in TF interactions with RNA Pol II, distance-dependent cooperativity, or spatial constraints on enhancer grammar. Position-dependent regulatory effects of TF binding have been demonstrated experimentally by Duttke et al., who found that several TFs showed different or opposing effects depending on binding-site position relative to the TSS, whereas p53 showed a comparatively position-independent effect [50]. The similarly position-invariant profile of TP53 in our model is notable, although our positional features reflect binding within the downstream STARR-seq insert rather than in the immediate vicinity of the TSS.

Despite its predictive power, the model has limitations inherent to motif-based approaches. Because it relies strictly on sequence motifs, it may fail to capture indirect TF binding mediated by protein–protein interactions rather than direct DNA contact. The linear formulation also represents a first-order approximation of transcriptional regulation and does not explicitly model cooperative or antagonistic interactions between TFs. However, large-scale MPRA experiments found that TF effects were generally additive and that most enhancer activity did not appear to require specific TF–TF interactions [51], providing empirical support for this simplifying assumption. The model also does not fully capture the inherently nonlinear relationship between TF binding and transcriptional output, although the use of activation functions introduces some nonlinearity in mapping binding energy to occupancy. Furthermore, the inferred weights should be interpreted in the context of motif similarity, as TFs with highly similar binding preferences may share or redistribute contribution weights. Finally, the model was trained on STARR-seq data from a plasmid-based reporter assay, which may not fully recapitulate chromatin context or long-range interactions present in the native genome.

However, the simplicity and interpretability of the model are key advantages. In contrast to deep learning approaches, which often achieve high predictive performance at the cost of biological transparency, this framework enables direct attribution of regulatory effects to specific TFs and sequence features with no additional need for computation to disentangle contributions. As such, it is particularly well suited for generating biological insight rather than solely optimizing predictive accuracy, and it facilitates hypothesis generation and downstream experimental validation.

This framework can be extended to other hormone-driven systems and cell types, such as estrogen receptor (ER) signaling in breast cancer cell lines, to compare regulatory programs across biological contexts. Future work could incorporate higher-order interaction terms to explicitly model cooperative TF binding, potentially improving both biological interpretability and predictive performance.

## Supporting information

40-enhancer mutational profiles

## Data Availability

The STARR-seq enhancer activity data used for model training were obtained from Huang et al. [34] and are publicly available. The saturation-mutagenesis data are available in NCBI’s Gene Expression Omnibus (GEO) under accession GSE335266, which will be made public upon publication of the associated mutagenesis study (Tekoglu et al., manuscript in preparation). Transcription factor expression data were obtained from the DepMap portal (Public 25Q2 release). Motif models were obtained from the JASPAR 2024 CORE and HOCOMOCO v12 CORE databases. Prostate cancer risk variants were obtained from the GWAS Catalog (MONDO 0008315). The code implementing the model and analyses is available at https://github.com/HodaTb/AR_motif_model.

## Supplementary Data statement

Supplementary Data are available with this preprint.

## References

[1] Samuel A Lambert, Arttu Jolma, Laura F Campitelli, Pratyush K Das, Yimeng Yin, Mihai Albu, Xiaoting Chen, Jussi Taipale, Timothy R Hughes, and Matthew T Weirauch. The human transcription factors. Cell, 172(4):650–665, 2018.

[2] Francois Spitz and Eileen EM Furlong. Transcription factors: from enhancer binding to developmental control. Nature reviews genetics, 13(9):613–626, 2012.

[3] Kanchan Vishnoi, Navin Viswakarma, Ajay Rana, and Basabi Rana. Transcription factors in cancer development and therapy. Cancers, 12(8):2296, 2020.

[4] Emily Zboril, Hannah Yoo, Lizhen Chen, and Zhijie Liu. Dynamic interactions of transcription factors and enhancer reprogramming in cancer progression. Frontiers in oncology, 11:753051, 2021.

[5] Cynthia A Heinlein and Chawnshang Chang. Androgen receptor in prostate cancer. Endocrine reviews, 25(2):276–308, 2004.

[6] MH Tan, Jun Li, H Eric Xu, Karsten Melcher, and Eu-leong Yong. Androgen receptor: structure, role in prostate cancer and drug discovery. Acta Pharmacologica Sinica, 36(1):3–23, 2015.

[7] Ken-ichi Takayama and Satoshi Inoue. Transcriptional network of androgen receptor in prostate cancer progression. International Journal of Urology, 20(8):756–768, 2013.

[8] Doğancan Özturan, Tunç Morova and Nathan A Lack. Androgen receptor-mediated transcription in prostate cancer. Cells, 11(5):898, 2022.

[9] Daisuke Obinata, Kenichi Takayama, Satoshi Inoue, and Satoru Takahashi. Exploring androgen receptor signaling pathway in prostate cancer: A path to new discoveries. International Journal of Urology, 31(6):590–597, 2024.

[10] Steve Paltoglou, Rajdeep Das, Scott L Townley, Theresa E Hickey, Gerard A Tarulli, Isabel Coutinho, Rayzel Fernandes, Adrienne R Hanson, Iza Denis, Jason S Carroll, et al. Novel androgen receptor coregulator grhl2 exerts both oncogenic and antimetastatic functions in prostate cancer. Cancer research, 77(13):3417–3430, 2017.

[11] Hannelore V Heemers and Donald J Tindall. Androgen receptor (ar) coregulators: a diversity of functions converging on and regulating the ar transcriptional complex. Endocrine reviews, 28(7):778–808, 2007.

[12] John D Norris, Ching-Yi Chang, Bryan M Wittmann, Rebecca S Kunder, Huaxia Cui, Daju Fan, James D Joseph, and Donald P McDonnell. The homeodomain protein hoxb13 regulates the cellular response to androgens. Molecular cell, 36(3):405–416, 2009.

[13] Natalya V Guseva, Oskar W Rokhlin, Thomas B Bair, Rebecca B Glover, and Michael B Cohen. Inhibition of p53 expression modifies the specificity of chromatin binding by the androgen receptor. Oncotarget, 3(2):183, 2012.

[14] Kerim Yavuz and Nathan A Lack. Coregulators determine androgen receptor activity in prostate cancer. Bioscience Reports, 45(08):BSR20253197, 2025.

[15] Mark M Pomerantz, Fugen Li, David Y Takeda, Romina Lenci, Apurva Chonkar, Matthew Chabot, Paloma Cejas, Francisca Vazquez, Jennifer Cook, Ramesh A Shivdasani, et al. The androgen receptor cistrome is extensively reprogrammed in human prostate tumorigenesis. Nature genetics, 47(11):1346–1351, 2015.

[16] Peter J Park. Chip–seq: advantages and challenges of a maturing technology. Nature reviews genetics, 10(10):669–680, 2009.

[17] Peter J Skene and Steven Henikoff. An efficient targeted nuclease strategy for high-resolution mapping of dna binding sites. elife, 6:e21856, 2017.

[18] Cosmas D Arnold, Daniel Gerlach, Christoph Stelzer, L ukasz M Boryń, Martina Rath, and Alexander Stark. Genome-wide quantitative enhancer activity maps identified by starr-seq. Science, 339(6123):1074–1077, 2013.

[19] Lacramioara Bintu, Nicolas E Buchler, Hernan G Garcia, Ulrich Gerland, Terence Hwa, Jane Kondev, Thomas Kuhlman, and Rob Phillips. Transcriptional regulation by the numbers: applications. Current opinion in genetics & development, 15(2):125–135, 2005.

[20] Lacramioara Bintu, Nicolas E Buchler, Hernan G Garcia, Ulrich Gerland, Terence Hwa, Jańe Kondev, and Rob Phillips. Transcriptional regulation by the numbers: models. Current opinion in genetics & development, 15(2):116–124, 2005.

[21] Pankaj Gautam and Sudipta Kumar Sinha. Anticipating response function in gene regulatory networks. Journal of the Royal Society Interface, 18(179), 2021.

[22] Amin Safaeesirat, Hoda Taeb, Emirhan Tekoglu, Tunc Morova, Nathan A Lack, and Eldon Emberly. Inference of transcriptional regulation from starr-seq data. Physical Review E, 111(2):024402, 2025.

[23] Muir Morrison, Manuel Razo-Mejia, and Rob Phillips. Reconciling kinetic and thermodynamic models of bacterial transcription. PLoS computational biology, 17(1):e1008572, 2021.

[24] Babak Alipanahi, Andrew Delong, Matthew T Weirauch, and Brendan J Frey. Predicting the sequence specificities of dna-and rna-binding proteins by deep learning. Nature biotechnology, 33(8):831–838, 2015.

[25] Žiga Avsec, Melanie Weilert, Avanti Shrikumar, Sabrina Krueger, Amr Alexandari, Khyati Dalal, Robin Fropf, Charles McAnany, Julien Gagneur, Anshul Kundaje, et al. Base-resolution models of transcription-factor binding reveal soft motif syntax. Nature genetics, 53(3):354–366, 2021.

[26] Bernardo P De Almeida, Franziska Reiter, Michaela Pagani, and Alexander Stark. Deepstarr predicts enhancer activity from dna sequence and enables the de novo design of synthetic enhancers. Nature genetics, 54(5):613–624, 2022.

[27] Liyuan Shu, Jiao Tang, Xiaoyu Guan, and Daoqiang Zhang. A comprehensive survey of genome language models in bioinformatics. Briefings in Bioinformatics, 27(1):bbaf724, 2026.

[28] Gherman Novakovsky, Nick Dexter, Maxwell W Libbrecht, Wyeth W Wasserman, and Sara Mostafavi. Obtaining genetics insights from deep learning via explainable artificial intelligence. Nature Reviews Genetics, 24(2):125–137, 2023.

[29] Avanti Shrikumar, Peyton Greenside, and Anshul Kundaje. Learning important features through propagating activation differences. In International conference on machine learning, pages 3145–3153. PMlR, 2017.

[30] Avanti Shrikumar, Katherine Tian, Žiga Avsec, Anna Shcherbina, Abhimanyu Banerjee, Mahfuza Sharmin, Surag Nair, and Anshul Kundaje. Technical note on transcription factor motif discovery from importance scores (tf-modisco) version 0.5. 6.5. arXiv preprint arXiv:1811.00416, 2018.

[31] Carl G de Boer, Eeshit Dhaval Vaishnav, Ronen Sadeh, Esteban Luis Abeyta, Nir Friedman, and Aviv Regev. Deciphering eukaryotic gene-regulatory logic with 100 million random promoters. Nature biotechnology, 38(1):56–65, 2020.

[32] Paola Cornejo-Páramo, Xuan Zhang, Lithin Louis, Zelun Li, Yihua Yang, and Emily S Wong. Motif-based models accurately predict cell type-specific distal regulatory elements. Nature communications, 16(1):10370, 2025.

[33] Mike Kirby, Ceri Hirst, and ED Crawford. Characterising the castration-resistant prostate cancer population: a systematic review. International journal of clinical practice, 65(11):1180–1192, 2011.

[34] Chia-Chi Flora Huang, Shreyas Lingadahalli, Tunc Morova, Dogancan Ozturan, Eugene Hu, Ivan Pak Lok Yu, Simon Linder, Marlous Hoogstraat, Suzan Stelloo, Funda Sar, et al. Functional mapping of androgen receptor enhancer activity. Genome biology, 22(1):149, 2021.

[35] Allison Piovesan, Maria Chiara Pelleri, Francesca Antonaros, Pierluigi Strippoli, Maria Caracausi, and Lorenza Vitale. On the length, weight and gc content of the human genome. BMC research notes, 12(1):106, 2019.

[36] Broad Institute. Depmap public 25q2 dataset. https://depmap.org/portal/download/all/, 2025. Accessed: 2026-03-15.

[37] Rand Arafeh, Tsukasa Shibue, Joshua M Dempster, William C Hahn, and Francisca Vazquez. The present and future of the cancer dependency map. Nature Reviews Cancer, 25(1):59–73, 2025.

[38] Ieva Rauluseviciute, Rafael Riudavets-Puig, Romain Blanc-Mathieu, Jaime A Castro-Mondragon, Katalin Ferenc, Vipin Kumar, Roza Berhanu Lemma, Jérémy Lucas, Jeanne Chèneby, Damir Baranasic, et al. Jaspar 2024: 20th anniversary of the open-access database of transcription factor binding profiles. Nucleic acids research, 52(D1):D174–D182, 2024.

[39] Ilya E Vorontsov, Irina A Eliseeva, Arsenii Zinkevich, Mikhail Nikonov, Sergey Abramov, Alexandr Boytsov, Vasily Kamenets, Alexandra Kasianova, Semyon Kolmykov, Ivan S Yevshin, et al. Hocomoco in 2024: a rebuild of the curated collection of binding models for human and mouse transcription factors. Nucleic Acids Research, 52(D1):D154–D163, 2024.

[40] Xiaoyan Ma, Daphne Ezer, Carmen Navarro, and Boris Adryan. Reliable scaling of position weight matrices for binding strength comparisons between transcription factors. BMC bioinformatics, 16(1):265, 2015.

[41] Martín Abadi, Ashish Agarwal, Paul Barham, Eugene Brevdo, Zhifeng Chen, Craig Citro, Greg S Corrado, Andy Davis, Jeffrey Dean, Matthieu Devin, et al. Tensorflow: Large-scale machine learning on heterogeneous distributed systems. arXiv preprint arXiv:1603.04467, 2016.

[42] Amin Safaeesirat, Hoda Taeb, and Eldon Emberly. Inference of enhancer-specific transcription factor interactions from gene expression data using a biophysical model. bioRxiv, pages 2026–06, 2026.

[43] Fabian Pedregosa, Gaël Varoquaux, Alexandre Gramfort, Vincent Michel, Bertrand Thirion, Olivier Grisel, Mathieu Blondel, Peter Prettenhofer, Ron Weiss, Vincent Dubourg, et al. Scikit-learn: Machine learning in python. the Journal of machine Learning research, 12:2825–2830, 2011.

[44] Elliot Sollis, Abayomi Mosaku, Ala Abid, et al. The nhgri-ebi gwas catalog: knowledgebase and deposition resource. Nucleic Acids Research, 51(D1):D977–D985, 2023.

[45] Angela S Hinrichs, Donna Karolchik, Robert Baertsch, et al. The ucsc genome browser database: update 2006. Nucleic Acids Research, 34(Database issue):D590–D598, 2006.

[46] Peter Oettgen, Eduardo Finger, Zijie Sun, Yasmin Akbarali, Usanee Thamrongsak, Jay Boltax, Franck Grall, Antoinise Dube, Avi Weiss, Lawrence Brown, et al. Pdef, a novel prostate epithelium-specific ets transcription factor, interacts with the androgen receptor and activates prostate-specific antigen gene expression. Journal of Biological Chemistry, 275(2):1216–1225, 2000.

[47] Mathieu Lupien, Jérôme Eeckhoute, Clifford A Meyer, Qianben Wang, Yong Zhang, Wei Li, Jason S Carroll, X Shirley Liu, and Myles Brown. Foxa1 translates epigenetic signatures into enhancer-driven lineage-specific transcription. Cell, 132(6):958–970, 2008.

[48] Johanna Siegel, Michael Fritsche, Sabine Mai, Gerhard Brandner, and Ralf D Hess. Enhanced p53 activity and accumulation in response to dna damage upon dna transfection. Oncogene, 11(7):1363–1370, 1995.

[49] Thomas J Hoffmann, Michael N Passarelli, Rebecca E Graff, Nima C Emami, Lori C Sakoda, Eric Jorgenson, Laurel A Habel, Jun Shan, Dilrini K Ranatunga, Charles P Quesenberry, et al. Genome-wide association study of prostate-specific antigen levels identifies novel loci independent of prostate cancer. Nature communications, 8(1):14248, 2017.

[50] Sascha H Duttke, Carlos Guzman, Max Chang, Nathaniel P Delos Santos, Bayley R McDonald, Jialei Xie, Aaron F Carlin, Sven Heinz, and Christopher Benner. Position-dependent function of human sequence-specific transcription factors. Nature, 631(8022):891–898, 2024.

[51] Biswajyoti Sahu, Tuomo Hartonen, Päivi Pihlajamaa, Bei Wei, Kashyap Dave, Fangjie Zhu, Eevi Kaasinen, Katja Lidschreiber, Michael Lidschreiber, Carsten O Daub, et al. Sequence determinants of human gene regulatory elements. Nature genetics, 54(3):283–294, 2022.

