## Supplementary Material for "Motif-based model of transcription predicts effects of sequence variants in AR enhancers and reveals distinct functions for AR-associated transcription factors"

### S1 Biophysically motivated model

Consider an enhancer fragment $\nu$ where transcription factors (TFs) can bind to the DNA and regulate transcription through their effect on RNA polymerase II (Pol II). The binding state of TF $\mu$ is represented by a binary variable $\sigma_\mu \in \{0,1\}$, where $\sigma_\mu = 1$ indicates that the TF is bound and $\sigma_\mu = 0$ indicates that it is unbound. Similarly, $\sigma_p$ denotes the binding state of Pol II.

The effective binding energy of TF $\mu$ on fragment $\nu$ is denoted by $h_\mu^\nu$, and its chemical potential, which reflects its concentration in the cellular environment, is denoted by $G_\mu$. The interaction between TF $\mu$ and Pol II is denoted by $J_{\mu p}$. A more general model could also include TF–TF interactions, but for the linear model derived here we neglect these terms, setting $J_{\mu\eta} = 0$ for $\mu \neq \eta$. Nonzero TF–TF interactions would introduce higher-order contributions that are not represented in the linear regression model. For clarity we suppress the fragment index $\nu$ on sequence-dependent quantities throughout the derivation, restoring it in the final expression.

With this simplification, the Hamiltonian is

$$H(\{\sigma_\mu\}, \sigma_p) = -\sum_\mu (h_\mu + G_\mu)\sigma_\mu - \sum_\mu J_{\mu p}\sigma_\mu\sigma_p - (h_p + G_p)\sigma_p. \tag{1}$$

The partition function is

$$Z = \sum_{\{\sigma_\mu\},\sigma_p} \exp[-H(\{\sigma_\mu\}, \sigma_p)], \tag{2}$$

where we work in units with $k_B T = 1$. The binding probability of Pol II is

$$m_p = P(\sigma_p = 1) = \frac{1}{Z} \sum_{\{\sigma_\mu\},\sigma_p} \sigma_p \exp[-H(\{\sigma_\mu\}, \sigma_p)]. \tag{3}$$

To connect this model to the linear regression used in the main text, we use a mean-field approximation in which each TF binding variable is replaced by its average occupancy, $m_\mu = \langle \sigma_\mu \rangle$. At zeroth order in the TF–Pol II couplings $J_{\mu p}$, this occupancy is

$$m_\mu^{(0)} = \frac{1}{2} + \frac{1}{2}\tanh\left[\frac{1}{2}(h_\mu + G_\mu)\right]. \tag{4}$$

Using these zeroth-order TF occupancies, the mean-field occupancy of Pol II is

$$m_p = \frac{1}{2} + \frac{1}{2}\tanh\left[\frac{1}{2}\left(h_p + G_p + \sum_\mu J_{\mu p} m_\mu^{(0)}\right)\right]. \tag{5}$$

Defining the baseline Pol II occupancy in the absence of TF–Pol II interactions as

$$m_p^{(0)} = \frac{1}{2} + \frac{1}{2}\tanh\left[\frac{1}{2}(h_p + G_p)\right], \tag{6}$$

and expanding to first order in the TF–Pol II interactions gives

$$m_p \approx m_p^{(0)} + m_p^{(0)}\left(1 - m_p^{(0)}\right)\sum_\mu J_{\mu p} m_\mu^{(0)}. \tag{7}$$

Thus, to first order, Pol II occupancy depends additively on TF occupancies. We assume that the transcriptional output of fragment $\nu$, denoted $Y^\nu$, is proportional to the Pol II binding probability, and we use the aggregated TF binding score $\bar{\beta}_\mu^\nu$ defined in Eq. 8 as a sequence-derived proxy for the effective binding of TF $\mu$ on fragment $\nu$. Absorbing the baseline response, the TF–Pol II interaction strengths, and the remaining proportionality constants into effective fitted coefficients, we obtain

$$Y^\nu = \sum_\mu \bar{\beta}_\mu^\nu W_\mu + C. \tag{8}$$

A first-order expansion of the log-transformed transcriptional output around its baseline retains the same additive form. Because LogFC is defined as the difference between log-transformed activities under DHT and EtOH, it likewise retains an additive first-order form. This corresponds to the position-independent linear model in Eq. 9. In this interpretation, $W_\mu$ represents the effective contribution of TF $\mu$ to the modeled transcriptional response.

#### Optimal hyperparameters, cross-validation, and test performance

Table S1: Cross-validation mean squared error (MSE), optimal hyperparameters, and held-out test MSE for each model architecture. All models were optimized via 5-fold cross-validation on the training set.

| **Model** | **Architecture** | **Pooling** | **Activation** | **Motif Agg.** | $\delta$ | $\alpha$ | **CV MSE** | **Test MSE** |
|---|---|---|---|---|---|---|---|---|
| LogFC | Position-independent | Average | ReLU | Mean | 6 | $10^{1.0}$ | $0.21 \pm 0.01$ | 0.20 |
| LogDHT | Position-independent | Max | ReLU | Mean | 8 | $10^{4.5}$ | $1.26 \pm 0.03$ | 1.23 |
| LogEtOH | Position-independent | Max | ReLU | Mean | 8 | $10^{4.5}$ | $1.30 \pm 0.03$ | 1.27 |
| LogFC | Position-dependent | Max | ReLU | Mean | 6 | $10^{5}$ | $0.21 \pm 0.01$ | 0.20 |
| Mutagenesis | Position-independent | Average | ReLU | Mean | 4 | $10^{-1.5}$ | $0.22 \pm 0.05$ | 0.17 |

### Hyperparameter optimization for the LogFC model

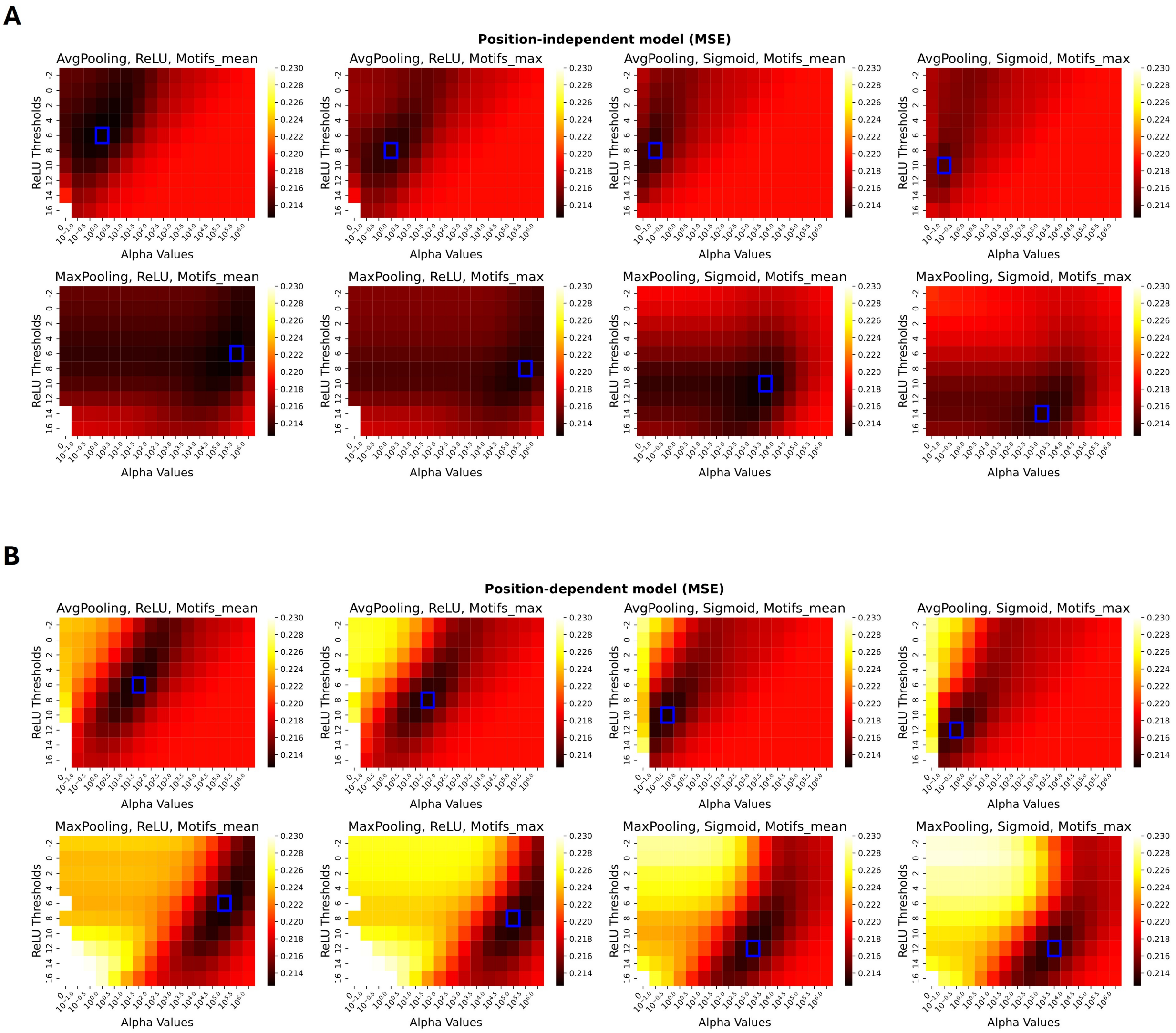


Figure S1: Heatmaps show cross-validation mean squared error (MSE) as a function of regularization strength ($\alpha$) and energy threshold ($\delta$) for the **(A)** position-independent and **(B)** position-dependent model architectures. Each panel represents one combination of spatial pooling (average or max), activation function (ReLU or Sigmoid), and motif aggregation strategy (mean or max). Blue boxes indicate the optimal hyperparameter combination for each configuration. The position-independent model achieved optimal performance with $\delta = 6$ and $\alpha = 1$ (average pooling, ReLU activation, mean aggregation), while the position-dependent model achieved optimal performance with $\delta = 6$ and $\alpha = 10^5$ (maximum pooling, ReLU activation, mean aggregation). Hyperparameter optimization for LogDHT and LogEtOH models showed similar patterns and are not shown.

### Fitted weights of LogEtOH model

**A**

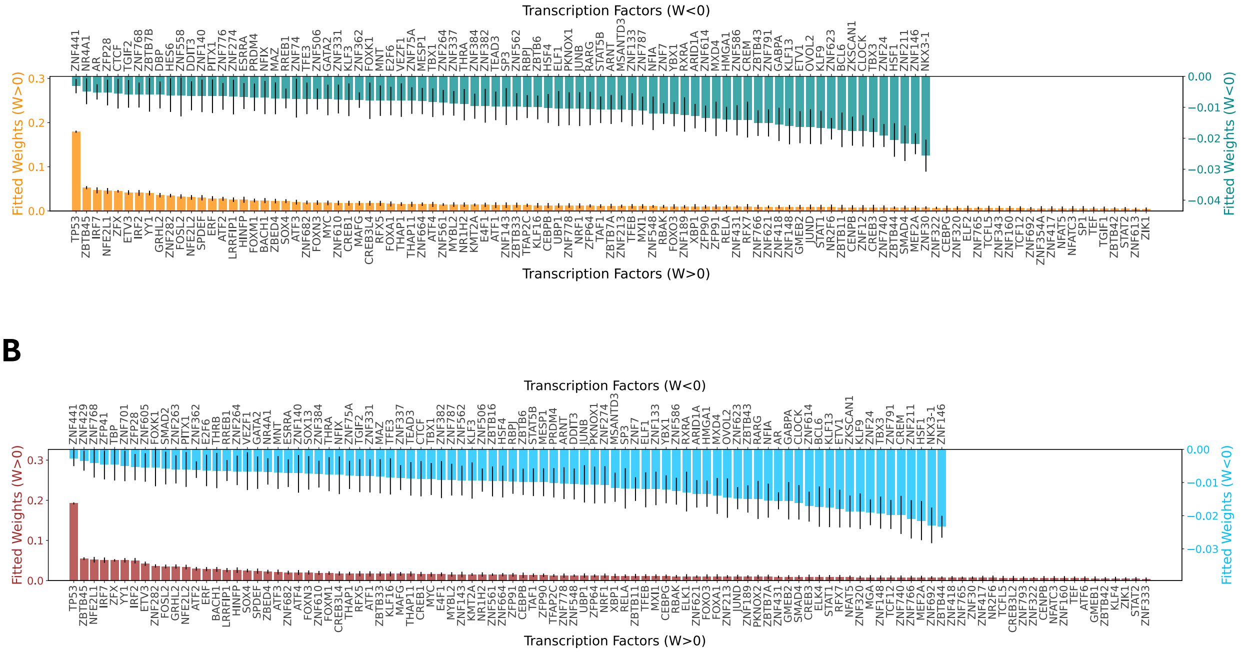


**B**

**C**

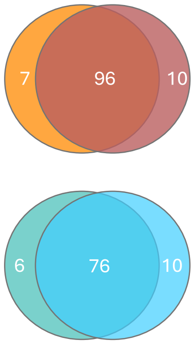


**D**

| Groups | Transcription Factors |
|---|---|
| LogEtOH (W>0) | ATF6, CREB3L2, ELK1, ELK4, GMEB1, KLF4, MGA, PKNOX2, ZNF333, ZNF793 |
| LogEtOH (W<0) | SMAD2, SOX13, TBP, THRB, ZBTB16, ZFP41, ZNF263, ZNF429, ZNF605, ZNF701 |
| LogDHT (W>0) | ELF2, SP1, TGIF1, ZNF12, ZNF343, ZNF354A, ZNF613 |
| LogDHT (W<0) | DBP, HES6, ZBTB7B, ZNF558, ZNF74, ZNF776 |

Figure S2: **A**, Ranked weights for the LogDHT model, representing TF contributions to overall enhancer activity. Bar heights indicate the mean weight $W_\mu$ across 1,000 bootstrap samples. **B**, Ranked weights for the LogEtOH model, representing TF contributions to baseline enhancer activity independent of AR, with bootstrap statistics as in **A**. **C**, Venn diagram showing the overlap of significant TFs between LogDHT and LogEtOH models. The top diagram compares positively weighted TFs, and the bottom compares negatively weighted TFs. **D**, Summary table of condition-specific TFs—those uniquely contributing to either DHT-stimulated activity (LogDHT) or baseline enhancer activity (LogEtOH).

### Hyperparameter optimization for the mutagenesis model

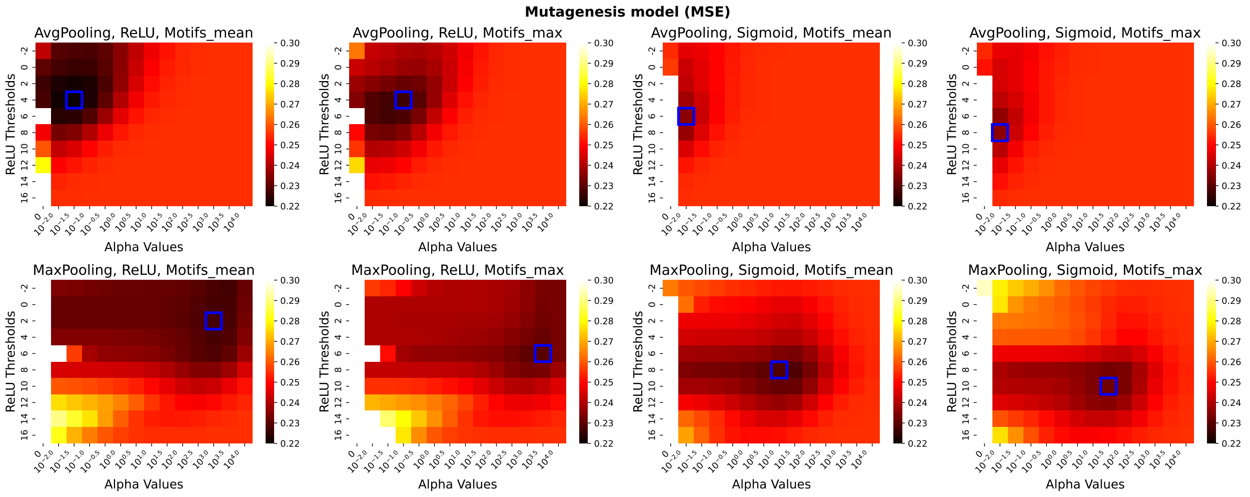


Figure S3: Hyperparameter optimization for the mutagenesis model. Heatmaps show cross-validation mean squared error (MSE) as a function of regularization strength ($\alpha$) and energy threshold ($\delta$) for the position-independent model trained directly on saturation mutagenesis data from 40 AR enhancers. Each panel represents one combination of spatial pooling (average or max), activation function (ReLU or Sigmoid), and motif aggregation strategy (mean or max). The blue box indicates the optimal hyperparameter combination: $\delta = 4$ and $\alpha = 10^{-1.5}$ (average pooling, ReLU activation, mean aggregation), achieving a cross-validation MSE of $0.22 \pm 0.05$.

### Correlation between position-independent and average position-dependent model weights for the LogFC model

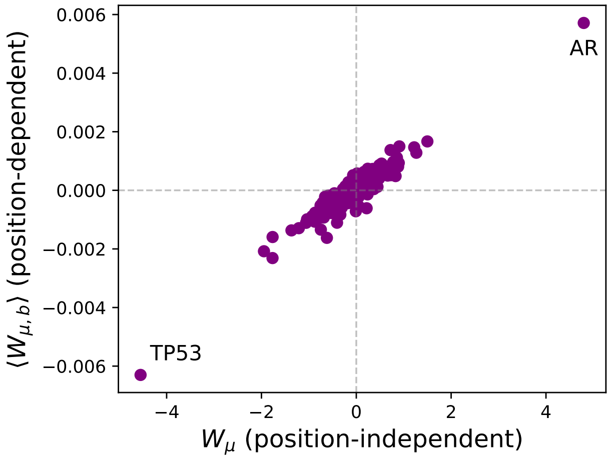


Figure S4: Each point represents one TF ($n = 293$). The high Pearson correlation coefficient (PCC = 0.95) indicates that position-dependent weights, when averaged across spatial bins, closely approximate position-independent weights, demonstrating overall consistency between the two modeling approaches. Notable outliers include AR (top right) with the strongest positive weight in both models, and TP53 (bottom left) with the strongest negative inducibility weight.

#### Summary of TF roles

Table S2: TF assignments by model weight significance across LogFC, LogDHT, and LogEtOH models.

| LogFC | LogDHT / LogEtOH | $n$ | Transcription Factors |
|---|---|---|---|
| *Inducibility + Absolute Activity (Shared Across Models)* | | | |
| + | both − | 16 | **AR**, **ARID1A**, CREM, CTCF, DDIT3, ESRRA, MESP1, NFIA, NR4A1, PRDM4, RARG, SP3, ZNF274, ZNF506, ZNF614, ZNF791 |
| + | both + | 8 | CENPB, **FOXA1**, FOXM1, FOXO3, SPDEF, ZNF322, ZNF740, ZNF766 |
| − | both + | 31 | ATF4, BACH1, CEBPB, CEBPG, E4F1, FOSL2, IRF2, IRF7, JUND, KLF16, NFAT5, NFE2L1, NFE2L2, NR1H2, NRF1, SMAD4, SOX4, STAT2, TAF1, TCF12, TEF, THAP1, **TP53**, YY1, ZBTB11, ZBTB45, ZFP90, ZFP91, ZFX, ZNF143, ZNF610 |
| − | both − | 13 | CLOCK, MNT, MXD4, NKX3-1, RREB1, STAT5B, VEZF1, ZNF264, ZNF384, ZNF623, ZNF7, ZNF768, ZNF787 |
| + | EtOH− only | 8 | SMAD2, SOX13, TBP, THRB, ZBTB16, ZFP41, ZNF263, ZNF429 |
| + | DHT+ only | 3 | SP1, TGIF1, ZNF343 |
| − | DHT− only | 2 | DBP, ZNF558 |
| − | EtOH+ only | 7 | ELK1, ELK4, GMEB1, KLF4, MGA, PKNOX2, ZNF793 |
| *Inducibility-specific (LogFC Exclusive)* | | | |
| + | not significant | 17 | ATF7, ELF3, FOXK2, FOXP1, FOXP4, HIF1A, **HOXB13**, KLF11, MLX, NFIC, NFKB1, PPARD, REST, USF1, ZNF276, ZNF467, ZNF714 |
| − | not significant | 11 | HMBOX1, IRF6, JUN, MNX1, MZF1, NFKB2, SP2, SREBF1, SRF, ZNF266, ZNF302 |
| *Absolute-activity-specific (LogDHT/LogEtOH Exclusive)* | | | |

Continued on next page

Table S2 – continued from previous page

| LogFC | LogDHT / LogEtOH | $n$ | Transcription Factors |
|---|---|---|---|
| not significant | both + | 57 | ATF1, ATF2, ATF3, CREB1, CREB3, CREB3L4, ERF, ETV3, FOXN3, GMEB2, GRHL2, HINFP, KMT2A, LRRFIP1, MAFG, MEF2A, MXI1, MYBL2, MYC, NFATC3, NR2F6, RBAK, RELA, RFX5, RFX7, STAT1, TCFL5, TFAP2C, TFEB, THAP11, UBP1, XBP1, ZBED4, ZBTB33, ZBTB42, ZBTB44, ZBTB7A, ZFP64, ZIK1, ZNF148, ZNF160, ZNF189, ZNF213, ZNF282, ZNF30, ZNF320, ZNF417, ZNF418, ZNF431, ZNF548, ZNF561, ZNF621, ZNF664, ZNF682, ZNF692, ZNF765, ZNF778 |
| not significant | both − | 47 | ARNT, BCL6, E2F6, ELF1, ETV1, FOXK1, GABPA, GATA2, HMGA1, HSF1, HSF4, JUNB, KLF13, KLF3, KLF9, MAZ, MSANTD3, NFIX, OVOL2, PITX1, PKNOX1, RBPJ, RXRA, TBX1, TBX3, TEAD3, TFE3, TGIF2, THRA, YBX1, ZBTB43, ZBTB6, ZFP28, ZKSCAN1, ZNF133, ZNF140, ZNF146, ZNF211, ZNF24, ZNF331, ZNF337, ZNF362, ZNF382, ZNF441, ZNF562, ZNF586, ZNF75A |
| not significant | DHT+ only | 4 | ELF2, ZNF12, ZNF354A, ZNF613 |
| not significant | DHT− only | 4 | HES6, ZBTB7B, ZNF74, ZNF776 |
| not significant | EtOH+ only | 3 | ATF6, CREB3L2, ZNF333 |
| not significant | EtOH− only | 2 | ZNF605, ZNF701 |

**Position-dependent TF binding preferences for the LogDHT and LogEtOH models**

Table S3: Four prostate cancer GWAS risk alleles prioritized based on predicted effects on ARBS inducibility. Each variant had a predicted ΔLogFC below the mean minus 1 SD of the substitution reference distribution constructed across the same 17 ARBS regions.

| SNP-Risk Allele | Position (GRCh38) | Position in Region | Region LogFC | # of Studies | $-\log_{10}(P)$ |
|---|---|---|---|---|---|
| rs72848121-G | chr5:172533408 | 83 | −0.36 | 2 | 8.70 |
| rs11986220-A | chr8:127519444 | 417 | −0.62 | 8 | 567.05 |
| rs1160267-G | chr8:23672008 | 254 | −0.09 | 10 | 207.00 |
| rs11665748-A | chr19:50851141 | 587 | 3.38 | 4 | 123.40 |

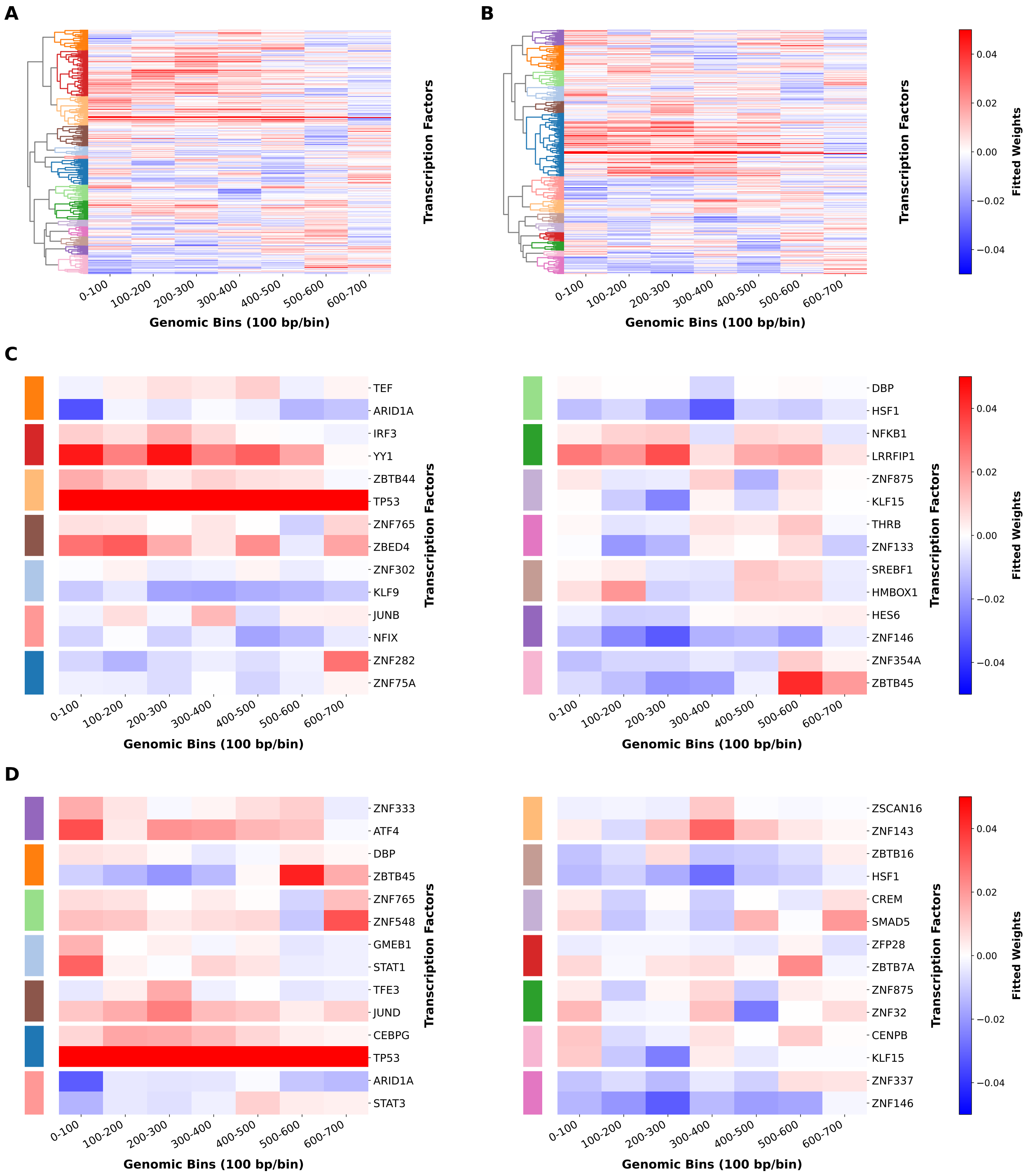


Figure S5: Analysis follows the same clustering approach as Fig. 5. **A,B**, Hierarchical clustering of position-dependent Ridge weights (7 spatial bins × 293 TFs) using 14 clusters, average linkage, and correlation distance for the (A) LogDHT and (B) LogEtOH models. **C,D**, Two representative TFs from each cluster are shown for the (C) LogDHT and (D) LogEtOH models: the TF closest to the cluster centroid and the TF with the largest absolute weight across spatial bins. Clusters are ordered according to the dendrogram leaf order and divided between two panels, with the first seven clusters on the left and the remaining seven on the right.
