## Supplementary material for "Motif-based model of transcription predicts effects of sequence variants in AR enhancers and reveals distinct functions for AR-associated transcription factors": 40-enhancer mutational profiles

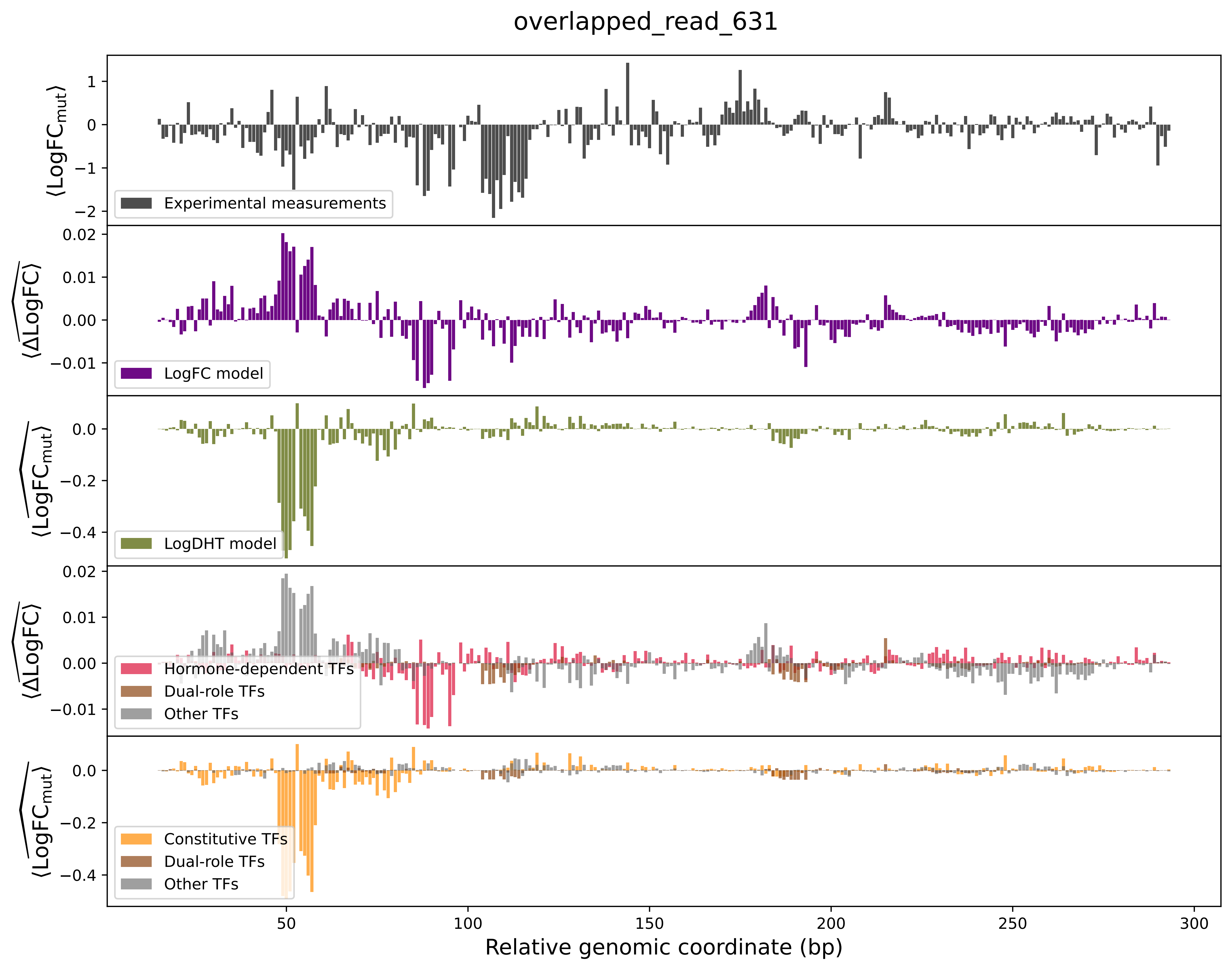

overlapped_read_631
⟨LogFC_mut⟩
Experimental measurements
⟨ΔLogFC⟩
LogFC model
⟨LogFC_mut⟩
LogDHT model
⟨ΔLogFC⟩
Hormone-dependent TFs
Dual-role TFs
Other TFs
⟨LogFC_mut⟩
Constitutive TFs
Dual-role TFs
Other TFs
Relative genomic coordinate (bp)

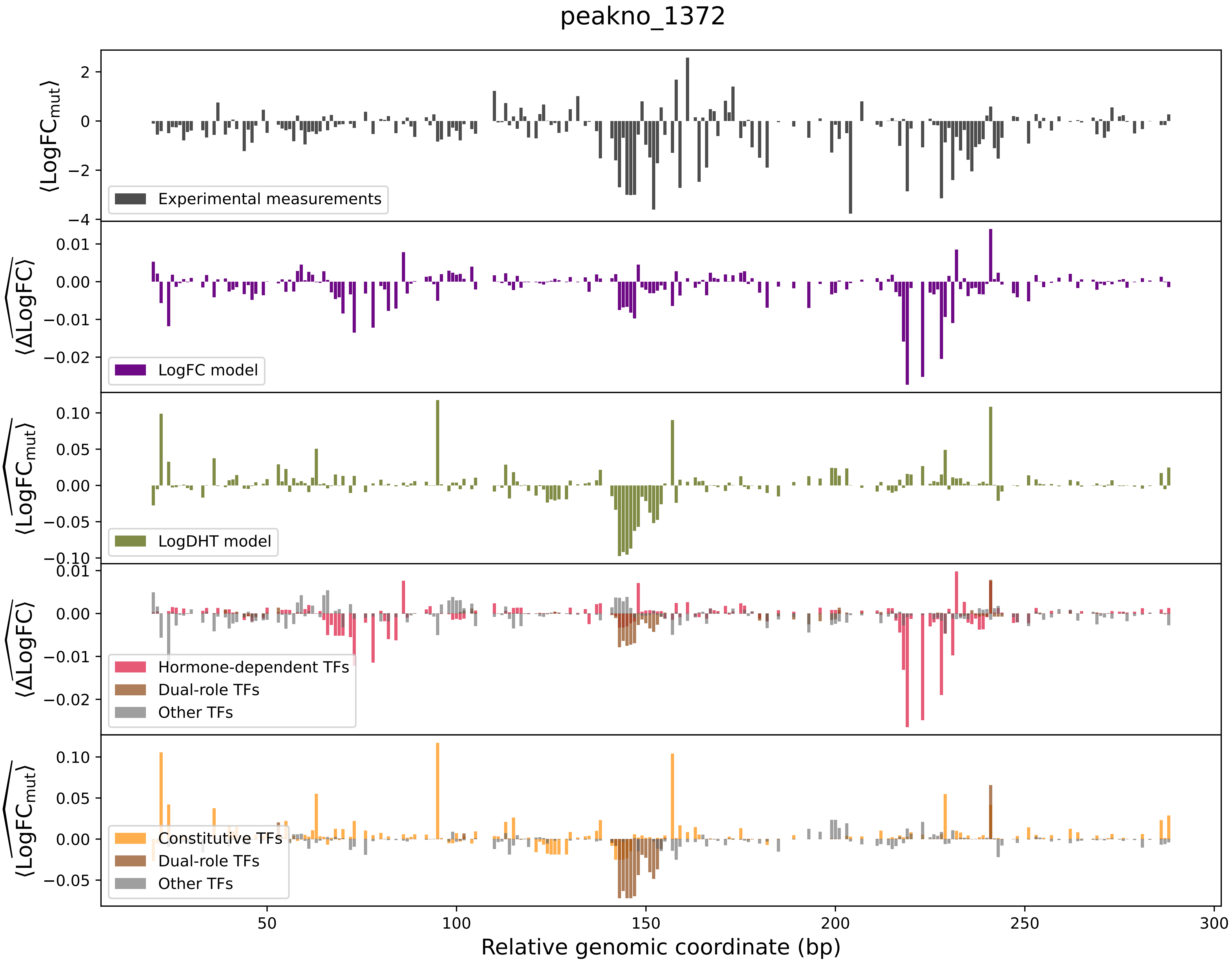
peakno_1372
Experimental measurements
LogFC model
LogDHT model
Hormone-dependent TFs
Dual-role TFs
Other TFs
Constitutive TFs
Dual-role TFs
Other TFs
⟨LogFC_mut⟩
⟨ΔLogFC⟩
⟨LogFC_mut⟩
⟨ΔLogFC⟩
⟨LogFC_mut⟩
Relative genomic coordinate (bp)

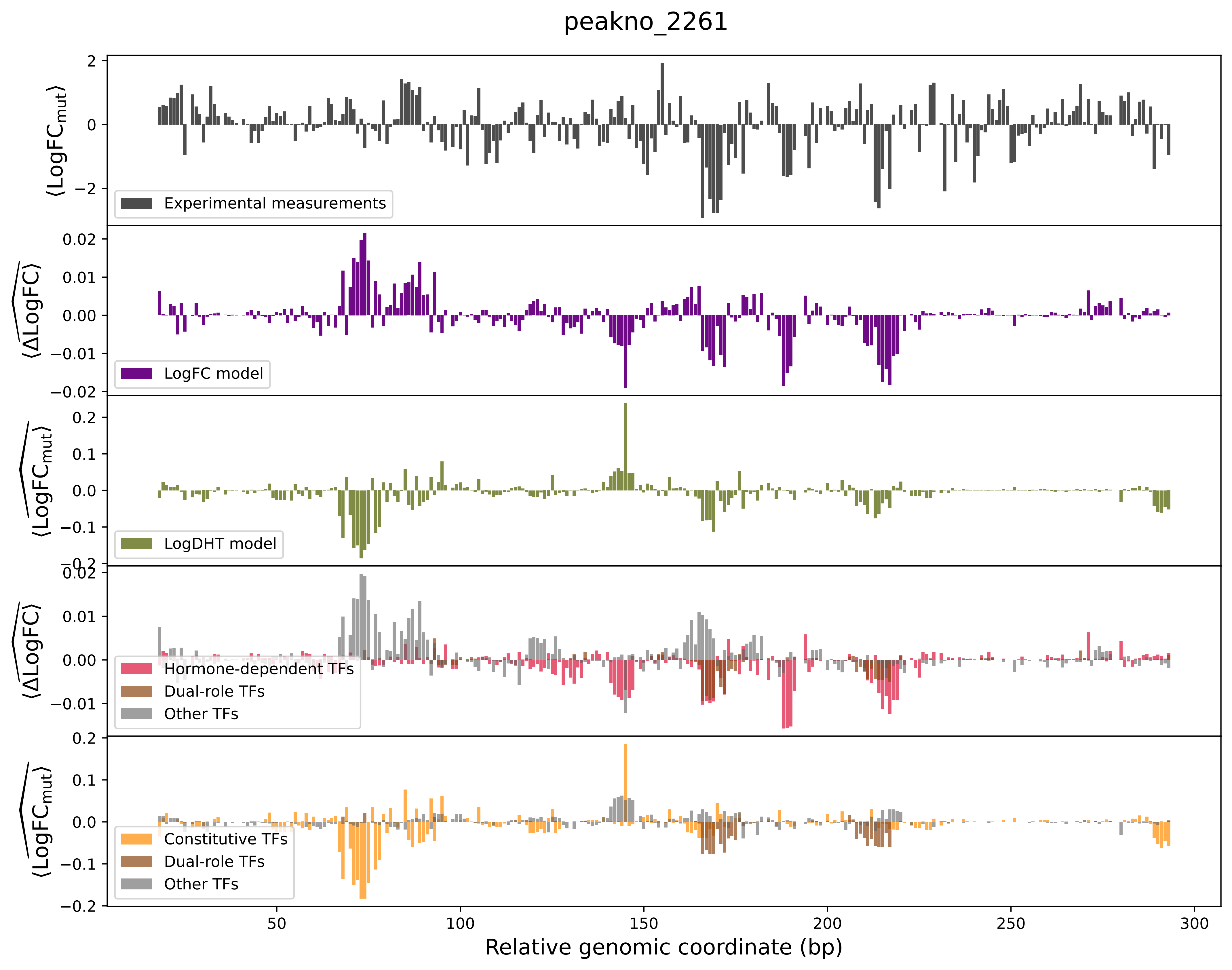
peakno_2261
Experimental measurements
LogFC model
LogDHT model
Hormone-dependent TFs
Dual-role TFs
Other TFs
Constitutive TFs
Dual-role TFs
Other TFs
Relative genomic coordinate (bp)
⟨LogFC$_{mut}$⟩
⟨$\widehat{\Delta LogFC}$⟩
⟨$\widehat{LogFC_{mut}}$⟩

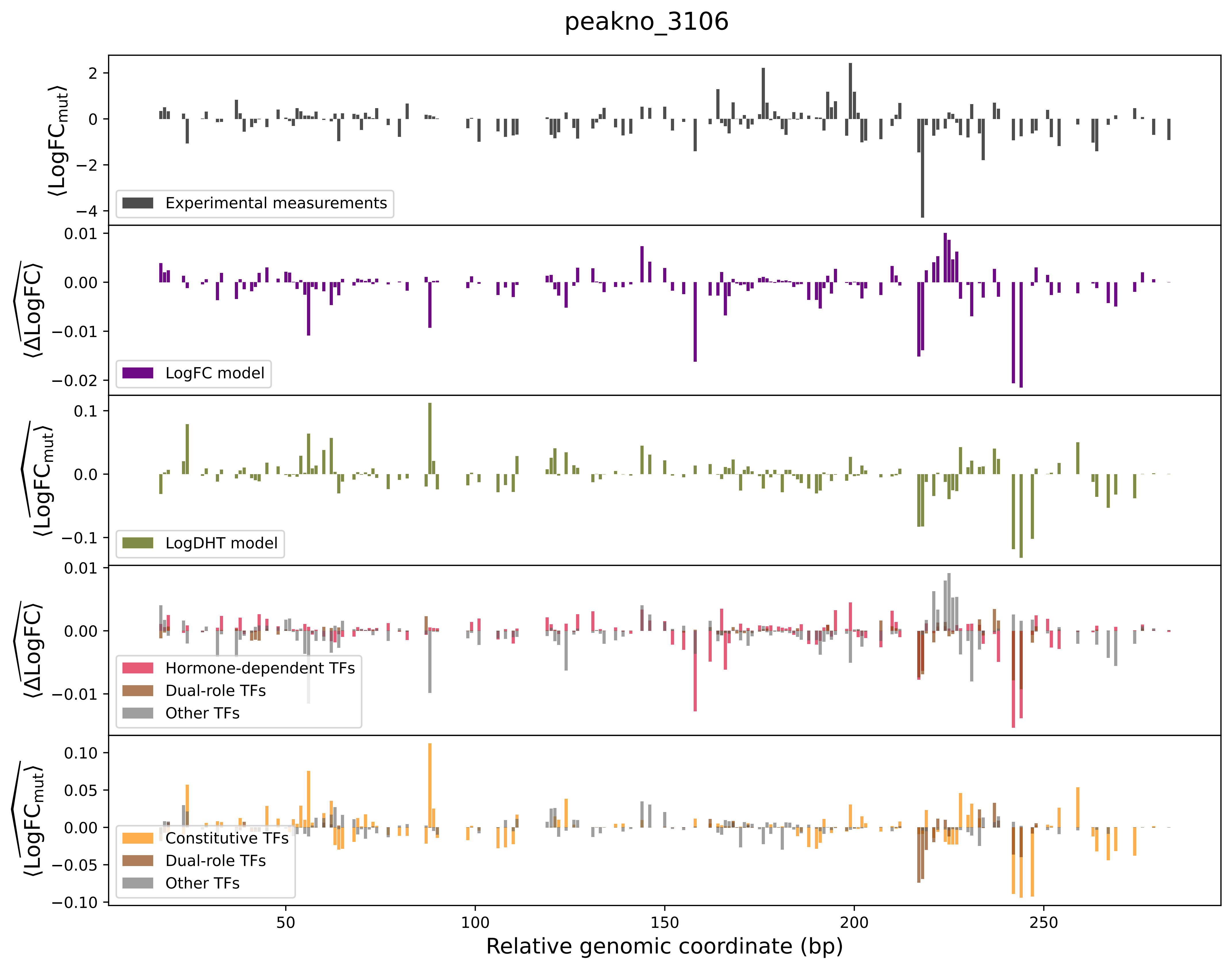

peakno_3106
⟨LogFC_mut⟩
Experimental measurements
⟨ΔLogFC⟩
LogFC model
⟨LogFC_mut⟩
LogDHT model
⟨ΔLogFC⟩
Hormone-dependent TFs
Dual-role TFs
Other TFs
⟨LogFC_mut⟩
Constitutive TFs
Dual-role TFs
Other TFs
Relative genomic coordinate (bp)

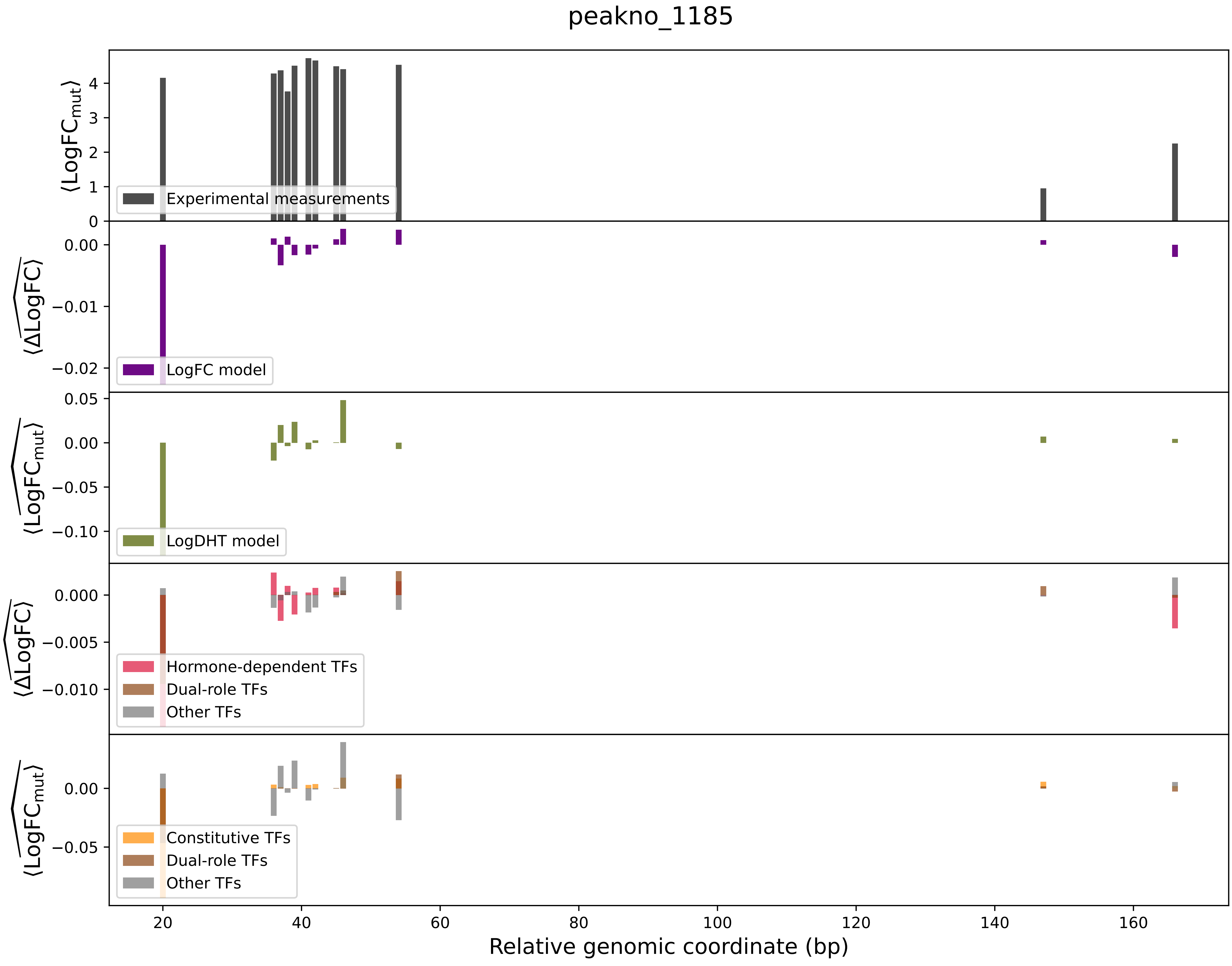

peakno_1185
⟨LogFC_mut⟩
Experimental measurements
⟨ΔLogFC⟩
LogFC model
⟨LogFC_mut⟩
LogDHT model
⟨ΔLogFC⟩
Hormone-dependent TFs
Dual-role TFs
Other TFs
⟨LogFC_mut⟩
Constitutive TFs
Dual-role TFs
Other TFs
Relative genomic coordinate (bp)

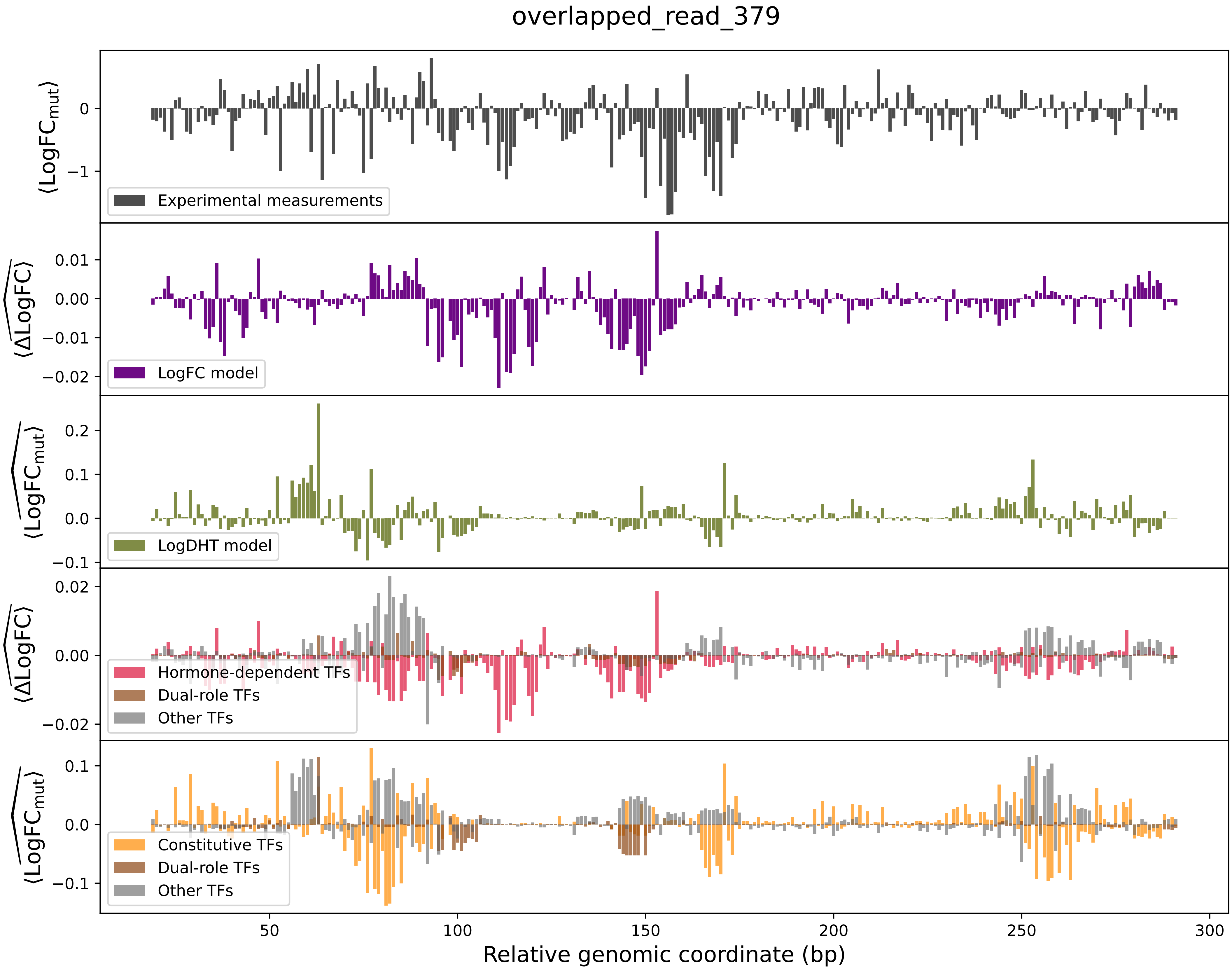

overlapped_read_379
⟨LogFC_mut⟩
Experimental measurements
⟨ΔLogFC⟩
LogFC model
⟨LogFC_mut⟩
LogDHT model
⟨ΔLogFC⟩
Hormone-dependent TFs
Dual-role TFs
Other TFs
⟨LogFC_mut⟩
Constitutive TFs
Dual-role TFs
Other TFs
Relative genomic coordinate (bp)

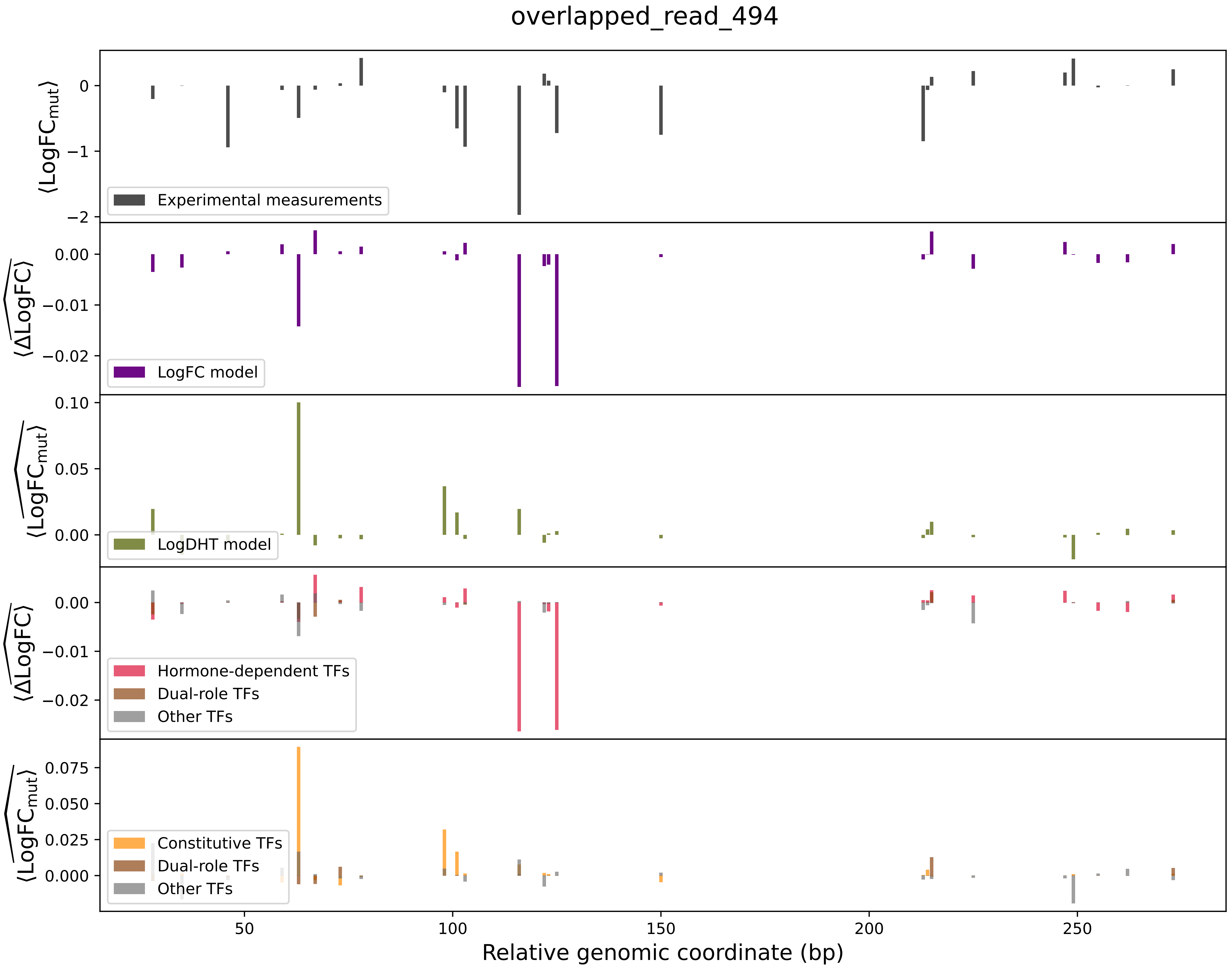
overlapped_read_494
Experimental measurements
LogFC model
LogDHT model
Hormone-dependent TFs
Dual-role TFs
Other TFs
Constitutive TFs
Dual-role TFs
Other TFs
$\langle \mathrm{LogFC_{mut}} \rangle$
$\langle \widehat{\Delta \mathrm{LogFC}} \rangle$
$\langle \widehat{\mathrm{LogFC_{mut}}} \rangle$
Relative genomic coordinate (bp)

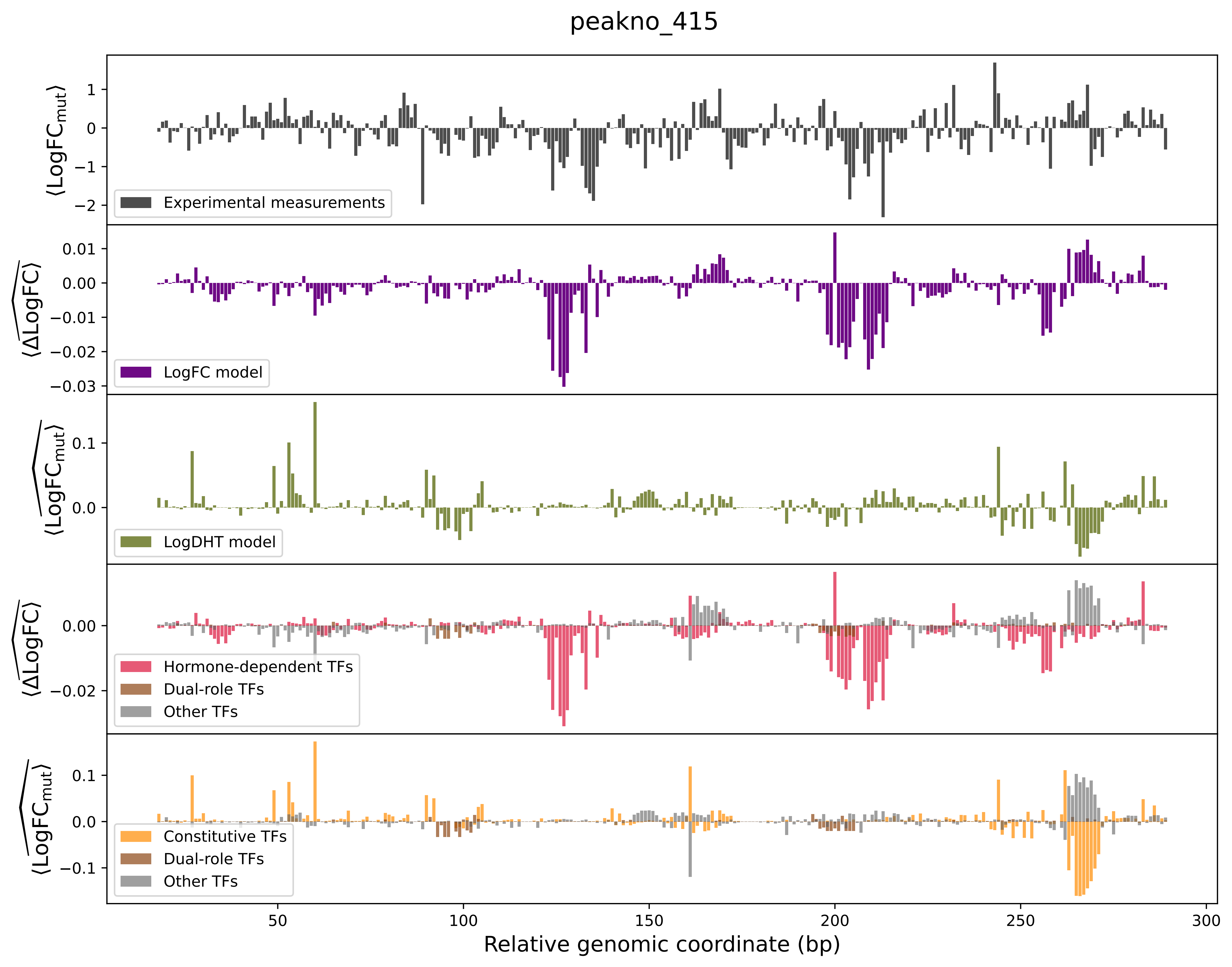

peakno_415
⟨LogFC_mut⟩
Experimental measurements
LogFC model
LogDHT model
Hormone-dependent TFs
Dual-role TFs
Other TFs
Constitutive TFs
Dual-role TFs
Other TFs
50
100
150
200
250
300
Relative genomic coordinate (bp)

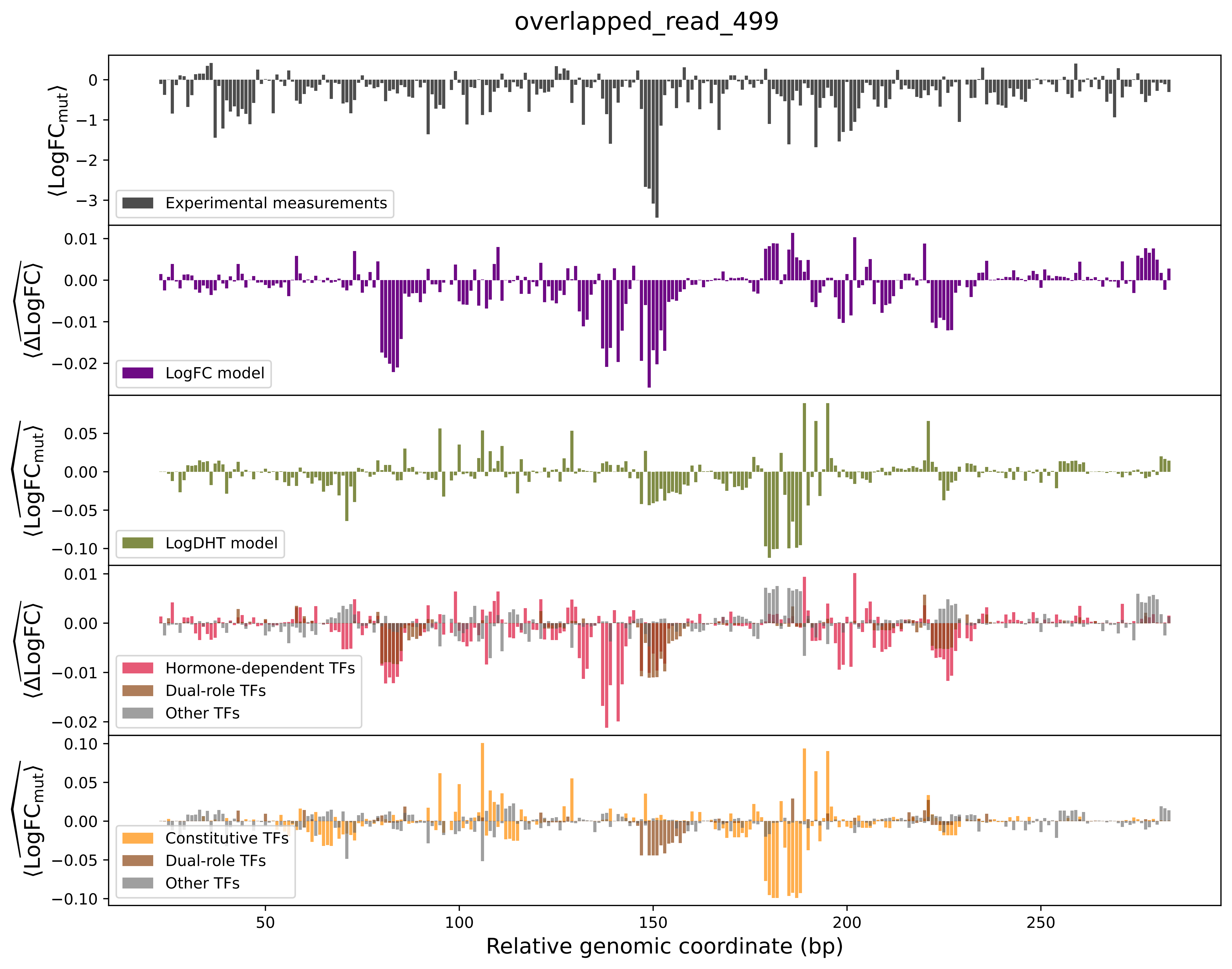
overlapped_read_499
⟨LogFCmut⟩
Experimental measurements
⟨ΔLogFC⟩
LogFC model
⟨LogFCmut⟩
LogDHT model
⟨ΔLogFC⟩
Hormone-dependent TFs
Dual-role TFs
Other TFs
⟨LogFCmut⟩
Constitutive TFs
Dual-role TFs
Other TFs
Relative genomic coordinate (bp)

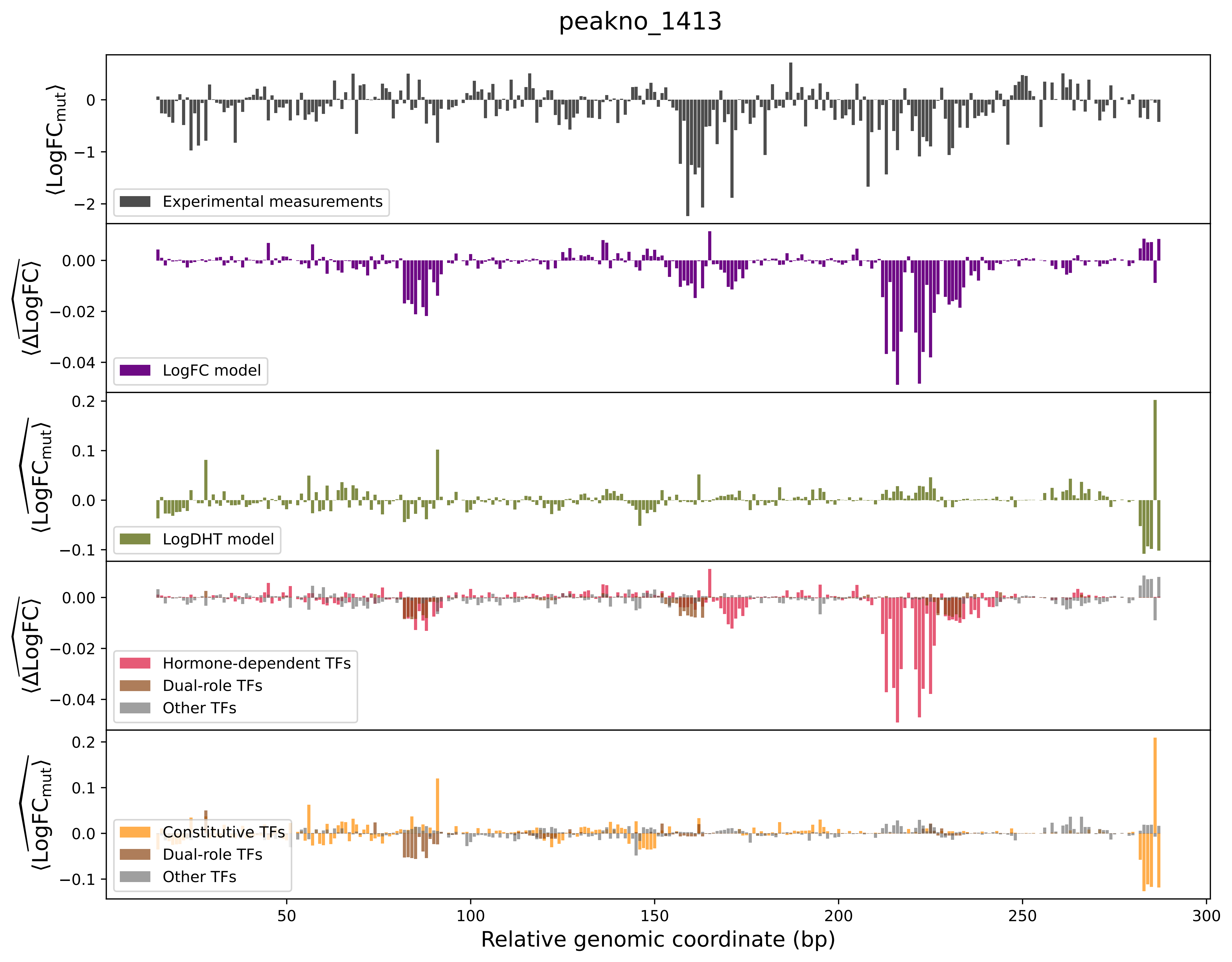

peakno_1413
Experimental measurements
LogFC model
LogDHT model
Hormone-dependent TFs
Dual-role TFs
Other TFs
Constitutive TFs
Dual-role TFs
Other TFs
Relative genomic coordinate (bp)

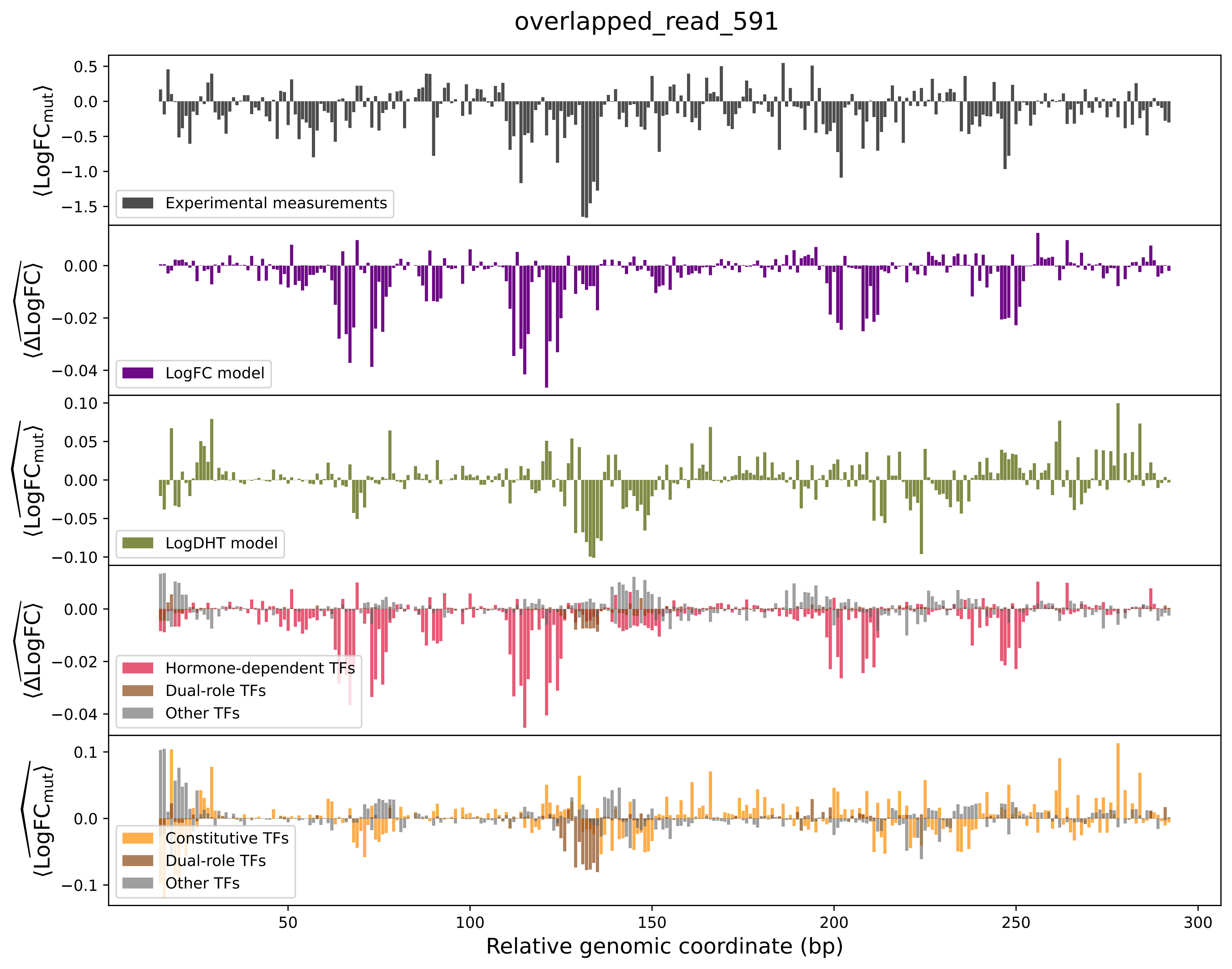

overlapped_read_591
0.5
0.0
−0.5
−1.0
−1.5
⟨LogFC_mut⟩
Experimental measurements
0.00
−0.02
−0.04
⟨ΔLogFC⟩
LogFC model
0.10
0.05
0.00
−0.05
−0.10
⟨LogFC_mut⟩
LogDHT model
0.00
−0.02
−0.04
⟨ΔLogFC⟩
Hormone-dependent TFs
Dual-role TFs
Other TFs
0.1
0.0
−0.1
⟨LogFC_mut⟩
Constitutive TFs
Dual-role TFs
Other TFs
50
100
150
200
250
300
Relative genomic coordinate (bp)

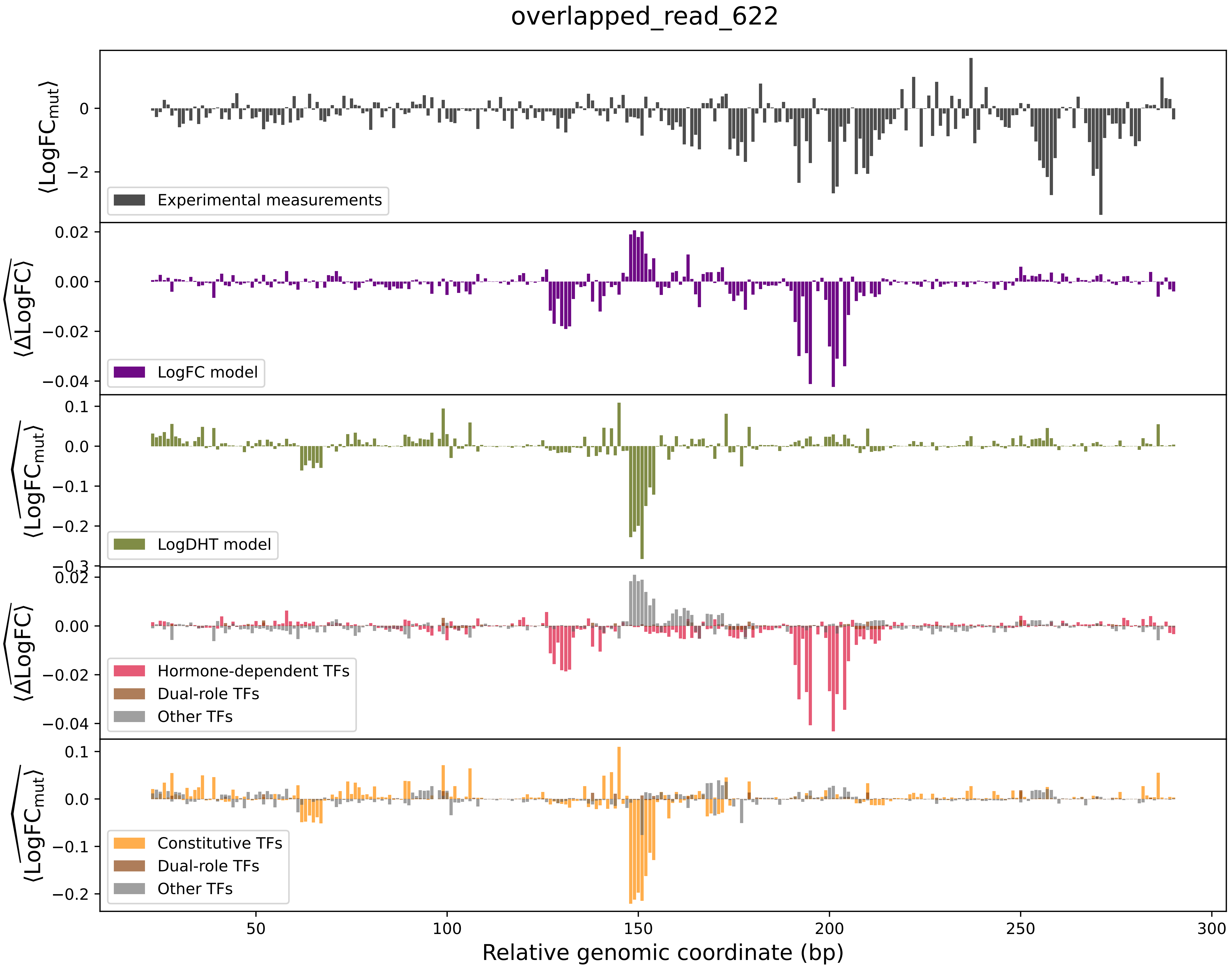
overlapped_read_622
⟨LogFC_mut⟩
0
−2
Experimental measurements
⟨ΔLogFC⟩
0.02
0.00
−0.02
−0.04
LogFC model
⟨LogFC_mut⟩
0.1
0.0
−0.1
−0.2
−0.3
LogDHT model
⟨ΔLogFC⟩
0.02
0.00
−0.02
−0.04
Hormone-dependent TFs
Dual-role TFs
Other TFs
⟨LogFC_mut⟩
0.1
0.0
−0.1
−0.2
Constitutive TFs
Dual-role TFs
Other TFs
50
100
150
200
250
300
Relative genomic coordinate (bp)

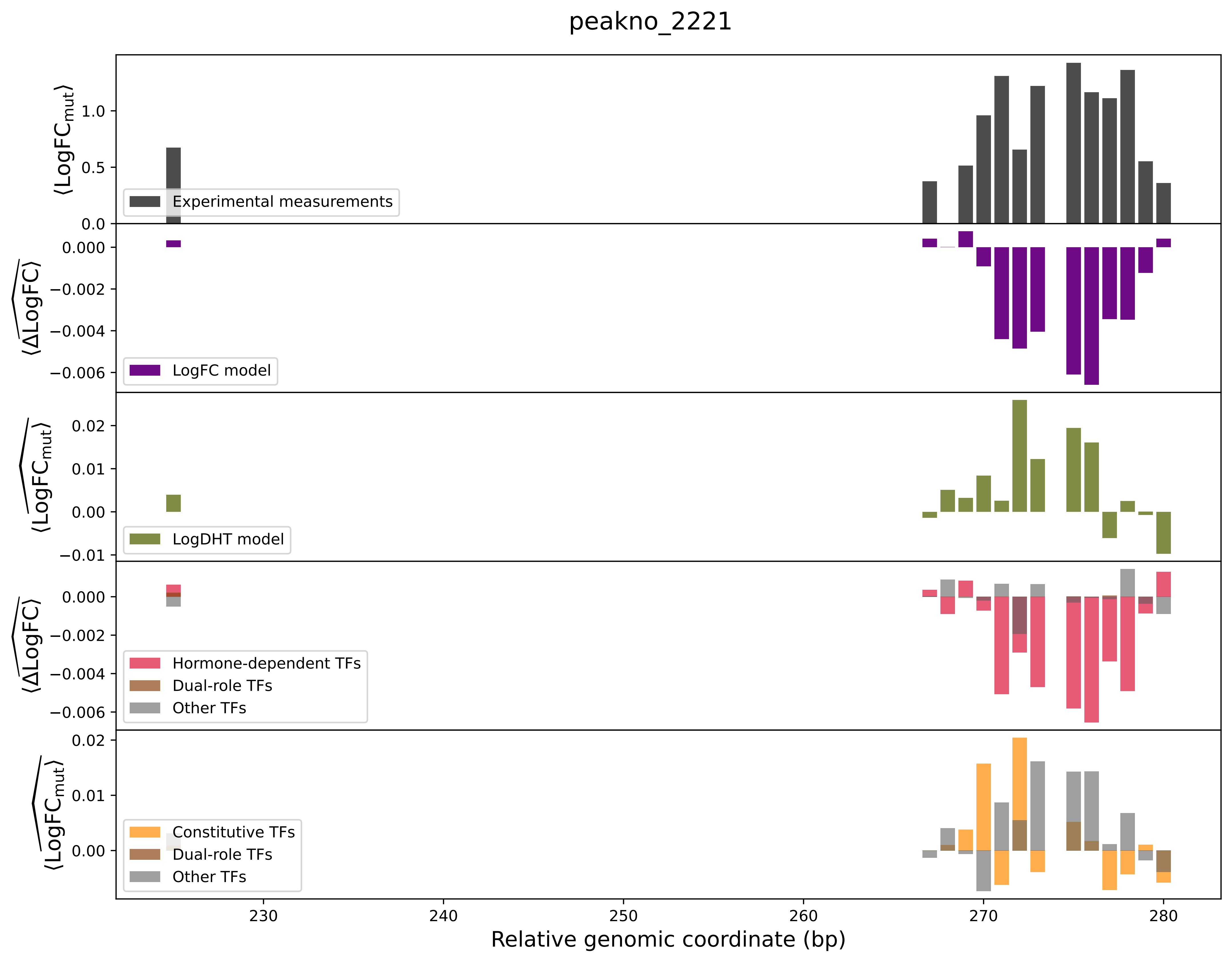

peakno_2221
Experimental measurements
LogFC model
LogDHT model
Hormone-dependent TFs
Dual-role TFs
Other TFs
Constitutive TFs
Dual-role TFs
Other TFs
Relative genomic coordinate (bp)

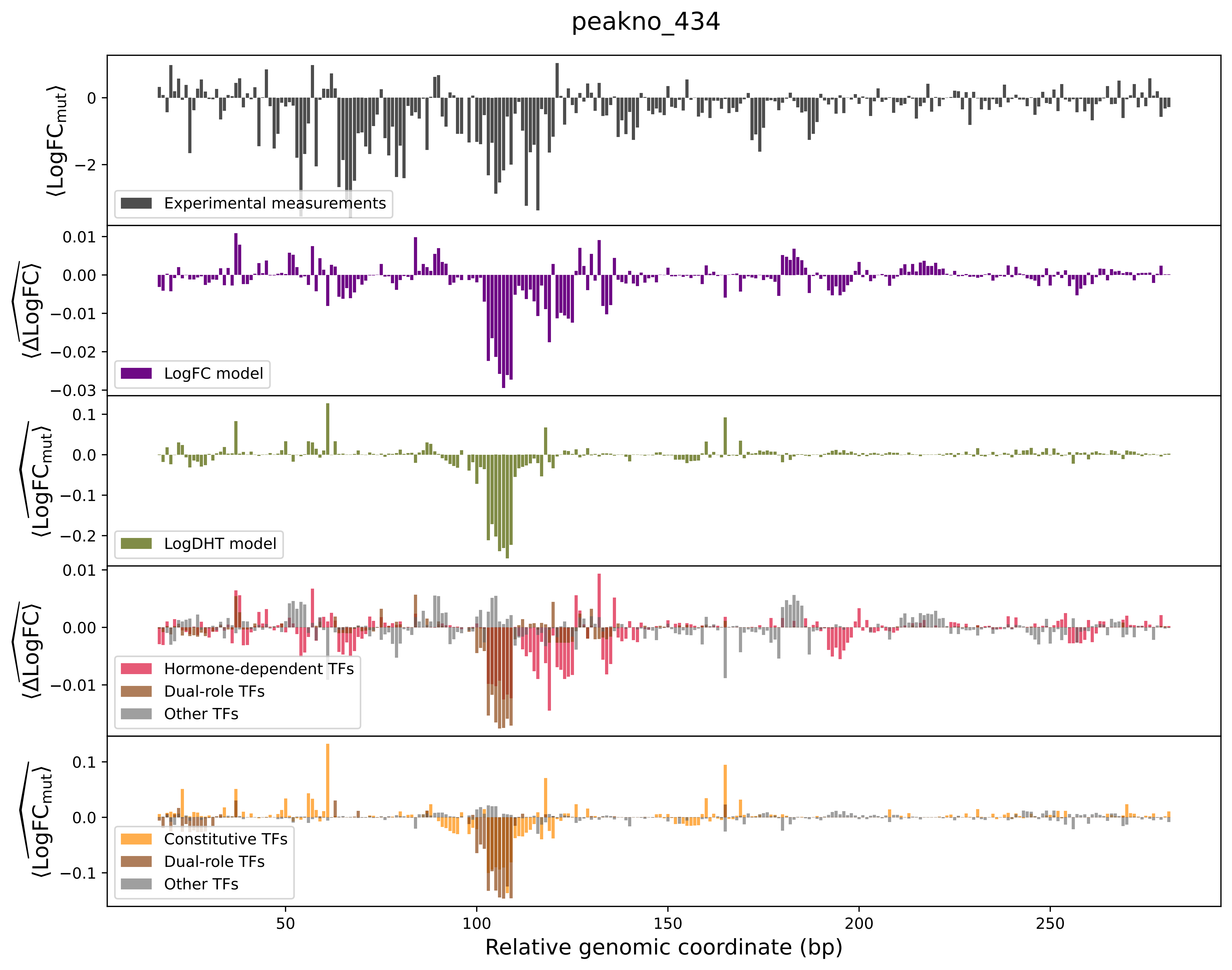
peakno_434
⟨LogFC_mut⟩
Experimental measurements
⟨ΔLogFC⟩
LogFC model
⟨LogFC_mut⟩
LogDHT model
⟨ΔLogFC⟩
Hormone-dependent TFs
Dual-role TFs
Other TFs
⟨LogFC_mut⟩
Constitutive TFs
Dual-role TFs
Other TFs
Relative genomic coordinate (bp)

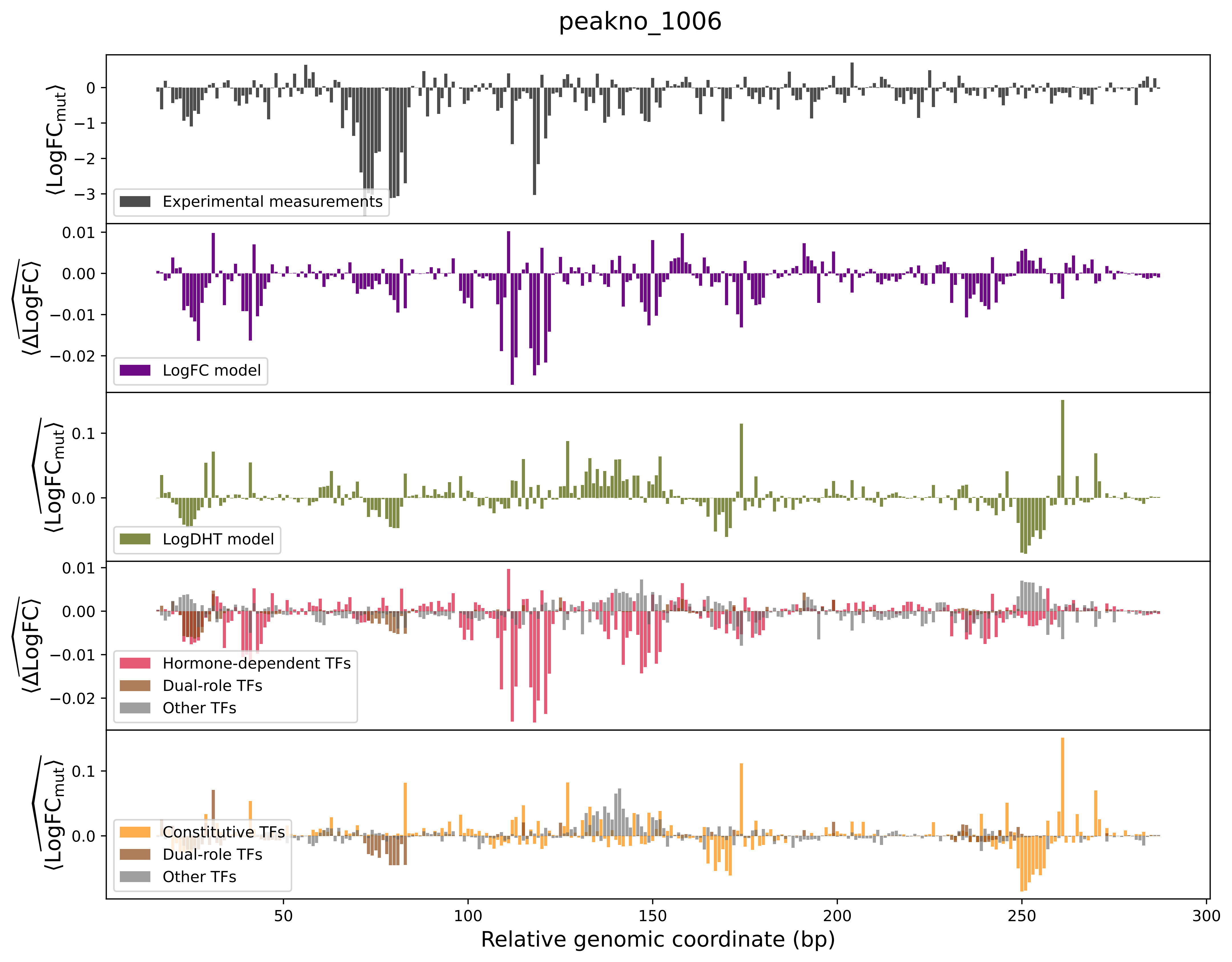

peakno_1006
⟨LogFC_mut⟩
0
−1
−2
−3
Experimental measurements
⟨ΔLogFC⟩
0.01
0.00
−0.01
−0.02
LogFC model
⟨LogFC_mut⟩
0.1
0.0
LogDHT model
⟨ΔLogFC⟩
0.01
0.00
−0.01
−0.02
Hormone-dependent TFs
Dual-role TFs
Other TFs
⟨LogFC_mut⟩
0.1
0.0
Constitutive TFs
Dual-role TFs
Other TFs
50
100
150
200
250
300
Relative genomic coordinate (bp)

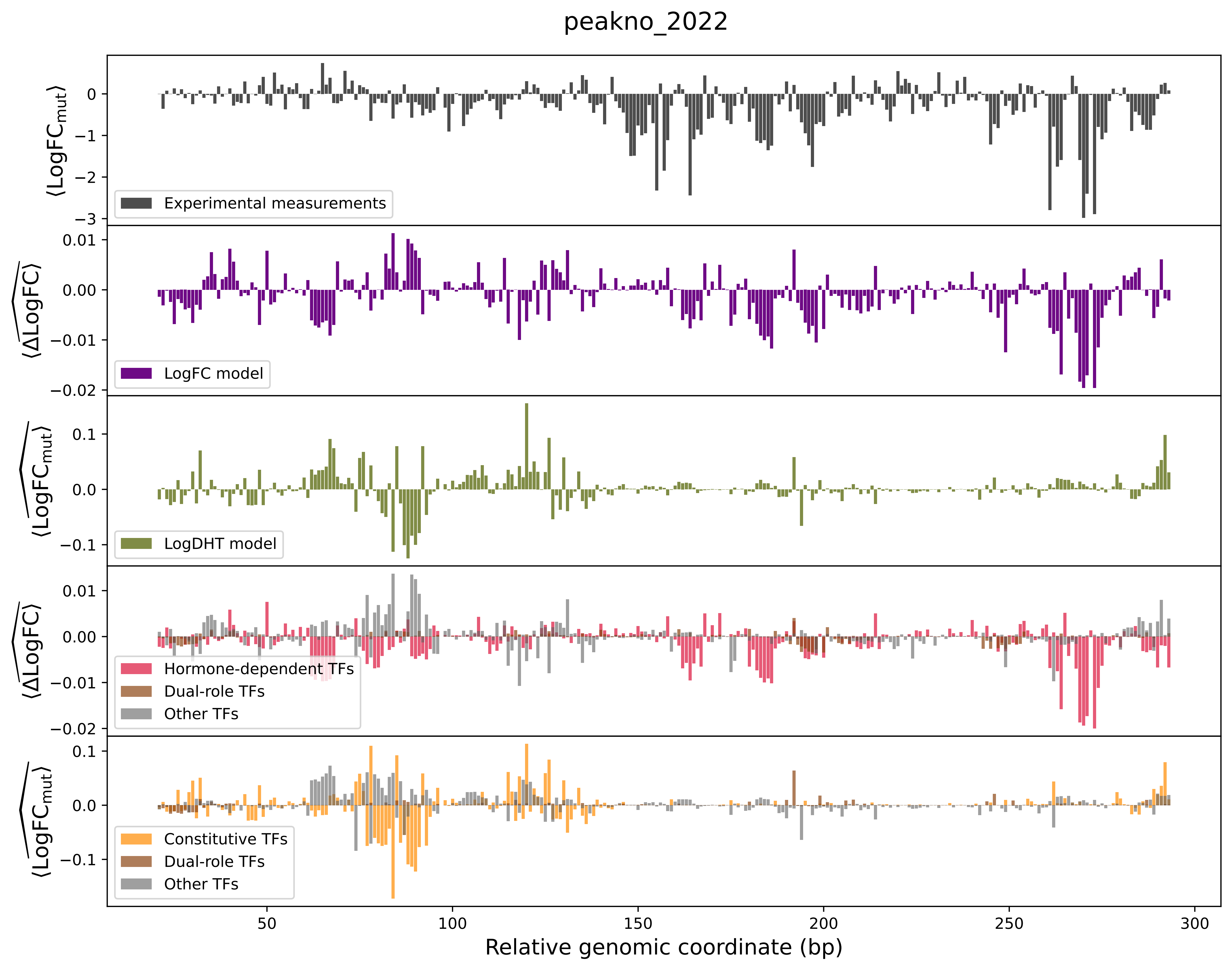
peakno_2022
Experimental measurements
LogFC model
LogDHT model
Hormone-dependent TFs
Dual-role TFs
Other TFs
Constitutive TFs
Dual-role TFs
Other TFs
Relative genomic coordinate (bp)

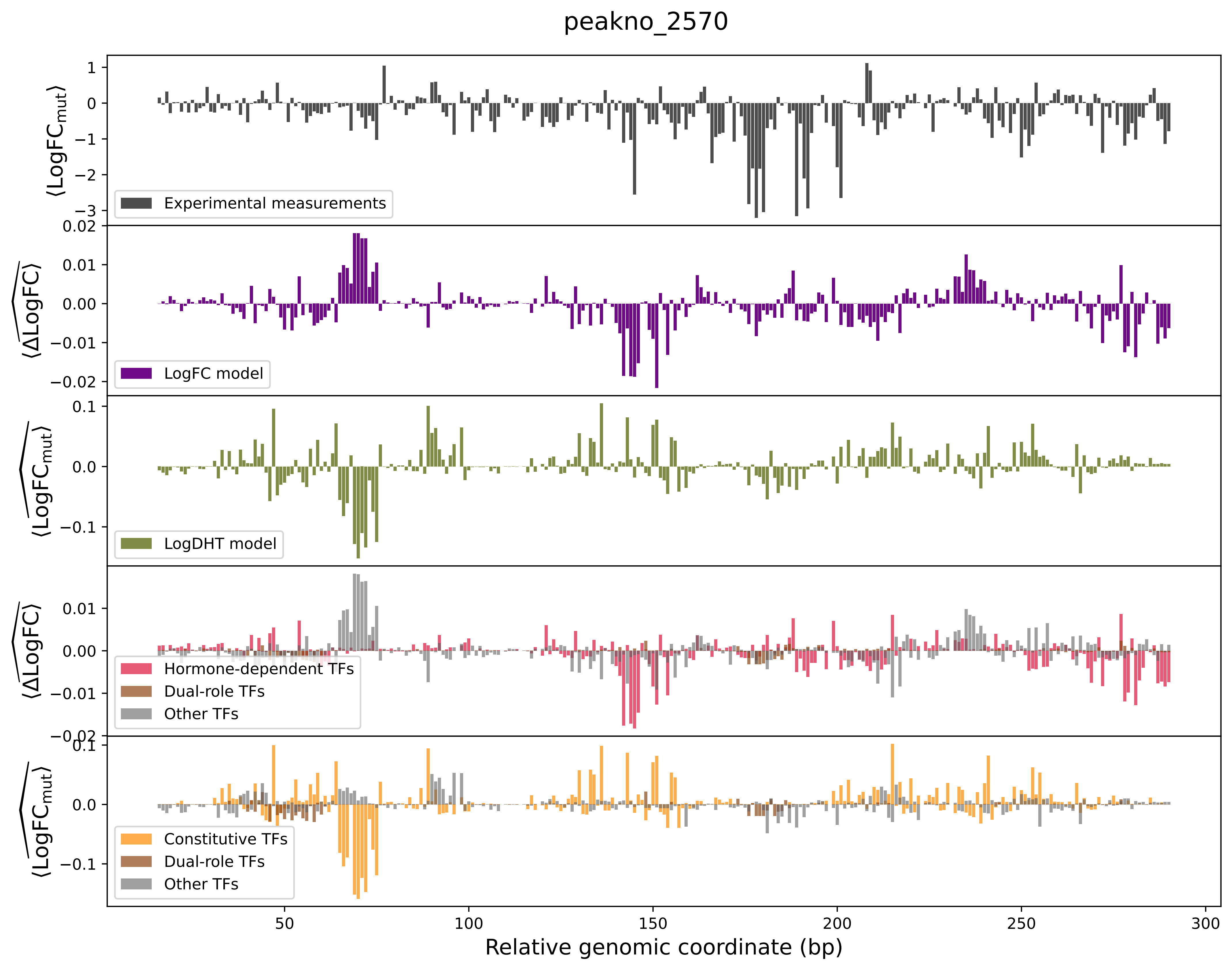

peakno_2570
Experimental measurements
LogFC model
LogDHT model
Hormone-dependent TFs
Dual-role TFs
Other TFs
Constitutive TFs
Dual-role TFs
Other TFs
Relative genomic coordinate (bp)

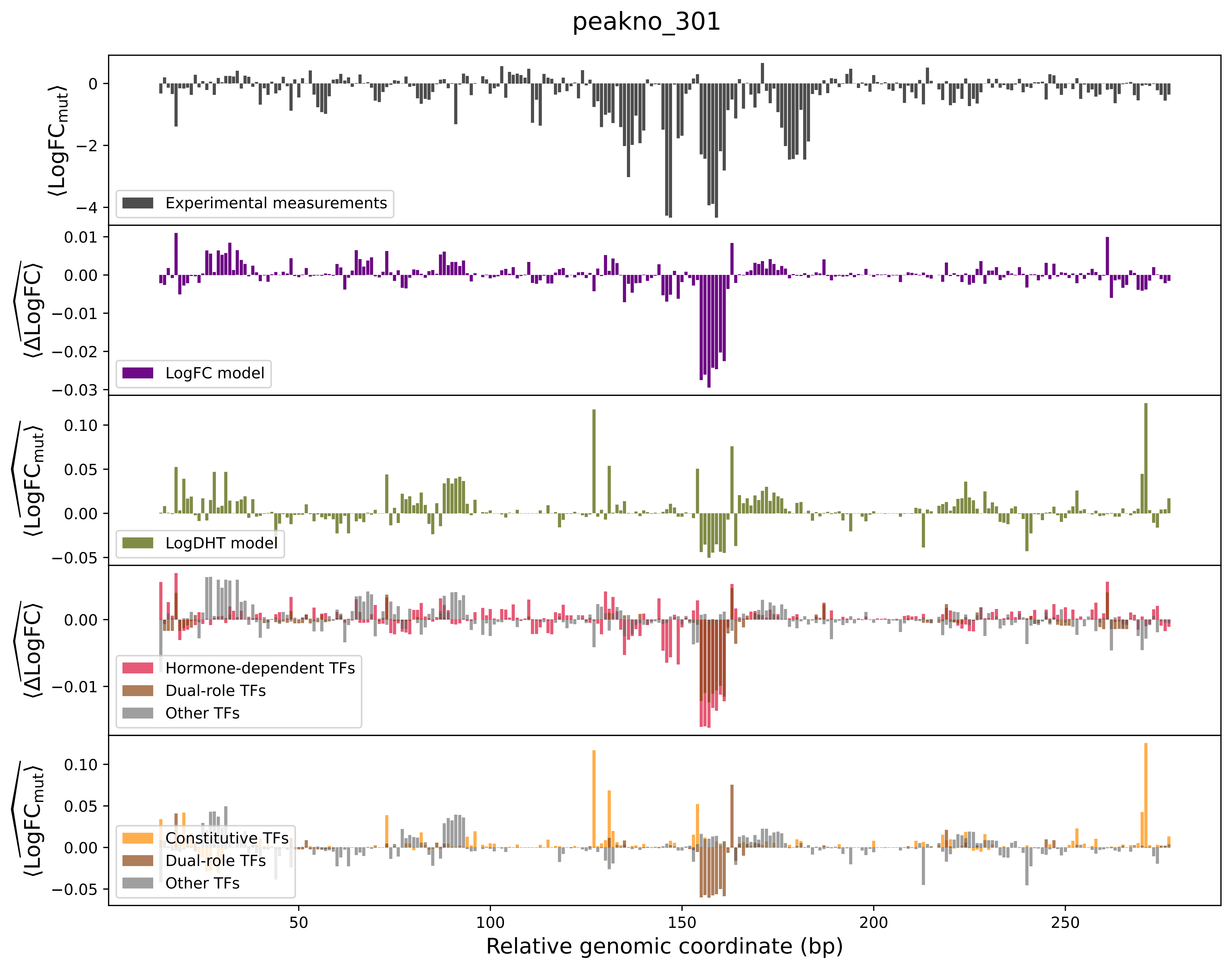

peakno_301
Experimental measurements
LogFC model
LogDHT model
Hormone-dependent TFs
Dual-role TFs
Other TFs
Constitutive TFs
Dual-role TFs
Other TFs
Relative genomic coordinate (bp)

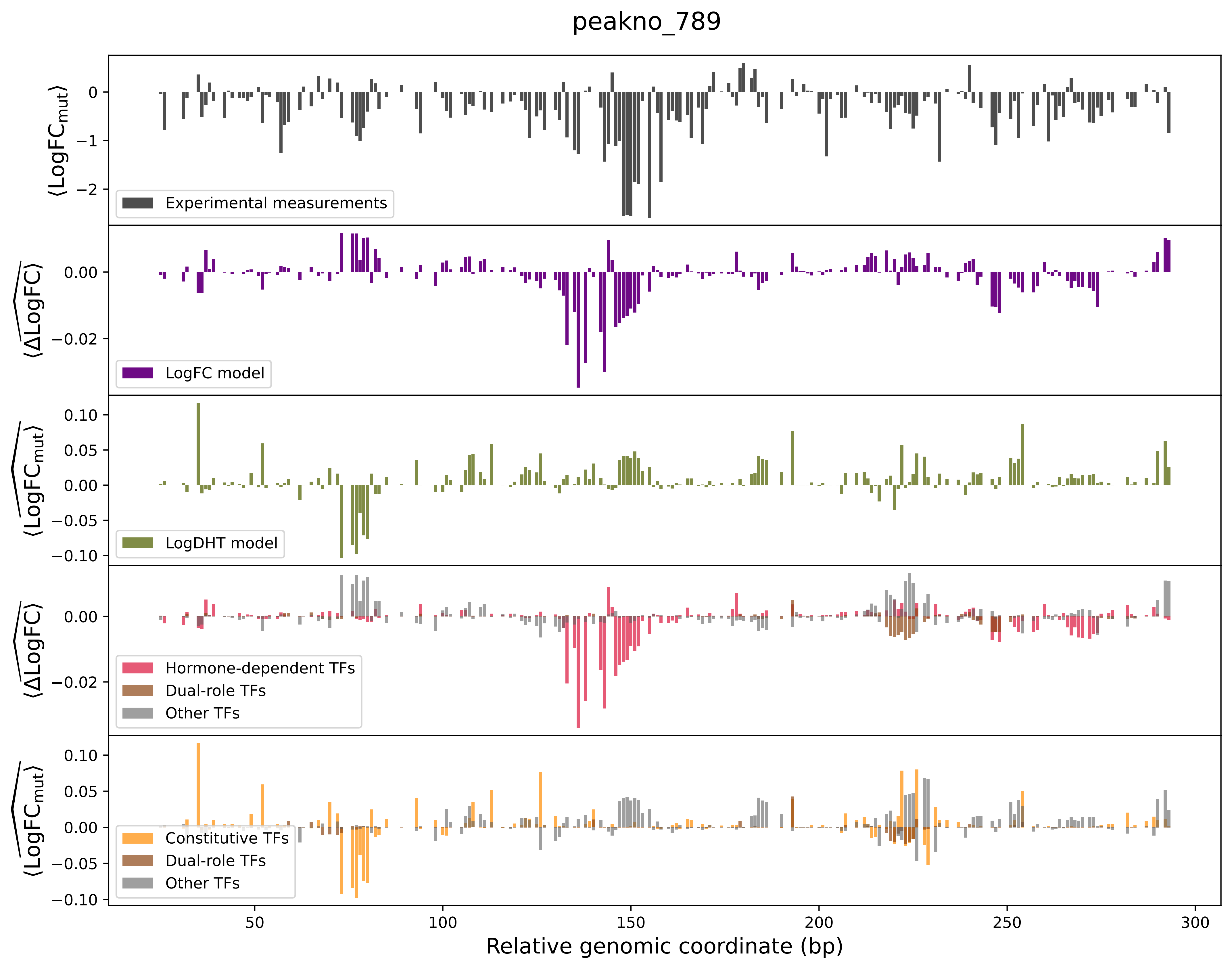

peakno_789
⟨LogFC_mut⟩
Experimental measurements
⟨ΔLogFC⟩
LogFC model
⟨LogFC_mut⟩
LogDHT model
⟨ΔLogFC⟩
Hormone-dependent TFs
Dual-role TFs
Other TFs
⟨LogFC_mut⟩
Constitutive TFs
Dual-role TFs
Other TFs
Relative genomic coordinate (bp)

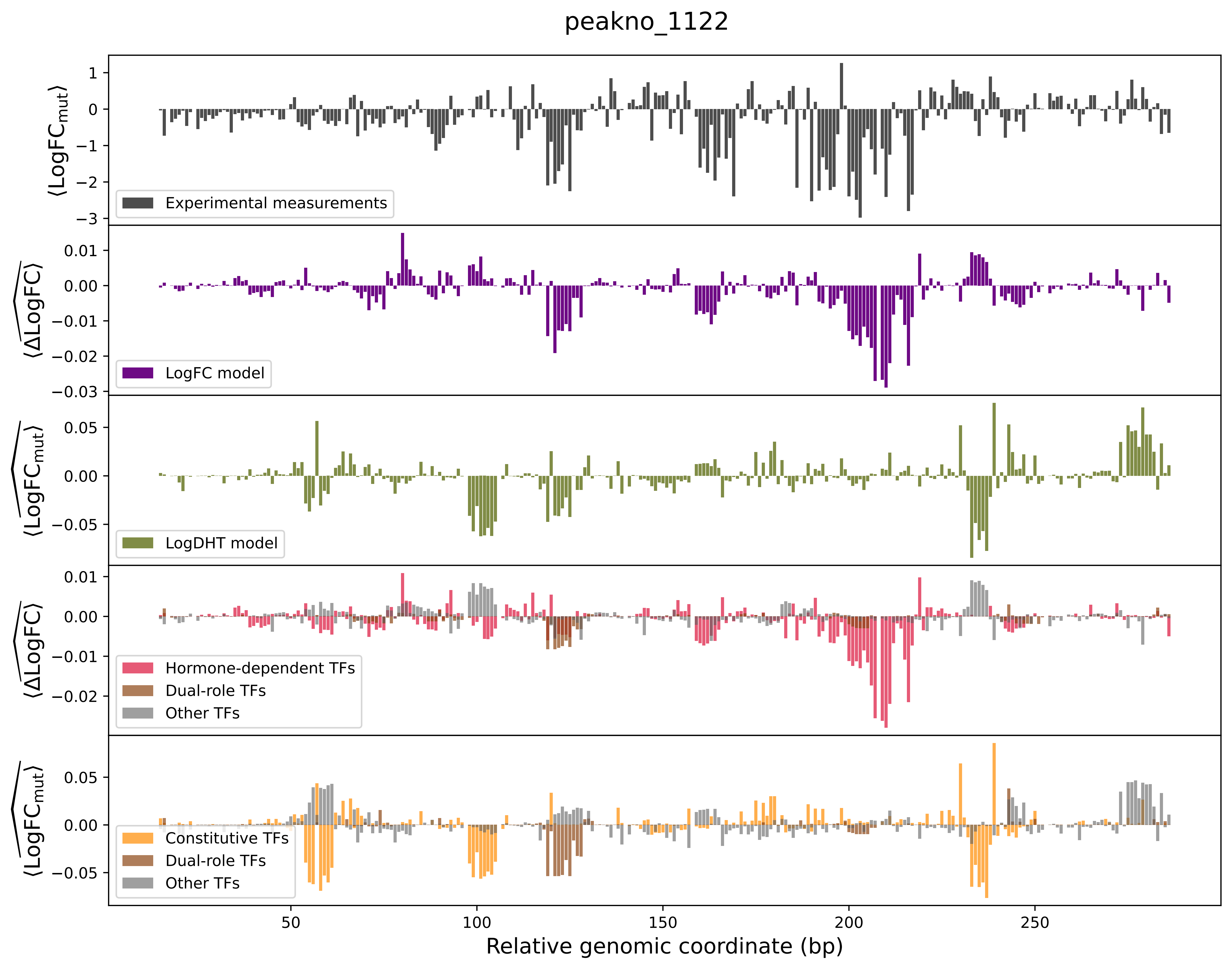

peakno_1122
⟨LogFCmut⟩
Experimental measurements
⟨ΔLogFC⟩
LogFC model
⟨LogFCmut⟩
LogDHT model
⟨ΔLogFC⟩
Hormone-dependent TFs
Dual-role TFs
Other TFs
⟨LogFCmut⟩
Constitutive TFs
Dual-role TFs
Other TFs
Relative genomic coordinate (bp)

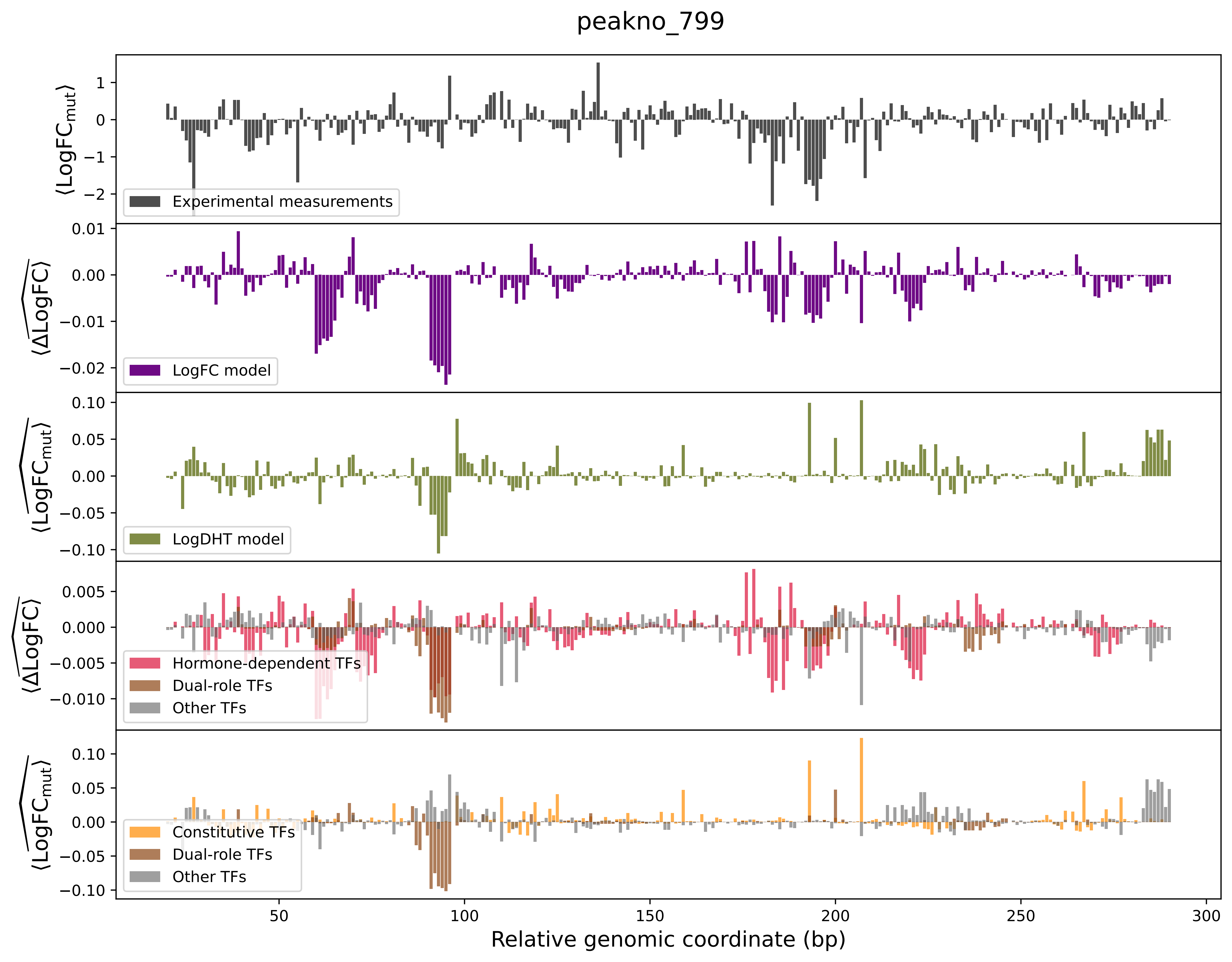

peakno_799
⟨LogFC_mut⟩
Experimental measurements
⟨ΔLogFC⟩
LogFC model
⟨LogFC_mut⟩
LogDHT model
⟨ΔLogFC⟩
Hormone-dependent TFs
Dual-role TFs
Other TFs
⟨LogFC_mut⟩
Constitutive TFs
Dual-role TFs
Other TFs
Relative genomic coordinate (bp)

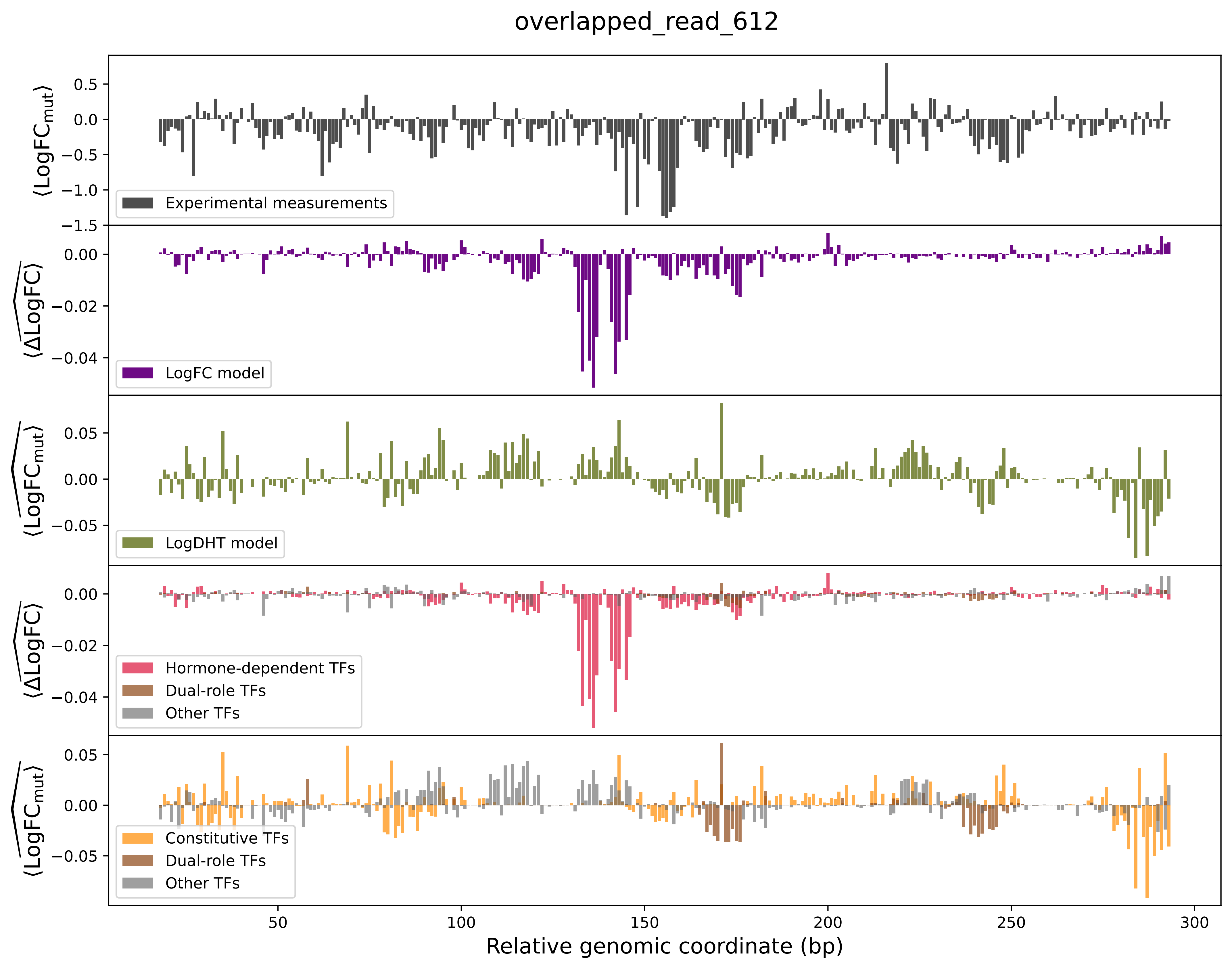

overlapped_read_612
Experimental measurements
LogFC model
LogDHT model
Hormone-dependent TFs
Dual-role TFs
Other TFs
Constitutive TFs
Dual-role TFs
Other TFs
Relative genomic coordinate (bp)

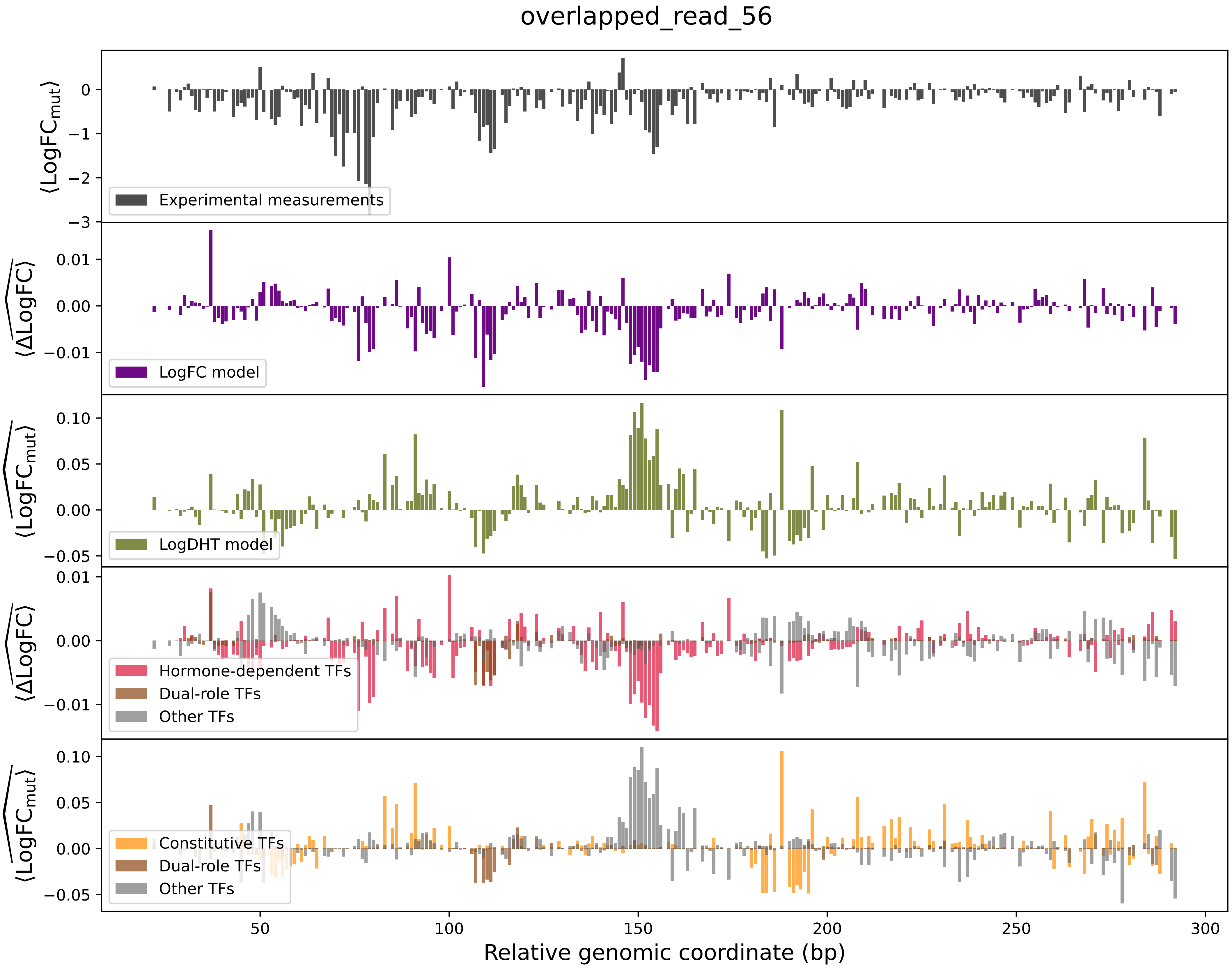
overlapped_read_56
⟨LogFC_mut⟩
Experimental measurements
⟨ΔLogFC⟩
LogFC model
⟨LogFC_mut⟩
LogDHT model
⟨ΔLogFC⟩
Hormone-dependent TFs
Dual-role TFs
Other TFs
⟨LogFC_mut⟩
Constitutive TFs
Dual-role TFs
Other TFs
Relative genomic coordinate (bp)

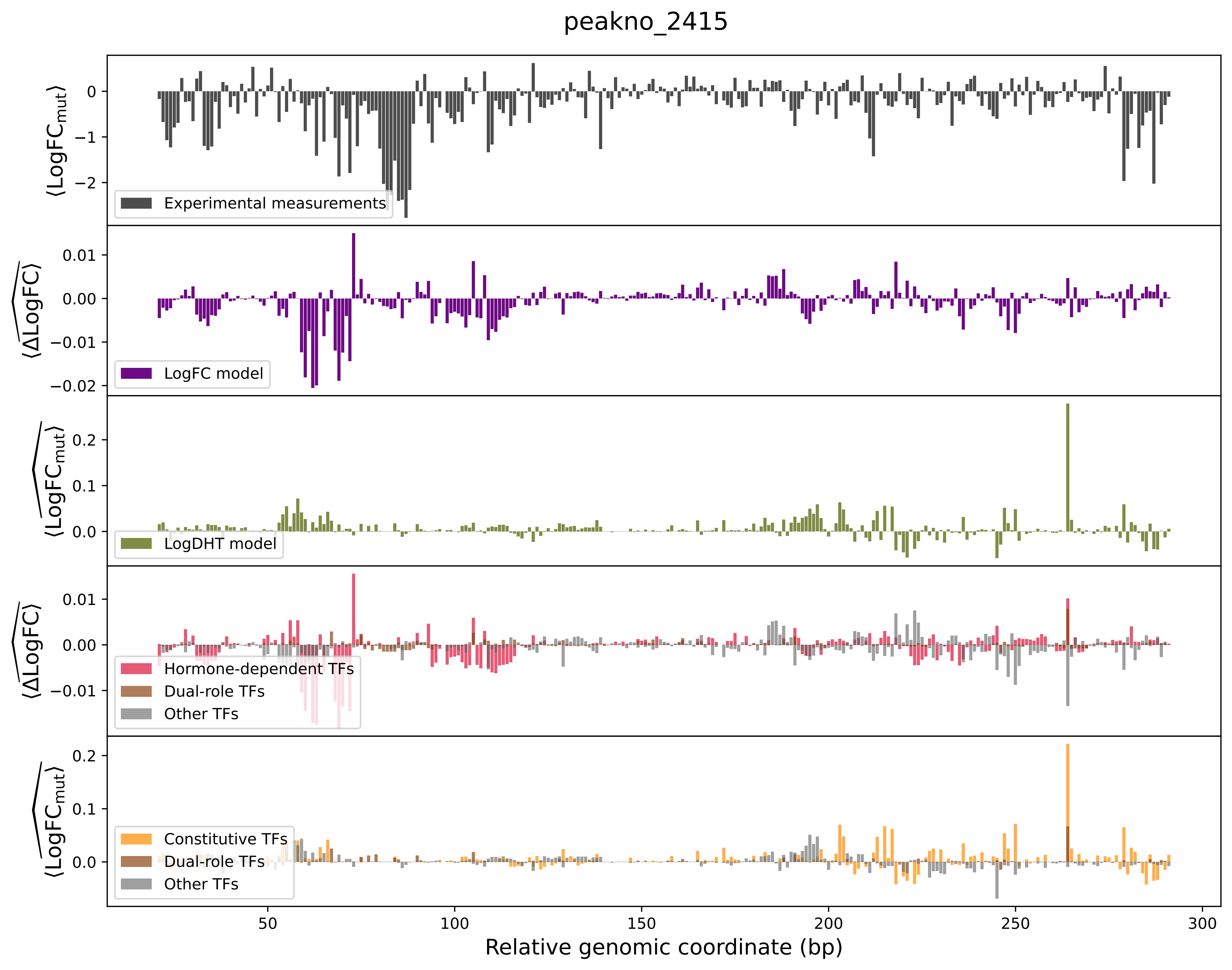

peakno_2415
Experimental measurements
LogFC model
LogDHT model
Hormone-dependent TFs
Dual-role TFs
Other TFs
Constitutive TFs
Dual-role TFs
Other TFs
Relative genomic coordinate (bp)

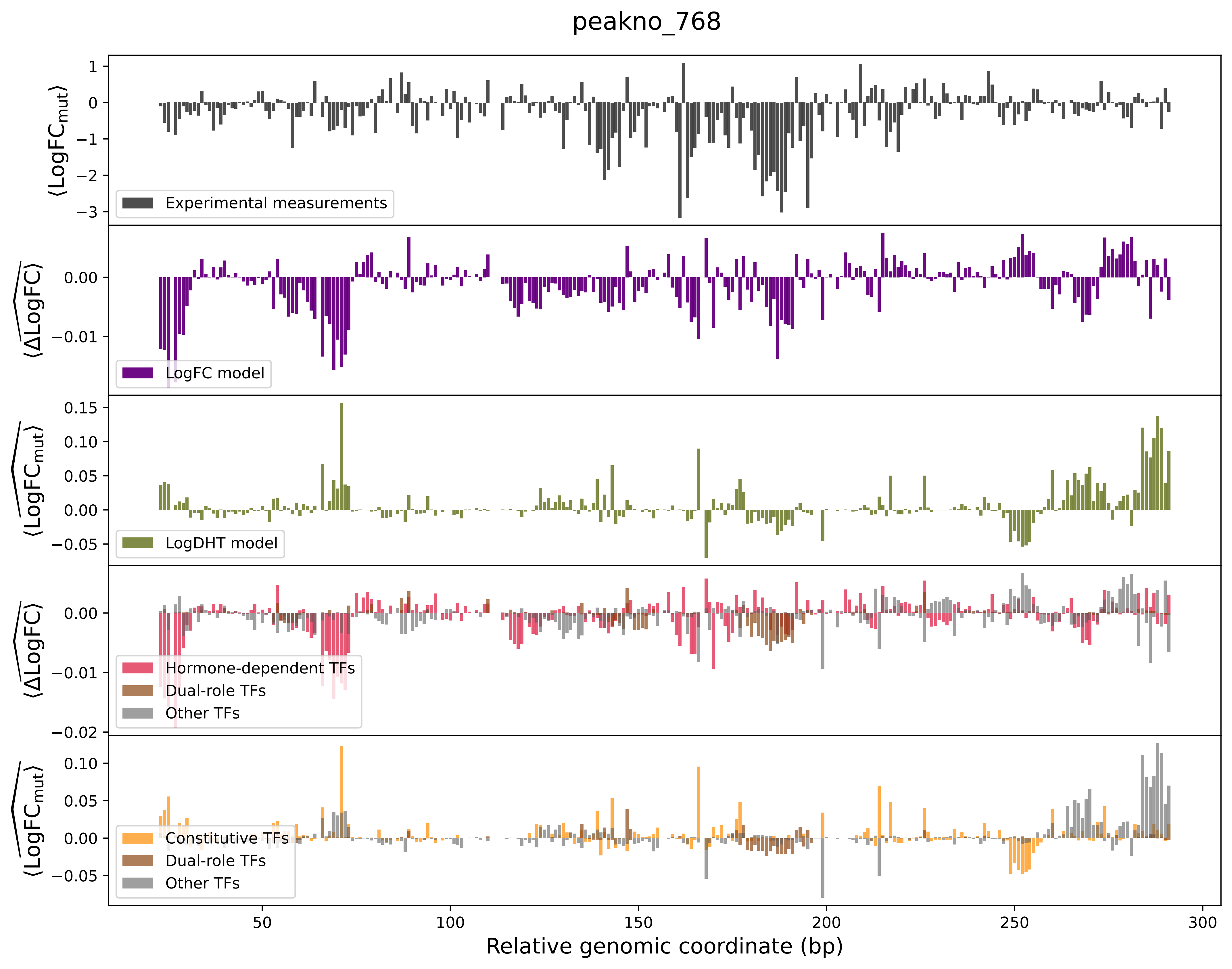

peakno_768
⟨LogFC_mut⟩
Experimental measurements
⟨ΔLogFC⟩
LogFC model
⟨LogFC_mut⟩
LogDHT model
⟨ΔLogFC⟩
Hormone-dependent TFs
Dual-role TFs
Other TFs
⟨LogFC_mut⟩
Constitutive TFs
Dual-role TFs
Other TFs
Relative genomic coordinate (bp)

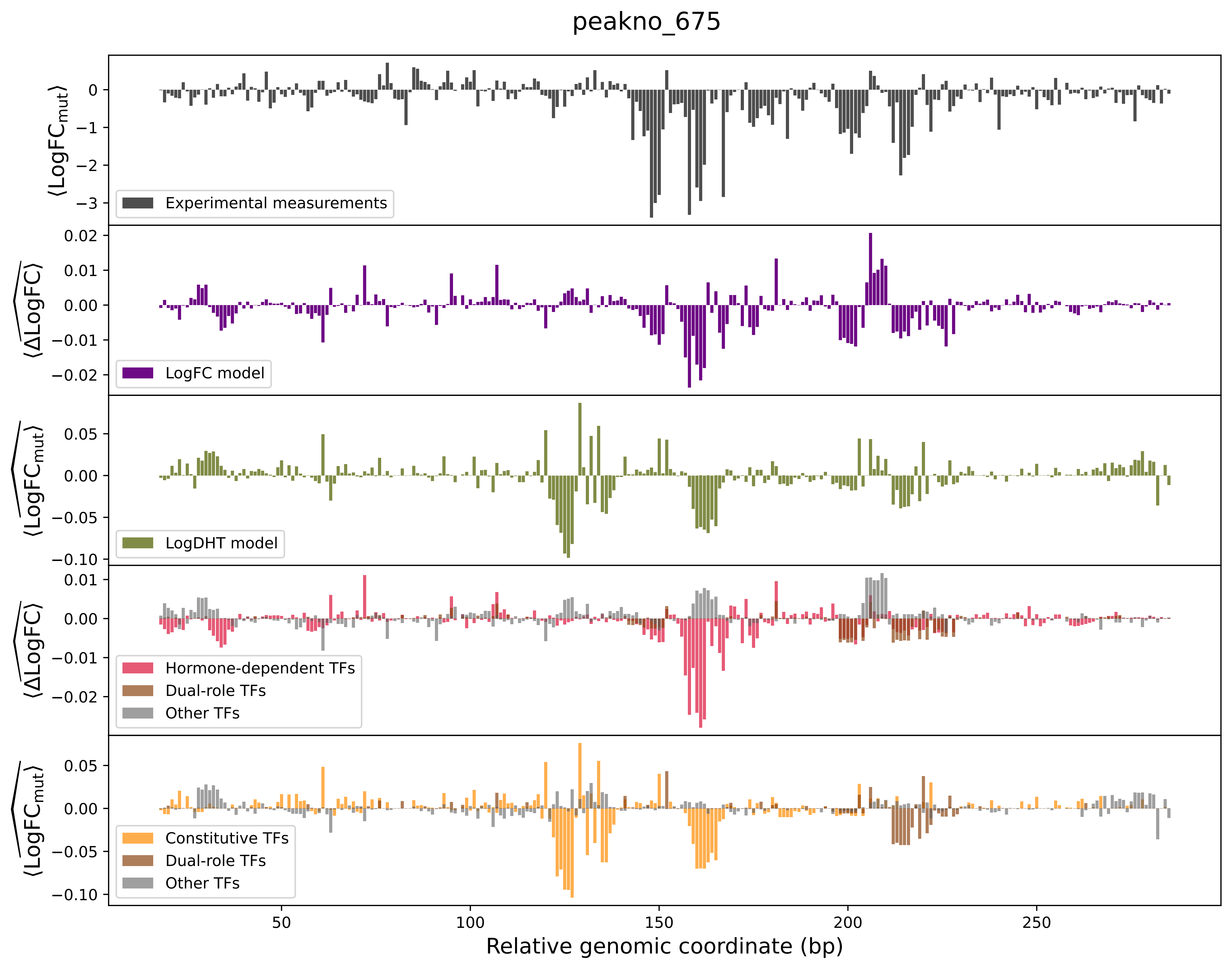

peakno_675
⟨LogFC_mut⟩
Experimental measurements
⟨ΔLogFC⟩
LogFC model
⟨LogFC_mut⟩
LogDHT model
⟨ΔLogFC⟩
Hormone-dependent TFs
Dual-role TFs
Other TFs
⟨LogFC_mut⟩
Constitutive TFs
Dual-role TFs
Other TFs
50
100
150
200
250
Relative genomic coordinate (bp)

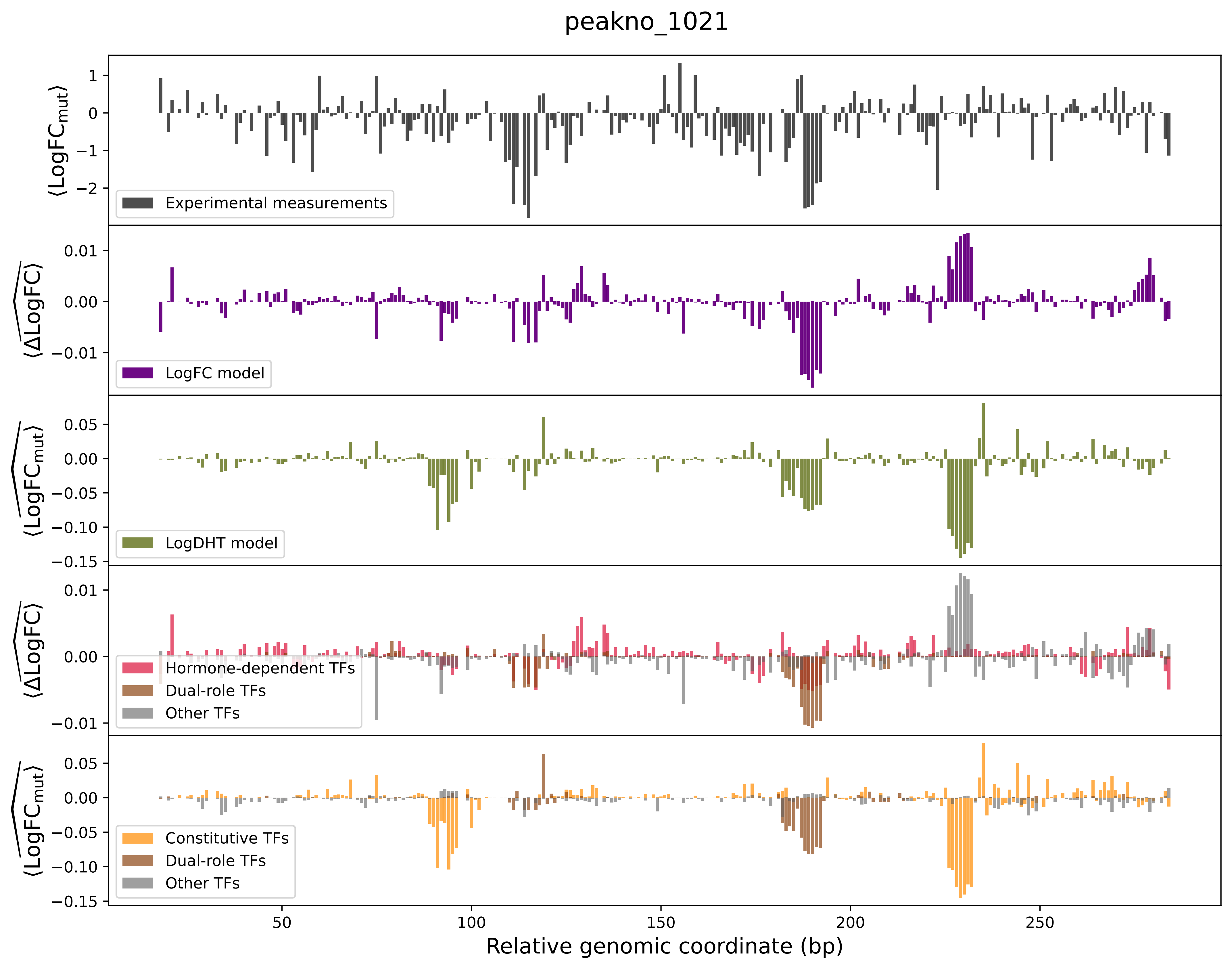
peakno_1021
⟨LogFC_mut⟩
Experimental measurements
⟨ΔLogFC⟩
LogFC model
⟨LogFC_mut⟩
LogDHT model
⟨ΔLogFC⟩
Hormone-dependent TFs
Dual-role TFs
Other TFs
⟨LogFC_mut⟩
Constitutive TFs
Dual-role TFs
Other TFs
Relative genomic coordinate (bp)

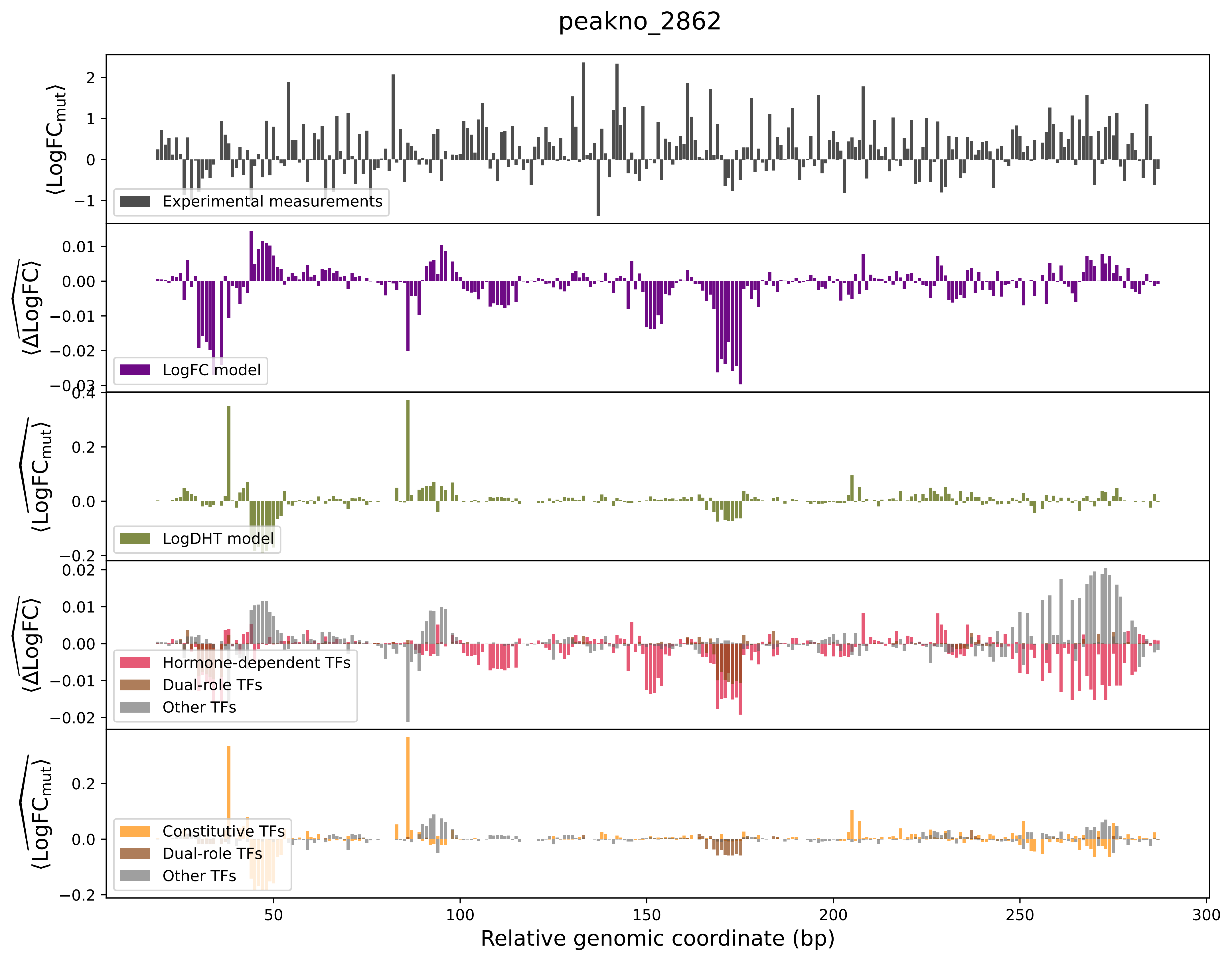

peakno_2862
Experimental measurements
LogFC model
LogDHT model
Hormone-dependent TFs
Dual-role TFs
Other TFs
Constitutive TFs
Dual-role TFs
Other TFs
Relative genomic coordinate (bp)

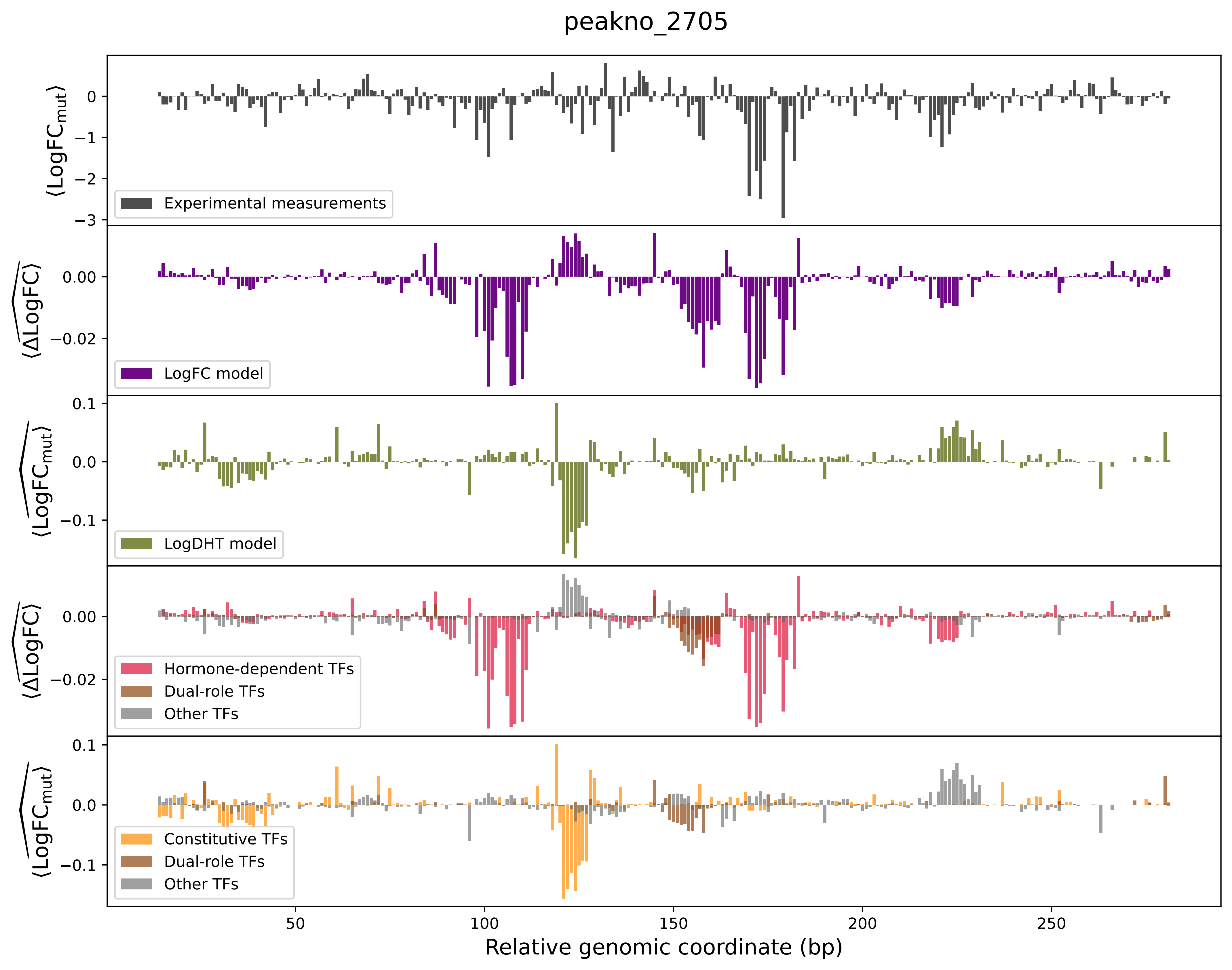
peakno_2705
Experimental measurements
LogFC model
LogDHT model
Hormone-dependent TFs
Dual-role TFs
Other TFs
Constitutive TFs
Dual-role TFs
Other TFs
Relative genomic coordinate (bp)
⟨LogFC_mut⟩
⟨ΔLogFC⟩
⟨LogFC_mut⟩
⟨ΔLogFC⟩
⟨LogFC_mut⟩

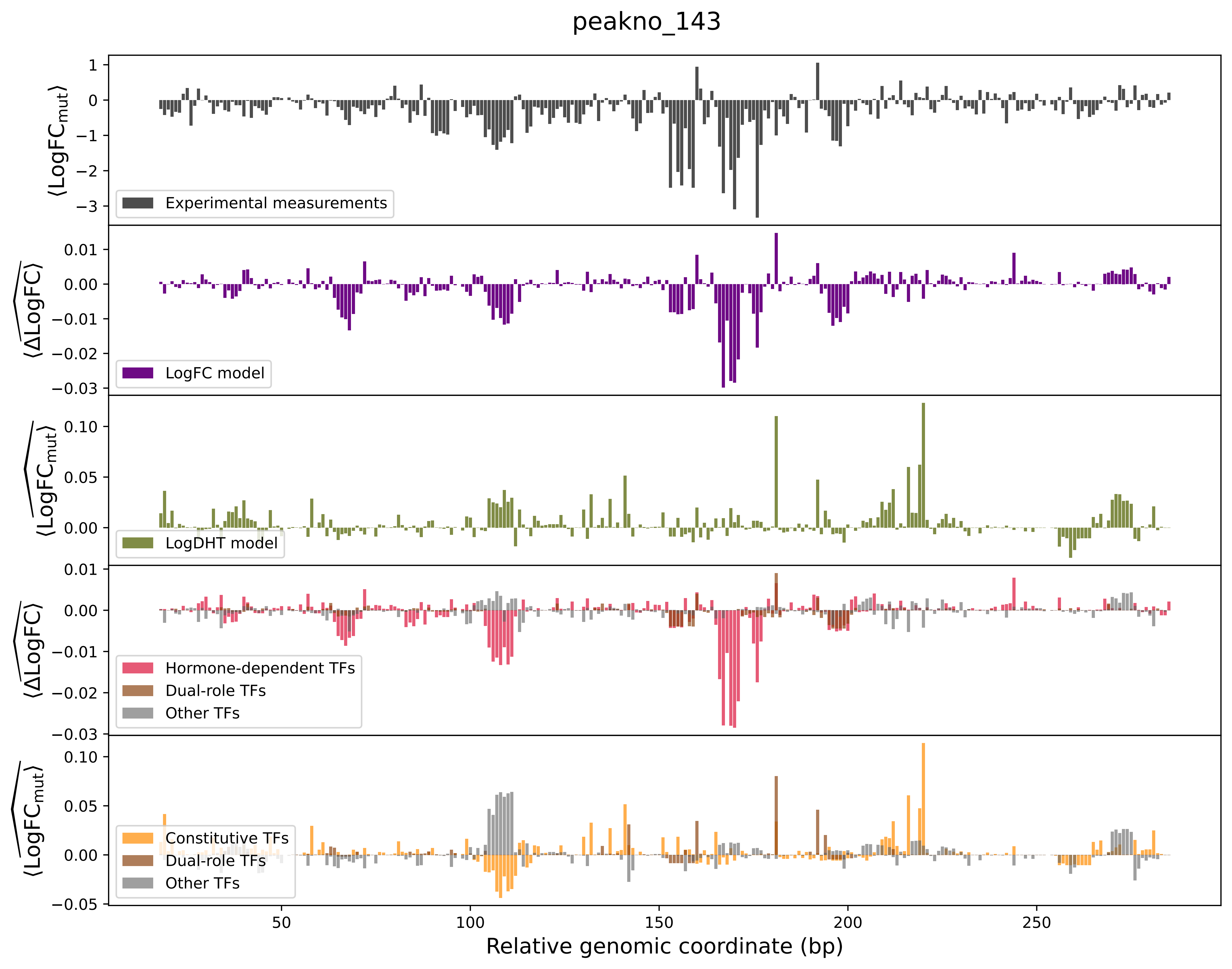

peakno_143
⟨LogFC_mut⟩
Experimental measurements
⟨ΔLogFC⟩
LogFC model
⟨LogFC_mut⟩
LogDHT model
⟨ΔLogFC⟩
Hormone-dependent TFs
Dual-role TFs
Other TFs
⟨LogFC_mut⟩
Constitutive TFs
Dual-role TFs
Other TFs
Relative genomic coordinate (bp)

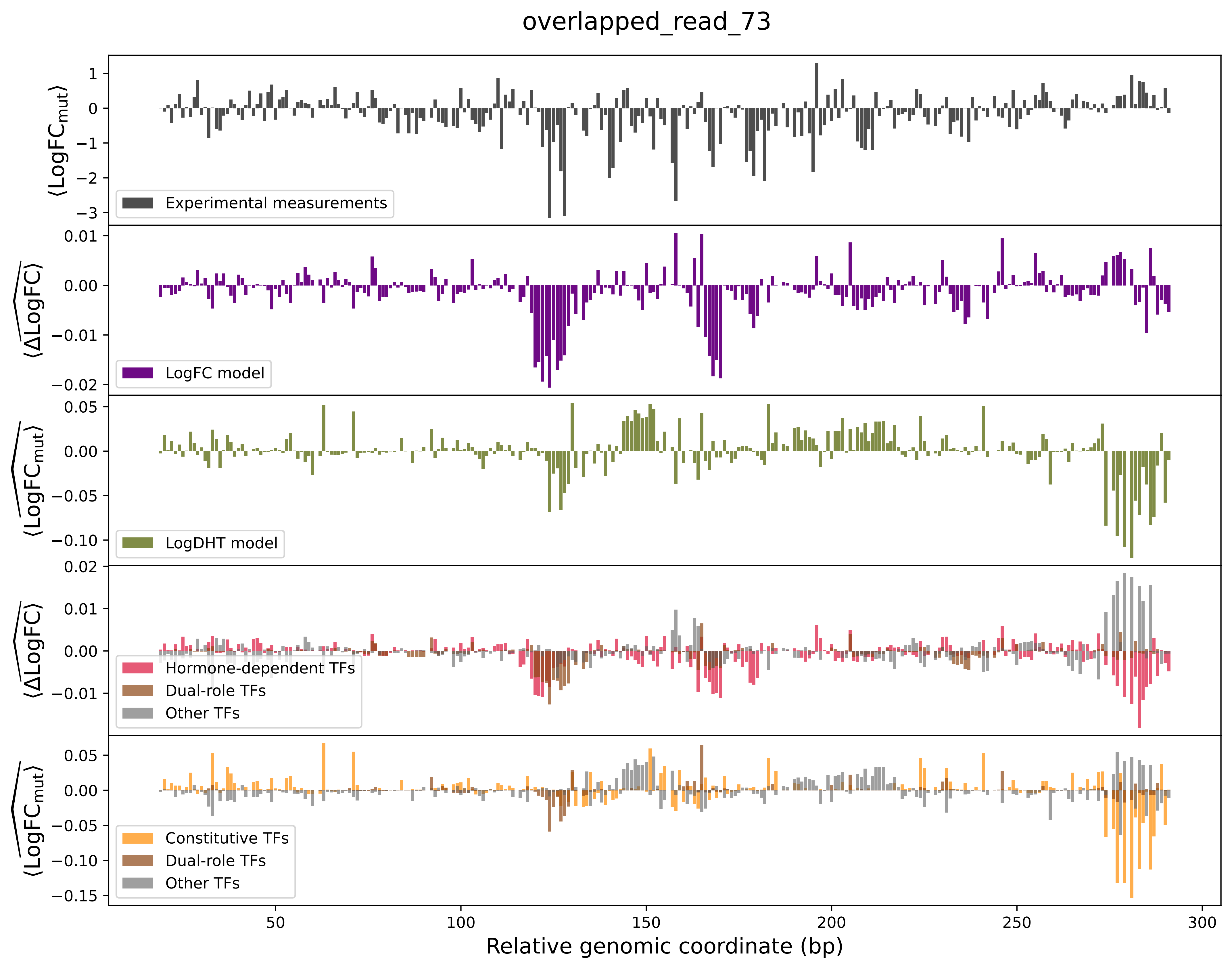

overlapped_read_73
⟨LogFC_mut⟩
Experimental measurements
⟨ΔLogFC⟩
LogFC model
⟨LogFC_mut⟩
LogDHT model
⟨ΔLogFC⟩
Hormone-dependent TFs
Dual-role TFs
Other TFs
⟨LogFC_mut⟩
Constitutive TFs
Dual-role TFs
Other TFs
Relative genomic coordinate (bp)

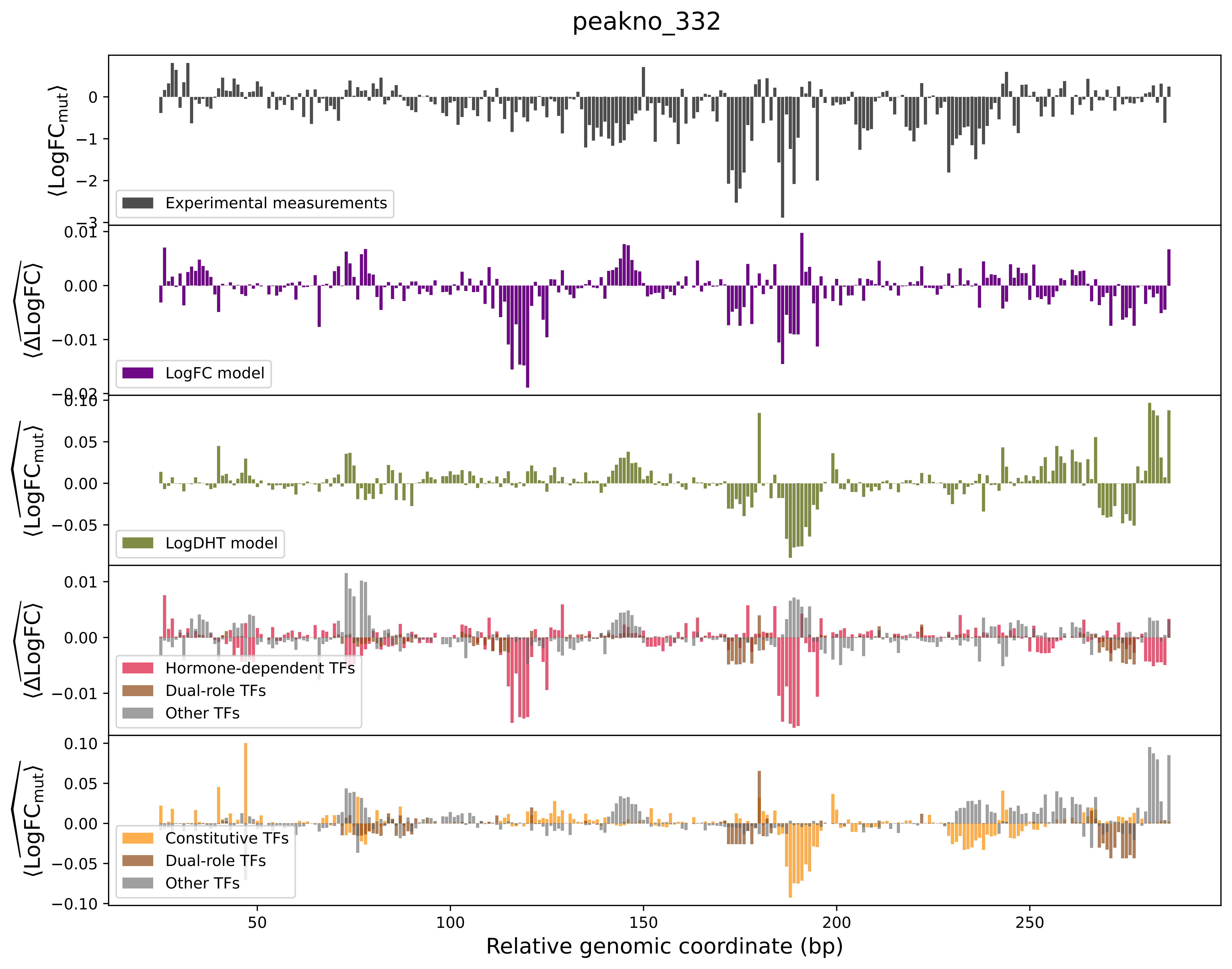

peakno_332
⟨LogFC_mut⟩
Experimental measurements
⟨ΔLogFC⟩
LogFC model
⟨LogFC_mut⟩
LogDHT model
⟨ΔLogFC⟩
Hormone-dependent TFs
Dual-role TFs
Other TFs
⟨LogFC_mut⟩
Constitutive TFs
Dual-role TFs
Other TFs
Relative genomic coordinate (bp)

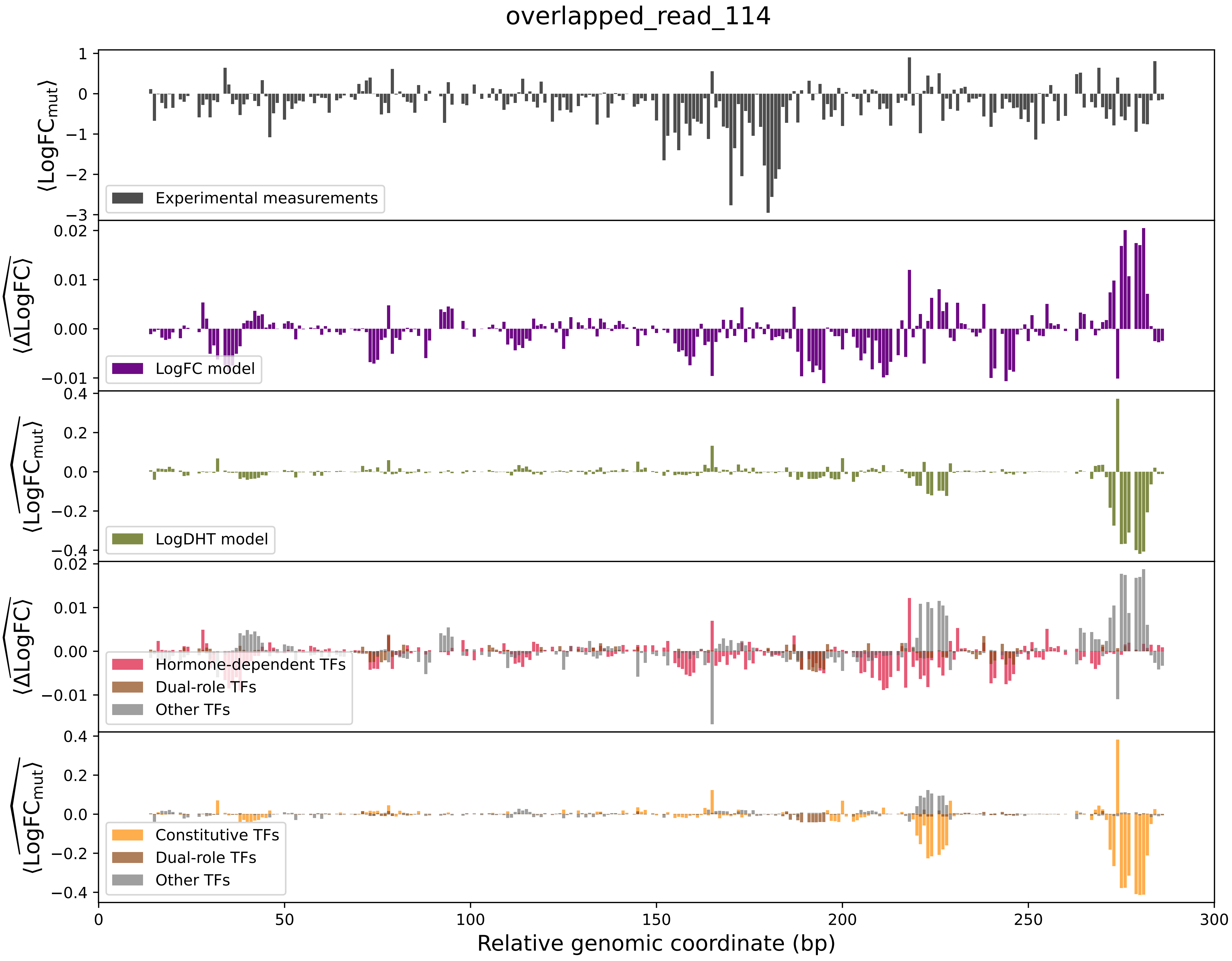

overlapped_read_114
⟨LogFC_mut⟩
Experimental measurements
LogFC model
LogDHT model
Hormone-dependent TFs
Dual-role TFs
Other TFs
Constitutive TFs
Dual-role TFs
Other TFs
Relative genomic coordinate (bp)

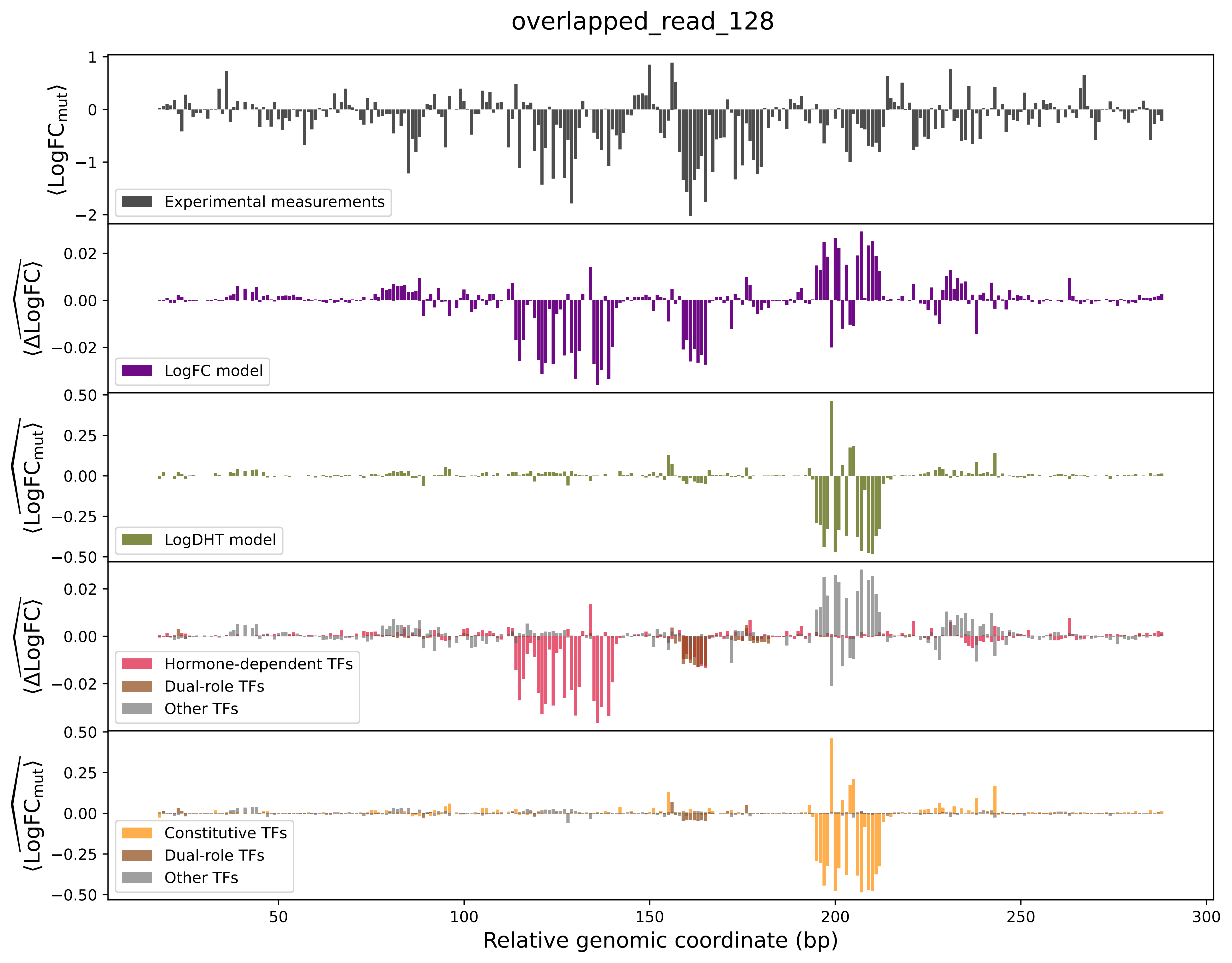

overlapped_read_128
⟨LogFC_mut⟩
Experimental measurements
⟨ΔLogFC⟩
LogFC model
⟨LogFC_mut⟩
LogDHT model
⟨ΔLogFC⟩
Hormone-dependent TFs
Dual-role TFs
Other TFs
⟨LogFC_mut⟩
Constitutive TFs
Dual-role TFs
Other TFs
Relative genomic coordinate (bp)

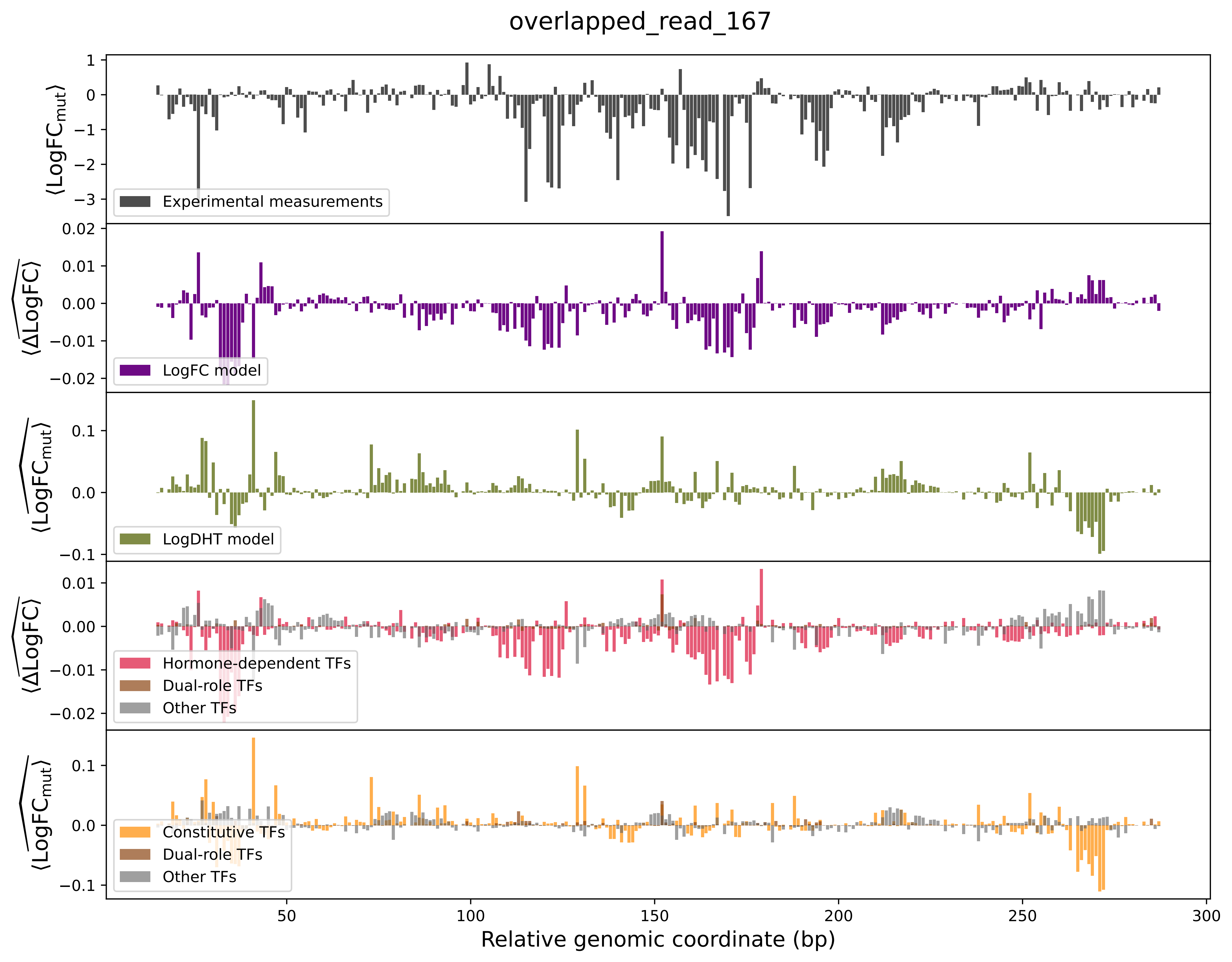

overlapped_read_167
⟨LogFC_mut⟩
Experimental measurements
⟨ΔLogFC⟩
LogFC model
⟨LogFC_mut⟩
LogDHT model
⟨ΔLogFC⟩
Hormone-dependent TFs
Dual-role TFs
Other TFs
⟨LogFC_mut⟩
Constitutive TFs
Dual-role TFs
Other TFs
Relative genomic coordinate (bp)

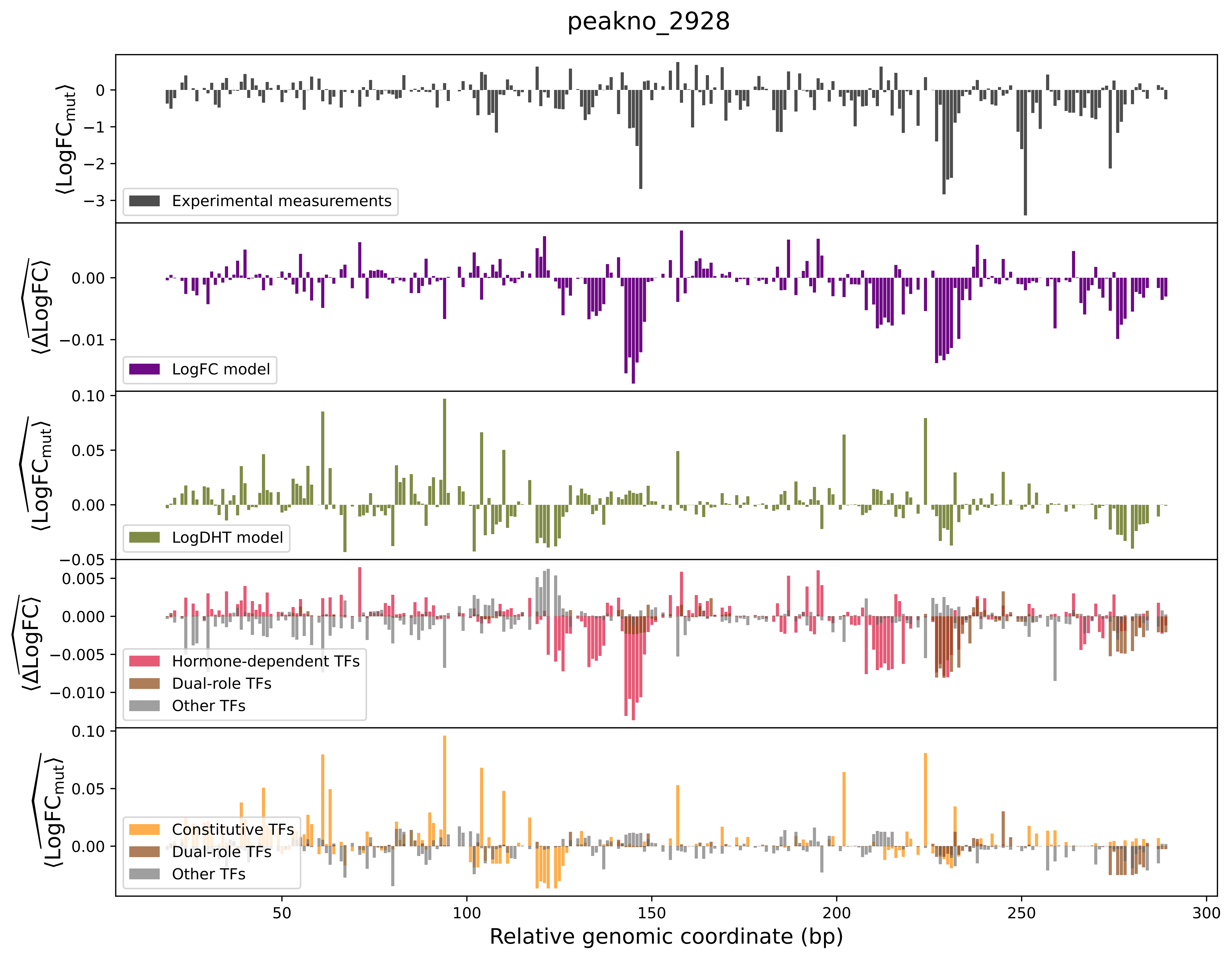

peakno_2928
Experimental measurements
LogFC model
LogDHT model
Hormone-dependent TFs
Dual-role TFs
Other TFs
Constitutive TFs
Dual-role TFs
Other TFs
Relative genomic coordinate (bp)

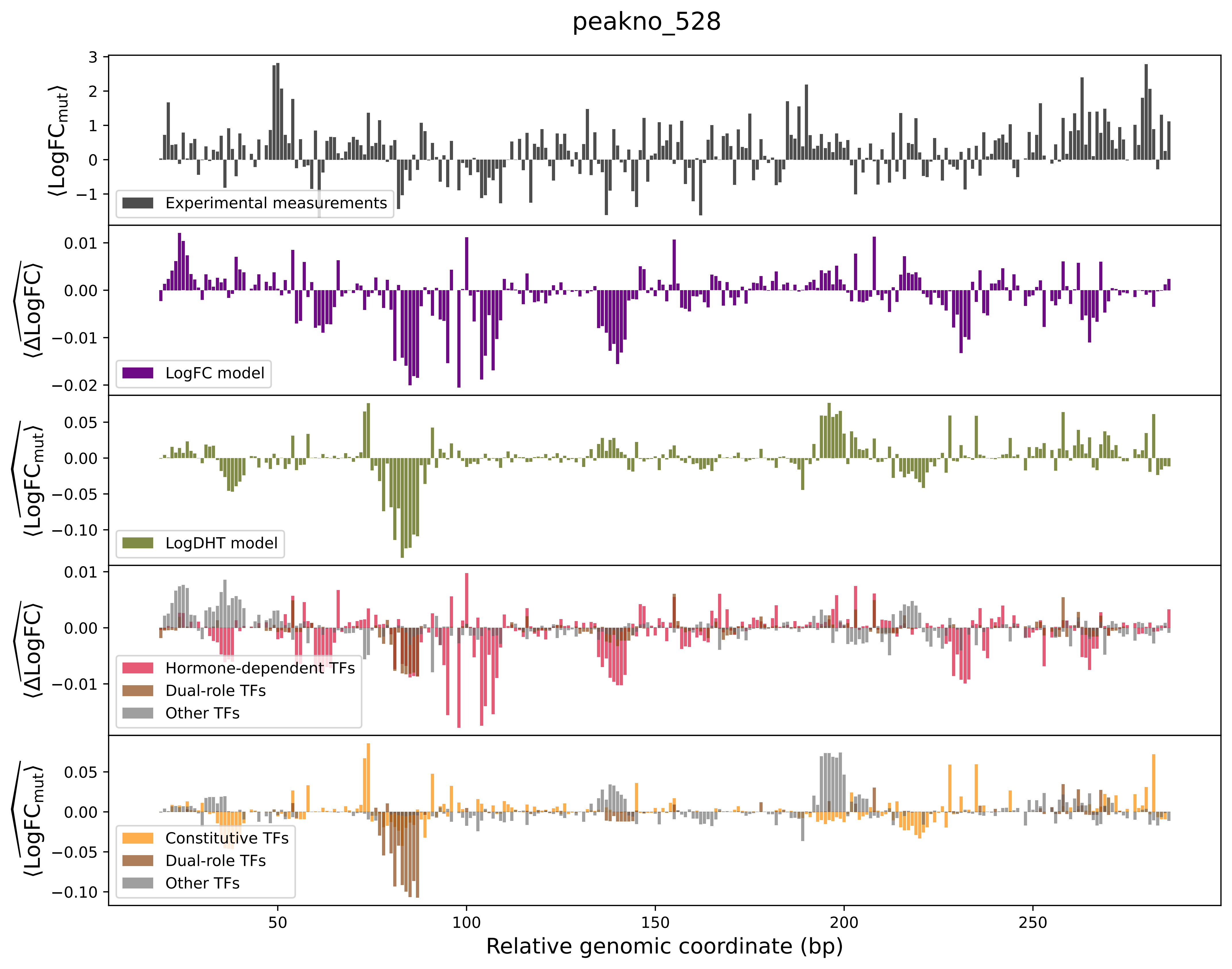
peakno_528
⟨LogFC$_{mut}$⟩
Experimental measurements
⟨$\widehat{\Delta\text{LogFC}}$⟩
LogFC model
⟨$\widehat{\text{LogFC}_{mut}}$⟩
LogDHT model
⟨$\widehat{\Delta\text{LogFC}}$⟩
Hormone-dependent TFs
Dual-role TFs
Other TFs
⟨$\widehat{\text{LogFC}_{mut}}$⟩
Constitutive TFs
Dual-role TFs
Other TFs
Relative genomic coordinate (bp)

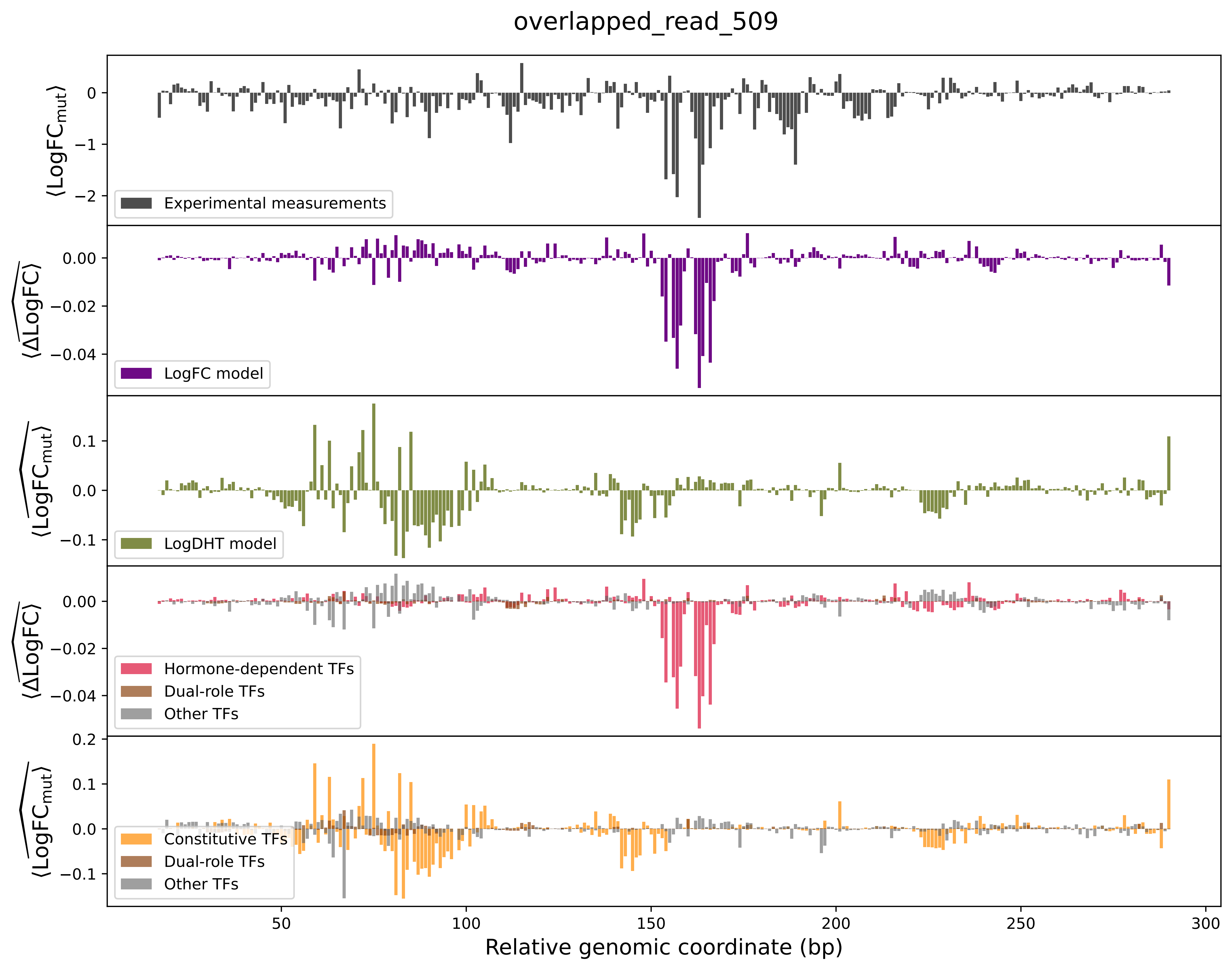

overlapped_read_509
⟨LogFC_mut⟩
Experimental measurements
⟨ΔLogFC⟩
LogFC model
⟨LogFC_mut⟩
LogDHT model
⟨ΔLogFC⟩
Hormone-dependent TFs
Dual-role TFs
Other TFs
⟨LogFC_mut⟩
Constitutive TFs
Dual-role TFs
Other TFs
Relative genomic coordinate (bp)

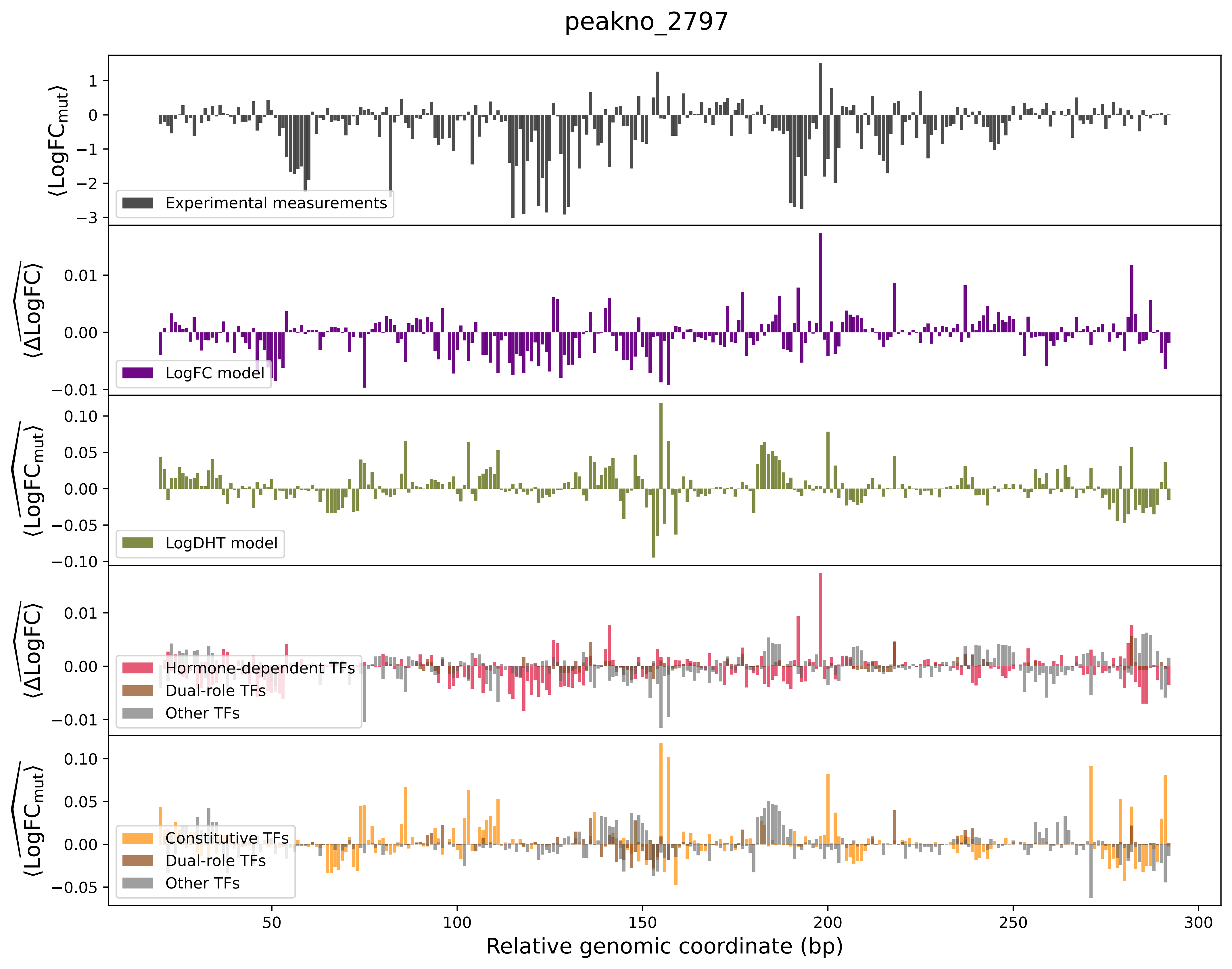
peakno_2797
Experimental measurements
LogFC model
LogDHT model
Hormone-dependent TFs
Dual-role TFs
Other TFs
Constitutive TFs
Dual-role TFs
Other TFs
Relative genomic coordinate (bp)

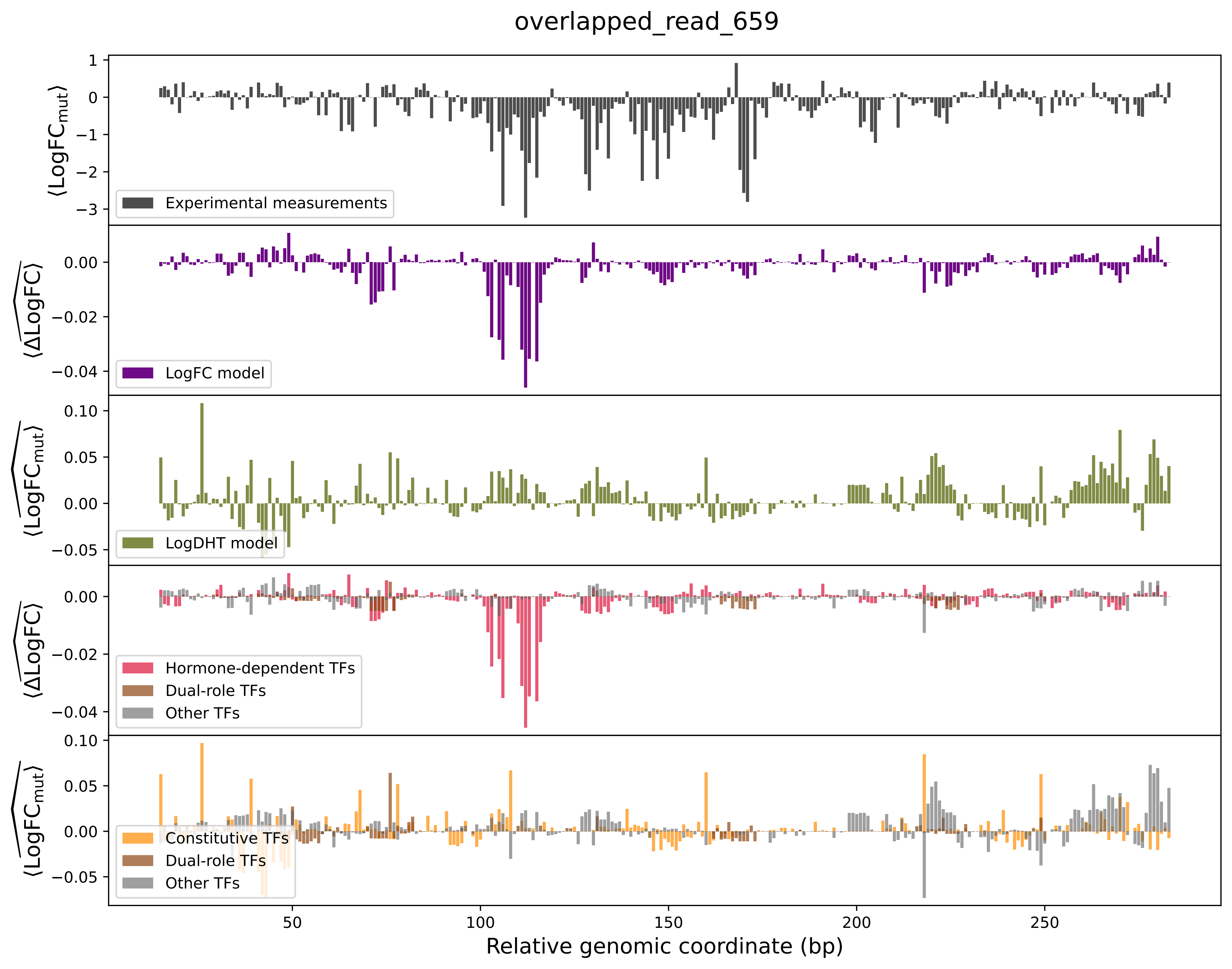
overlapped_read_659
⟨LogFC_mut⟩
Experimental measurements
⟨ΔLogFC⟩
LogFC model
⟨LogFC_mut⟩
LogDHT model
⟨ΔLogFC⟩
Hormone-dependent TFs
Dual-role TFs
Other TFs
⟨LogFC_mut⟩
Constitutive TFs
Dual-role TFs
Other TFs
Relative genomic coordinate (bp)
50
100
150
200
250
